# Porphyrin in prebiotic catalysis: Ascertaining a route for the emergence of early metalloporphyrins

**DOI:** 10.1101/2021.12.20.473587

**Authors:** Shikha Dagar, Susovan Sarkar, Sudha Rajamani

**Affiliations:** Department of Biology, Indian Institute of Science Education and Research, Pune 411008, India

**Keywords:** Molecular evolution, Protometalloenzymes, Porphyrins, Transition metals

## Abstract

Metal ions are known to catalyze certain prebiotic reactions. However, the transition from metal ions to extant metalloenzymes remains unclear. Porphyrins are found ubiquitously in the catalytic core of many ancient metalloenzymes. In this study, we evaluated the influence of porphyrin-based organic scaffold, on the catalysis, emergence and putative molecular evolution of prebiotic metalloporphyrins. We studied the effect of porphyrins on the transition metal ion-mediated oxidation of hydroquinone (HQ). We report a change in the catalytic activity of the metal ions in the presence of porphyrin. This was observed to be facilitated by the coordination between metal ions and porphyrins or by formation of non-coordinated complexes. The metal-porphyrin complexes also oxidized NADH, underscoring its versatility at oxidizing more than one substrate. Our study highlights the selective advantage that some of the metal ions would have had in the presence of porphyrin, underscoring their role in shaping the evolution of protometalloenzymes.

## Introduction

Electron transfer reactions are ubiquitously utilized in contemporary biology to maintain chemical disequilibrium and to produce energy. Such redox reactions are predominantly catalyzed by metalloenzymes, which constitute about one-third of the contemporary enzymes repertoire ^1,2^. The metal ions present in these enzymes are either part of the catalytic core, or the structural scaffold^3–6^. Previous reports have demonstrated metal and mineral-mediated catalysis in prebiotic chemistry ^7–13^, especially in the context of Iron-sulfur World hypothesis^5,14,15^. Iron-sulfur clusters and metal-coordinated porphyrins are potentially few of the earliest catalysts to have emerged on the prebiotic Earth owing to their presence in the catalytic core of some of the most ancient enzymes ^1,16,2^. In this regard, few earlier studies have shown the interaction of such iron-sulfur clusters with peptide chains to form a protoenzyme-like entity^17,18^. In 2018, Bonfio *et. al.* demonstrated the ability of iron-coordinated peptide scaffolds to transfer electrons and generate a pH gradient across membranes^15^. These studies have also tried to address the evolution of this class of protoenzymes from free metal ions, under early Earth conditions. Nonetheless, evolutionary studies in the context of metal-organic scaffolds such as metalloporphyrins are still lacking.

A pioneering study by Melvin Calvin noted a drastic increase in the catalytic efficiency of ferric ions when it was coordinated to a porphyrin scaffold^19,20^. This indicated a central role for metal-bound organic scaffolds such as porphyrins in modulating the catalytic efficiency of metal ions, alluding to the possibility of discerning how evolution of metalloporphyrins could have potentially ensued. Metalloporphyrins are utilized by a wide variety of extant enzymes including magnesium porphyrin in chlorophyll pigments, heme (ferric porphyrin) in hemoglobin, peroxidase and myoglobin, and copper porphyrin in cytochrome c oxidase, to name a few^21,22^. A highly desired property of transition metal ions is their ability to attain multiple oxidation states, which is crucial to catalyze different redox reactions. This “tunability” is employed in biology by coordinating metal ions with the porphyrin rings. Given this, porphyrin has been studied previously as a modular catalyst in biomimetic systems^21,23,24^. Recent studies have reported prebiotically plausible abiotic synthesis of porphyrins, while they have also been detected in the interstellar dust^23,25–31^. Considering their prebiotic plausibility, presence in contemporary biology and the functional diversity of porphyrins, it is reasonable to hypothesize that porphyrins might have played a pivotal role in the emergence and evolution of metalloenzymes on the early Earth.

Given this, we set out to investigate the role of porphyrin scaffolds on the oxidizing ability of metal ions to better understand the emergence of metalloporphyrins on the prebiotic Earth. Towards this, we first evaluated the oxidizing ability of different biologically relevant metal ions namely, magnesium, manganese, iron, cobalt, nickel, copper and zinc. Thereafter, the modulation of the oxidizing ability of the metal ions was studied when porphyrin was added as a co-solute in the reaction. We also investigated the ability of porphyrin to coordinate with these different metal ions under pertinent prebiotic settings. In this regard, the formation kinetics of different metalloporphyrins were studied using steady-state fluorescence and UV spectroscopy. Subsequently, the effect of porphyrin coordination on the oxidizing ability of metal ions was also investigated, to understand the oxidizing ability of preformed metal-coordinated porphyrin complexes.

We observed varying oxidizing ability of different metal ions, with Fe^3+^ showing the maximum oxidation among all the metal ions investigated. Interestingly, in co-solute reactions, significant changes were observed in the oxidizing ability of some metals when compared to the respective free metal ion-based reactions, indicating the influence of porphyrins on the catalytic ability of the metal ions in question. We also report the formation of metal-coordinated porphyrin complexes under prebiotically plausible conditions. The rate of coordination with the porphyrin was found to greatly vary for different metal ions. Also, the oxidizing ability of the preformed metal-coordinated porphyrins showed that the coordination process does not always result in an increase in the oxidizing ability of these metal ions. This is in contrast to a previously reported study wherein the catalytic efficiency was hypothesized to increase upon complexation^19,20^. Specifically, we observed that upon coordination, certain metal ions showed a steep increase in catalysis (in the case of Co^2+^), whereas in other cases, the coordination process dampens the catalytic efficiency of the otherwise catalytic metal ion (in the case of Fe^3+^). Pertinently, we also report an alternate route by which porphyrins would have positively influenced the catalytic efficiency of the metal ions even without coordination. This was found to be achieved by forming non-coordinated assemblies between the metal ion and the porphyrin scaffold. Importantly, metal-coordinated porphyrin complexes could also catalyze the oxidation of nicotinamide adenine dinucleotide hydride (NADH), another biologically relevant molecule in this context. This highlighted their substrate versatility towards catalyzing oxidation reaction. Overall, our study highlights the important role that porphyrin scaffolds would have played in modulating the oxidizing ability of different metal ions. With this study, our attempt is to also set the stage for further exploration of porphyrin-based complexes and, in general, metal organic scaffolds, due to their potential central role in prebiotic catalysis.

## Results

### Experimental section

Hydroquinone (HQ) was used as the primary substrate for the catalytic reaction. This was used as a proxy for ubiquinol (UbQOH), which gets oxidized to ubiquinone^22^ (UbQone, Fig. 1, panel A) by cytochrome in extant biology. NADH oxidation was also evaluated to investigate the promiscuity of the metal-porphyrin complexes.

**Figure 1:**
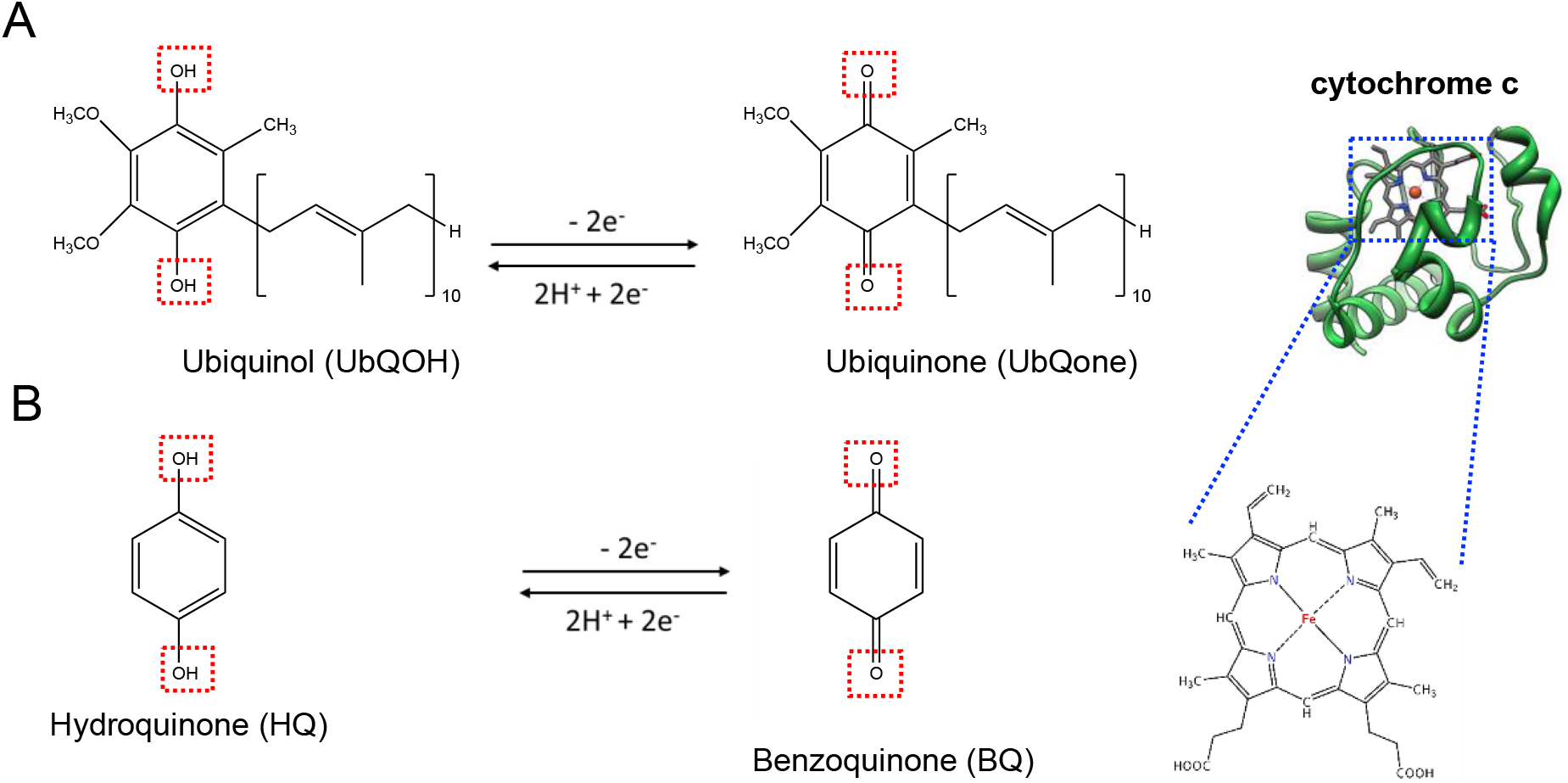
Schematic showing the oxidation of a quinol to its corresponding quinone. A) Oxidation of hydroxyl groups of ubiquinol (UbQOH) to keto groups to form ubiquinone (UbQnone), which involves the loss of 2 electrons and is catalyzed by the iron-porphyrin containing protein cytochrome c (shown in the upper right section; inset shows the iron-porphyrin complex). B) Oxidation of hydroquinone (HQ) to benzoquinone (BQ), a reaction used as a proxy to mimic oxidation of UbQOH to UbQnone. As shown, it also involves the oxidation of 2 hydroxyl groups by loss of 2 electrons, to result in keto groups in BQ.

HQ gets oxidized to benzoquinone (BQ) as is shown in Fig. 1, panel B. This transition from HQ to BQ involves oxidation of the hydroxyl groups to keto groups, with an overall removal of 2 electrons. As a first step, the oxidizing ability of different metal ions on this process was evaluated. A typical reaction mixture contained 0.3 mM HQ with 0.3 mM metal ions (1:1 molar ratio, unless specified otherwise). All the oxidation reactions were performed in the anaerobic glove bag with oxygen levels < 200 ppm. The mixtures were incubated at 40°C and 350 rpm for four hours without any purging with inert gas or stirring (see Methods section). All the reactions were performed under unbuffered conditions. The initial pH was near neutral for most of the reactions (except Fe^3+^ and Cu^2+^ containing reactions) (Supplementary Table 2).

The pH of the reaction mixture was also measured at the end of the reaction (i.e., after four hours), to check for any changes in the pH of the solution (Supplementary Table 2). The extent of oxidation at the initiation of the reaction, and after four hours, was analyzed using High Performance Liquid Chromatography (HPLC) (Supplementary Fig. 1, see Methods). Typically, 30 µl of sample was withdrawn after different time periods, out of which 25 µl was loaded onto the HPLC. Standard curves of HQ (at 288 nm) and BQ (at 244 nm) were utilized to quantify the extent of oxidation by calculating the percentage of BQ produced in the reactions (Supplementary Figs. 5 and 6). The ratio of HQ to metal ions and duration of the reaction, were chosen based on the results of Cu^2+^-mediated oxidation reaction in varying ratios (Supplementary Fig.7). Following this, the metal ion-mediated oxidation was performed in the presence of tetraphenyl porphyrin tetra sulfonic acid (TPPS) as a co-solute. In these co-solute reactions, in addition to 0.3 mM HQ and 0.3 mM metal ion, 0.03 mM TPPS (ten-fold lesser than the HQ and metal ion concentrations) was also added.

Subsequently, the tendency of different metal ions to form metal-TPPS complexes (M-TPPS) under prebiotically relevant conditions, was investigated by incubating 0.3 mM of the respective metal ions with 0.03 mM TPPS at high temperature (70°C). The formation of M-TPPS was monitored using UV and steady-state fluorescence spectroscopy and High-resolution Mass Spectrometry (HRMS). Following this, the oxidizing capability of preformed M-TPPS (metalloporphyrins) was evaluated. Towards this, M-TPPS were prepared by incubating 0.03 mM TPPS with 0.03 mM respective metal ion (1:1) at 70°C, and followed till the completion of coordination. In the case of Fe^3+^-TPPS, commercially acquired reagent was used. These complexes, which were devoid of any free metal ions, were then used to investigate the catalytic ability of preformed M-TPPS. For M-TPPS mediated HQ oxidation, 0.3 mM HQ was incubated with 0.03 mM M-TPPS (ten-fold lesser than HQ), at 40°C. In the case of M-TPPS mediated NADH oxidation reactions, 0.1 mM NADH was incubated at 25°C with 0.01 mM M-TPPS for varying time periods, and the reaction progress was analyzed with UV spectroscopy (see Methods).

### Oxidizing capability of metal ions

Firstly, we compared the oxidizing ability of the different metal ions including Mg^2+^, Mn^2+^, Fe^2+^, Fe^3+^, Co^2+^, Ni^2+^, Cu^2+^ and Zn^2+^ ions. These metal ions were selected based on their presence in the active core of extant metalloporphyrins^33–35^. On the prebiotic Earth, Fe^2+^ would have readily photo-oxidized to Fe^3+^ in the presence of UV and under aqueous conditions, irrespective of the presence of oxygen^36^. Therefore, owing to the susceptibility of Fe^2+^ towards oxidation, and the prebiotic relevance of Fe^3+^, we evaluated the oxidizing ability of iron in its +2 as well as +3 states.

No significant oxidation of HQ was observed in the control reaction, i.e., in the absence of any metal ions (Supplementary Fig.7). In presence of different metals, excepting for Fe^3+^, no oxidation was observed at the initiation of the reaction. For Mg^2+^, Co^2+^, Ni^2+^ and Zn^2+^ ions, no significant oxidation of HQ was observed even after four hours (Fig. 2, Supplementary Fig. 7 and Table 7). In the presence of Mn^2+^, Fe^2+^ and Cu^2+^ ions, 4%, 4.7% and 7% of HQ oxidation, respectively, was observed after four hours. Interestingly, in Fe^3+^ containing reactions, up to 30% BQ was produced immediately upon the addition of the Fe^3+^ ions to the solution. The extent of oxidation by Fe^3+^ did not change significantly after four hours. Overall, the extent of oxidation for the different metal ions was in the following order: Fe^3+^> Cu^2+^> Fe^2+^> Mn^2+^. Interestingly, this trend of HQ oxidation remained unchanged even upon doubling the metal ion concentration (HQ: metal ion = 1:2; Fig. 2, Supplementary Figs. 8 and 9).

**Figure 2:**
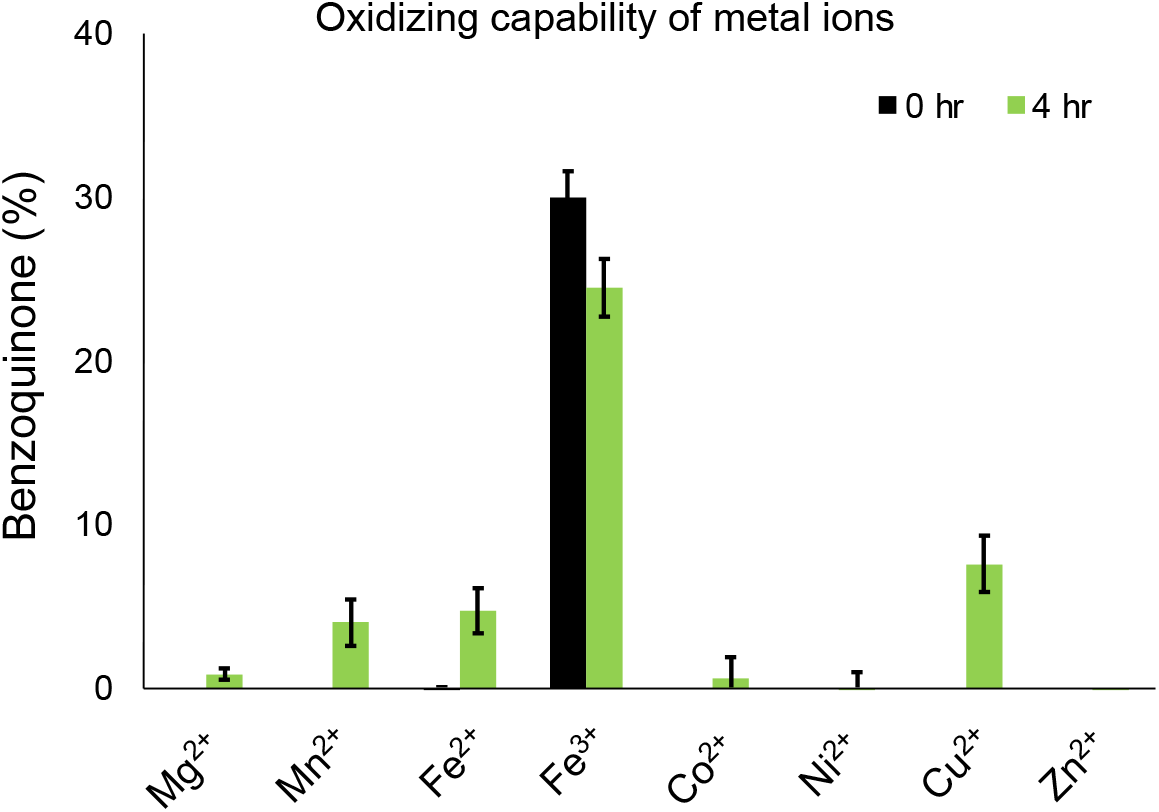
Oxidation of HQ to BQ in the presence of different metal ions. Bar graph shows the percentage of benzoquinone (BQ) produced, from oxidation of hydroquinone (HQ), with different metal ions used in 1:1 ratio of HQ to metal ions. Y-axis shows the percentage of benzoquinone (BQ) produced and X-axis shows the different metal ions used. Black and green colors indicate % BQ produced at the initiation (0 hr) and at the end (4 hr) of the reaction. Error bars depict standard deviation; N=3.

The standard reduction potential associated with the oxidation of HQ to BQ for 2 e^−^/ 2 H^+^ and 1 e^−^/ 1 H^+^ are 0.643 V and 0.099 V, respectively^37^. In order to oxidize HQ, the reduction potential of the oxidizing agent (metal ions) needs to be higher (more positive) than the reduction potential of BQ. The standard reduction potentials for the transition of Fe^3+^ to Fe^2+^ and Cu^2+^ to Cu^1+^ are 0.771 V and 0.159 V, respectively^38–40^ (Table 1), potentially justifying the ability of Fe^3+^ and Cu^2+^ to oxidize HQ. However, the slight oxidation observed in Fe^2+^ (−0.44 V) and Mn^2+^ (−1.170 V) could be attributed to their oxidation ability under aqueous condition. Altogether, the ability of even free metal ions to oxidize HQ, further emphasized the influence of these ions on these prebiotically pertinent reactions, prior to the emergence of more efficient and complex (proto) metalloenzymes^7,19,20,41–44^.

**Table 1:**
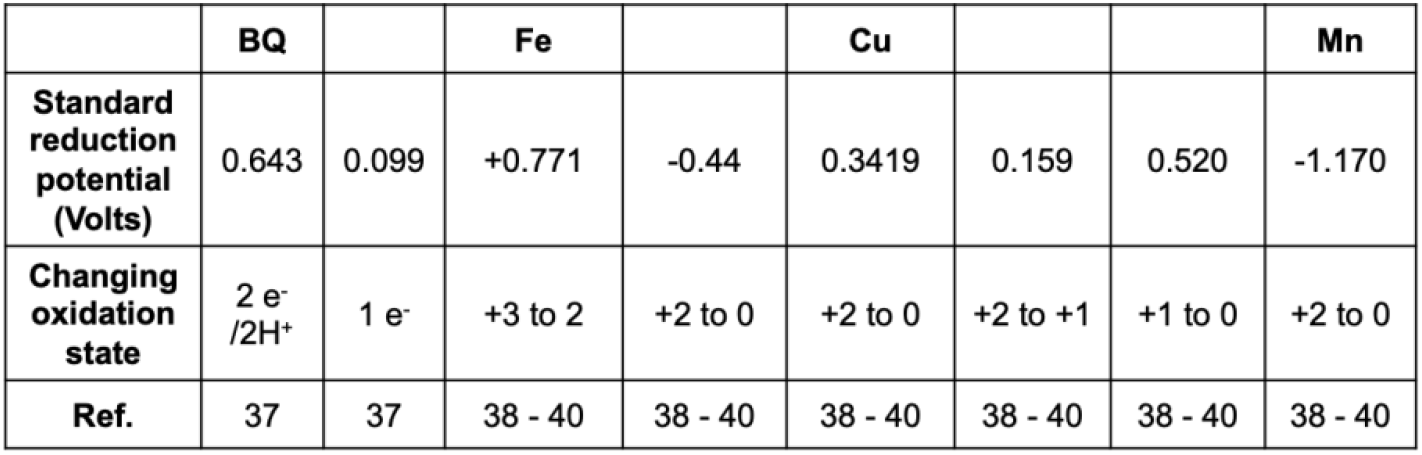
Reduction potentials for Cu, Mn and Fe with their respective oxidation state change; adapted from references 37 – 40 (Supplementary Table 1 contains the extended table with the reduction potentials of all the metal ions that we have used in our studies).

### Reactions containing different metal ions along with TPPS as a co-solute

Taking into account the complex nature of prebiotic soup, metal ions would have co-existed with other relevant organic co-solutes, including scaffolds similar to porphyrins^45^. The interactions between metal ions and such scaffolds might have impinged on the catalytic ability of these metal ions. To study this, we evaluated the effect of TPPS (tetraphenyl porphyrin tetra sulfonic acid) scaffold as a co-solute, in the reaction mixtures containing the different metal ions (see Methods).

As seen in Fig. 3 panel A, negligible oxidation of HQ was observed in the control reactions containing just HQ and HQ with TPPS in the absence of any metal ions. In the ‘co-solute reactions’, the oxidizing ability of Mg^2+^, Ni^2+^, Fe^2+^ and Zn^2+^ ions were found to be unaltered (Fig. 3, panel A). Interestingly, we observed significantly enhanced activity for Co^2+^ and Fe^3+^ ions in the presence of TPPS as a co-solute, as shown in Fig. 3, panel A (Supplementary Figs. 3 and 4, panels A-D; Supplementary Table 10-12). In the case of Co^2+^ and TPPS based co-solute reaction, up to ~12.3% of BQ was produced. Whereas, in the case of only free Co^2+^ ions, and only TPPS containing reaction, negligible HQ oxidation was observed (Fig. 2 and Fig. 3 panel A, respectively). In the Fe^3+^ and TPPS based co-solute reaction, the HQ oxidation was enhanced to ~51.7% just at the initiation of the reaction. Whereas, in the case of only Fe^3+^ ions, up to 30% HQ oxidation was observed. Interestingly, in the case of Mn^2+^ and Cu^2+^ containing TPPS co-solute reactions, the BQ production was found to be diminished when compared to only Mn^2+^ ions (~4%) and Cu^2+^ ions (~7.6%) containing reactions (Fig. 2, Fig. 3 panel A and Supplementary Fig. 2). This was in contrast to what was observed with the other metal ions, wherein the oxidation either increased or remained unchanged in the presence of TPPS. Importantly, a color change (from green to red) was also observed in co-solute reactions containing Fe^3+^, Co^2+^, Cu^2+^ and Zn^2+^ ions, over the course of the reaction.

**Figure 3:**
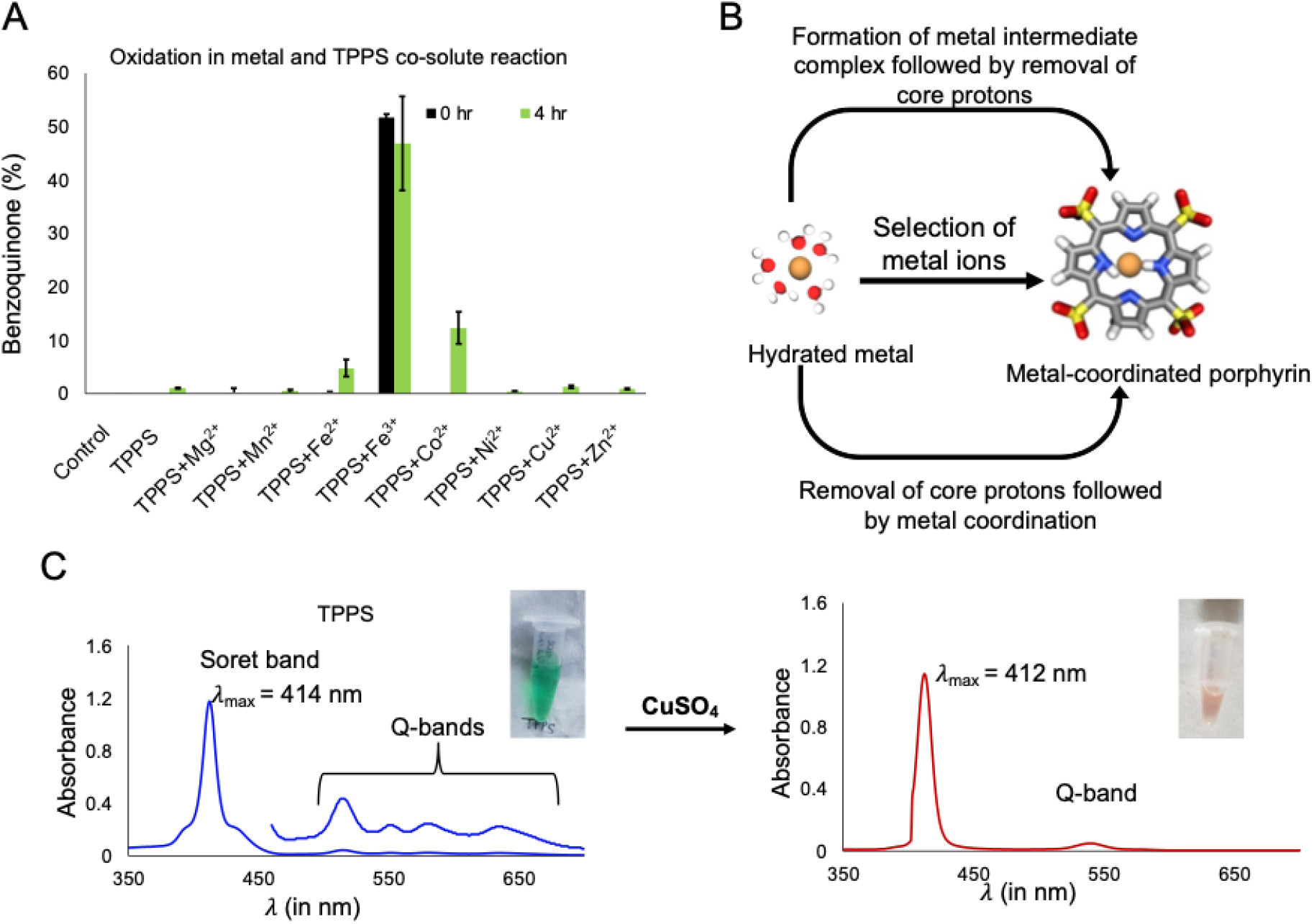
Oxidation of HQ to BQ in TPPS and metal ion containing co-solute reactions (panel A). The bar graph shows the percentage of benzoquinone (BQ) produced in the co-solute reactions. X-axis shows the different reactions i.e., only HQ control (control), HQ with TPPS (TPPS), and the different co-solute reactions containing HQ and TPPS along with the different metal ions viz. Mg^2+^ (TPPS+ Mg^2+^), Mn^2+^ (TPPS+ Mn^2+^), Fe^2+^ (TPPS+ Fe^2+^), Fe^3+^ (TPPS+ Fe^3+^), Co^2+^ (TPPS+ Co^2+^), Ni^2+^ (TPPS+ Ni^2+^), Cu^2+^ (TPPS+ Cu^2+^) and Zn^2+^ (TPPS+ Zn^2+^), respectively. Error bars depict standard deviation; N=2. Black and green colors indicate % BQ produced at the initiation of reaction (0 hr) and at the end (4 hr) of the reaction, respectively. B) Schematic showing the two mechanisms by which a metal ion can coordinate with porphyrin scaffold, to form a metal-porphyrin coordinated complex (M-TPPS). C) Representative UV spectrum of TPPS alone (left panel) and Cu^2+^-TPPS (right panel), showing changes in the absorbance spectrum on the metal’s coordination to porphyrin. Typically, change in the λ_max_ of the soret band and decrease in the number of Q bands was observed upon complexation.

Next, in order to investigate the underlying molecular change, leading to the change in the color and oxidation activity for Fe^3+^, Co^2+^ and Cu^2+^ ions in the presence of TPPS as a co-solute, UV absorbance spectroscopy was used. Free (uncomplexed) TPPS has an absorption maximum around 414 nm due to the transition of electrons from ground state (S_0_) to second excited state (S_2_), along with four weak Q bands in the 500-700 nm region (due to S_0_ to S_1_ transitions)46 (Fig. 3 panel C; left graph and Supplementary Fig. 10). A change in the λ_max_ of TPPS was observed for the Co^2+^ (to 427 nm), Cu^2+^ (to 412 nm), Zn^2+^ (to 421 nm) and Fe^3+^ (to 432 and 493 nm) containing co-solute reactions. Additionally, a decrease in the number of Q-bands in the 500-700 nm region was also observed, which also reflected in the color change for all the aforementioned reactions (Fig. 3 panel C; right graph shows this change for Cu^2+^). Similar changes in the UV spectra of TPPS was also observed when incubated with Co^2+^, Cu^2+^, Zn^2+^ and Fe^3+^ ions even in the absence of HQ (Supplementary Table 3). All these observations indicate that the change in the activity of specific metal ions observed was potentially due to the coordination between metal ions and TPPS. This implied that for specific metal ions, the coordination with TPPS occurs readily at 40°C (as reflected in the change in the UV spectrum)^47,48^, which resulted in a change in their catalytic activity.

### Prebiotic synthesis of preformed metal-porphyrin complexes

We next investigated the ability of metal ions to coordinate with TPPS and result in metal-coordinated TPPS (M-TPPS) complexes under prebiotically plausible terrestrial geothermal conditions. These niches are generally found to harbor a range of temperature and pH^49^, which makes them conducive for the formation of such M-TPPS complexes as described in this study. Aqueous solution containing 0.3 mM metal ions and 0.03 mM (ten times lower than metal ion) TPPS was incubated at 70° C, simulating putative early Earth conditions (details in Methods section). Metal coordination with the TPPS scaffold was monitored using the temporal change in the UV absorption spectrum and steady-state fluorescence spectroscopy, at specific time points as indicated (Supplementary Figs. 11-22). The decrease in the intensity at 414 nm (λ_max_ of soret band for TPPS), along with the appearance of other prominent bands with λ_max_ corresponding to M-TPPS complexes, was monitored as an indication of metal coordination46–48. A change in λ_max_ was observed for all metals except for Mg2+. The λ_max_ of TPPS (414 nm) shifted to 467 nm for Mn^2+^, 425 nm for Co^2+^, 410 nm for Ni^2+^, 412 nm for Cu^2+^ and 421 nm for Zn^2+^ (Supplementary Figs. 11-16). Additionally, a decrease in the number of Q-bands was also observed upon complexation. However, Fe^3+^ ions gave rise to two strong bands at 432 nm and 493 nm (Supplementary Fig. 12), with no Q bands in between 500-700 nm when incubated with TPPS. In the case of Mg^2+^, no change was observed in absorbance even after thirty-six hours of incubation at 70°C.

Uncomplexed TPPS, upon excitation at 414 nm, shows emission bands at 640 nm and 697-698 nm^47,50^. After coordinating with incompletely filled d-orbital containing metals, its fluorescence gets quenched^47^. This could be due to the paramagnetic nature of these metal ions, which allows excited electrons to lose their energy by intersystem crossing, thereby quenching the fluorescence of TPPS. Mn^2+^, Fe^3+^, Co^2+^, Ni^2+^ and Cu^2+^ ions that we used in this study, quenched the fluorescence of TPPS upon coordination due to their paramagnetic nature (Supplementary Figs. 17-21). This allowed us to examine the M-TPPS complex formation by monitoring the disappearance of the fluorescence of TPPS, for all the metals that we investigated (except Zn^2+^). In the case of Zn^2+^, the Zn^2+^-TPPS complex showed a blue shift in its emission bands (605 nm and 656 nm) upon excitation at 414 nm^47,50^ (Supplementary Fig. 22). Therefore, formation of Zn^2+^-TPPS complex was monitored by observing the blue-shift of the fluorescence emission bands (Supplementary Fig. 22). In the case of Mg^2+^, the fluorescence spectrum of TPPS was found to be unaffected even after thirty-six hours of incubation. The formation of coordinated M-TPPS was also confirmed using High-Resolution Mass Spectrometry (HRMS) (Supplementary Table 4). Our results showed that all the metal ions investigated here, excepting for Mg^2+^ and Fe^3+^, could coordinate with TPPS to form M-TPPS under prebiotically pertinent conditions. This Mg^2+^ observation was in agreement with what has been reported in a previous study, where it failed to coordinate with uroporphyrin in neutral aqueous conditions^23^. Importantly, the M-TPPS formation even in the micromolar range and under high temperature and aqueous conditions, underscores the prebiotic plausibility of these M-TPPS complexes.

To investigate the formation kinetics of M-TPPS complexes, fluorescence quenching of TPPS (or shifting in the emission maxima in case of Zn^2+^) was monitored (Fig. 4). TPPS fluorescence was quenched within one hour for Cu^2+^, Mn^2+^, Co^2+^ and Zn^2+^ containing reactions. Strangely, the fluorescence intensity was found to increase with increasing period of incubation in the case of Fe^3+^ ions, confirming the absence of the coordination between Fe^3+^ ions and TPPS (Fig. 4, panel A). In the case of Ni^2+^, the TPPS fluorescence was quenched completely only after thirty-six hours of incubation, confirming the formation of Ni-TPPS. This also indicated lesser affinity of Ni^2+^ ions towards porphyrin when compared to Cu^2+^, Mn^2+^, Co^2+^ and Zn^2+^ ions. In the case of Cu^2+^, the fluorescence intensity quenched immediately after the addition of Cu^2+^ to the TPPS solution (Fig. 4 panel A). In order to compare the affinity of Mn^2+^, Co^2+^ and Zn^2+^ ions towards TPPS, fluorescence intensities were monitored for 60 minutes (Fig. 4 panels B, C and D). The formation of Mn-TPPS was found to reach completion after 30 minutes, whereas for Co^2+^ and Zn^2+^ ions, the coordination completed within just 15 minutes.

**Figure 4:**
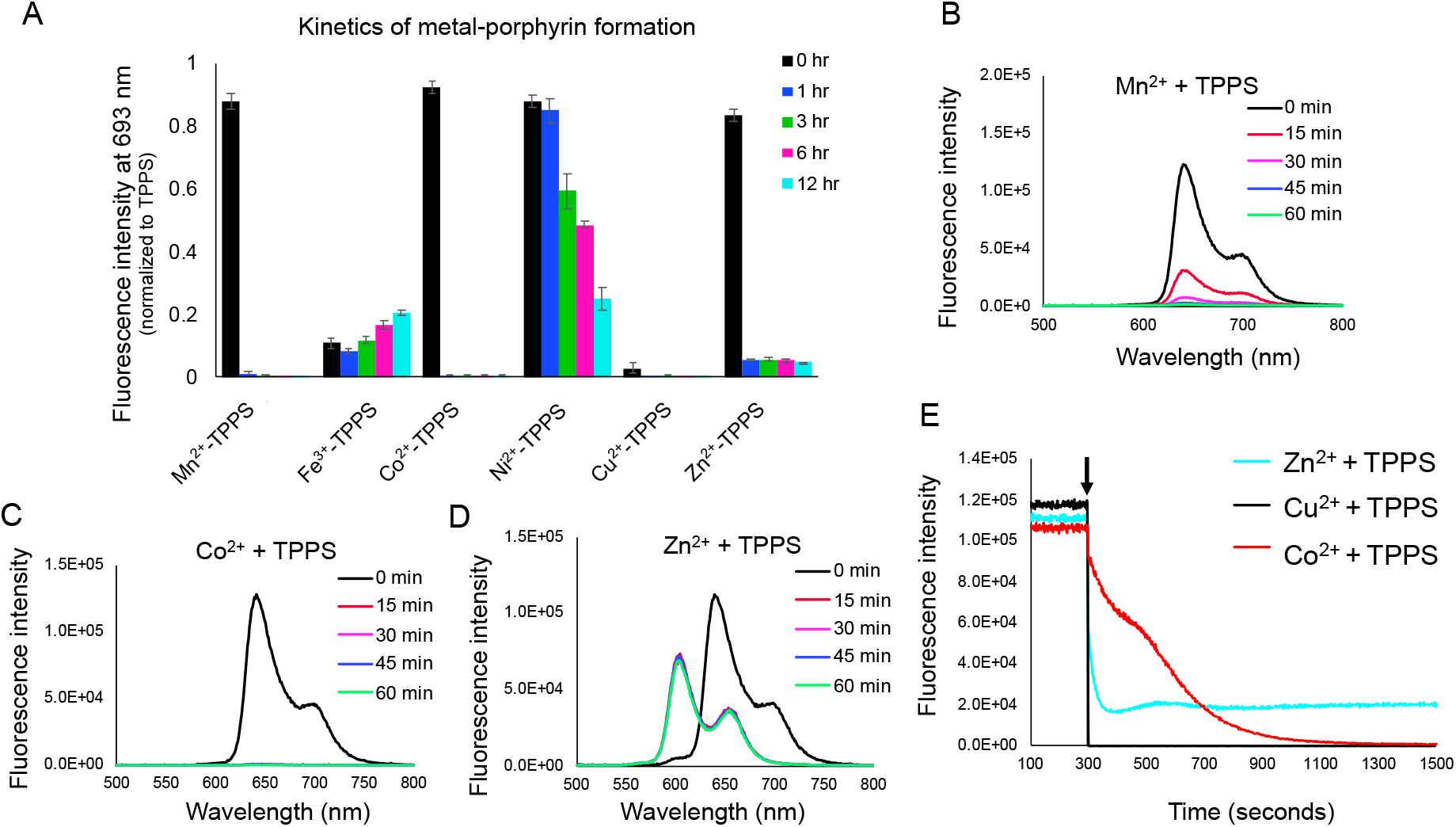
Formation kinetics of metal-coordinated TPPS (M-TPPS) complexes using fluorescence spectrum (λ_ex_ = 414 nm). A) Time-based fluorescence quenching of TPPS at 693 nm in the presence of different metal ions. Panels B, C and D show the fluorescence spectrum for the TPPS solution containing Mn^2+^, Co^2+^ and Zn^2+^, respectively, after different time periods of incubation (depicted in different colors as indicated above). E) Shows the fast-kinetics for the fluorescence quenching of TPPS at 693 nm in the case of Zn^2+^ (cyan curve), Cu^2+^ (black curve) and Co^2+^ (red curve). Black arrow indicates the timepoint of addition of corresponding metal ions to the TPPS solution after 300 seconds. N = 3.

To compare the affinity of the faster coordinating ions i.e., Cu^2+^, Zn^2+^ and Co^2+^ towards TPPS, these reactions were monitored continuously at a time interval of 0.1 seconds, using steady state kinetics (Fig. 4, panel E and Supplementary Figs. 29-31). The fluorescence of TPPS was monitored for the first 300 seconds, following which metal salt solution was added to reach a final concentration of 0.3 mM metal ions and 0.03 mM TPPS, and the solution was mixed thoroughly (Fig. 4, panel E; arrow depicts the addition of metal salt solution). The fluorescence was then monitored till 1500 seconds. The kinetic (fluorescence) decay data was fitted with first-order exponential decay curve to calculate the decay time constants. For Cu^2+^, Zn^2+^ and Co^2+^, the decay time constants were found to be < 4 seconds, 6.56 ± 1.13 seconds and 334.54 ± 42.93 seconds, respectively (Supplementary Figs. 29-31). The coordination rate (affinity) of metal ions with TPPS was in the following order; Cu^2+^ > Zn^2+^ > Co^2+^ > Mn^2+^ >>> Ni^2+^. This trend remained the same even in the competition experiments which contained a mixture of multiple metal ions in a single pot (see Methods, Supplementary Fig. 32).

Porphyrins are proposed to coordinate with metal ions via two possible mechanisms (Fig. 3 panel B)^21,51–53^. One is via the formation of a metal-TPPS intermediate complex, followed by the removal of the TPPS core protons, to yield the M-TPPS complex. The second mechanism occurs via a base-catalyzed removal of the TPPS core protons, subsequently followed by metal coordination. Both of these mechanisms depend on the capability of the metal ions to get rid of their hydration shell, typically known as ‘deconvolution’, to result in “naked” metal ions. Magnesium has been suggested to follow the base-catalyzed mechanism for coordination^21,53^.

This potentially explains the absence of Mg-TPPS formation under our reaction conditions (which were carried out at near neutral pH). Also, the trend observed for other metal ions could stem from their ease of deconvolution and the formation of intermediate complexes with the porphyrin scaffold.

### Oxidizing capability of preformed metal complexes

We observed the formation of M-TPPS complexes for all the metal ions investigated except for Mg^2+^ and Fe^3+^ under our reaction conditions. However, the rate of coordination with TPPS varied greatly for different metal ions, ranging from seconds (Cu^2+^and Zn^2+^) to minutes (Co^2+^ and Mn^2+^), to even days (Ni^2+^). Nonetheless, this confirms the ready availability of the different M-TPPS complexes in relevant geological niches of the early Earth^27,28^. Given this, we investigated the oxidizing capability of pre-formed metal-TPPS (M-TPPS) complexes, to characterize the influence of TPPS coordination on the oxidizing activity of the metal ions. Fe^3+^-TPPS was the only one that was acquired commercially as it failed to form under our simulated primitive Earth reaction conditions.

#### A. Oxidation of HQ to BQ by different M-TPPS complexes

In the M-TPPS mediated oxidation reactions, the molar ratio of HQ to M-TPPS was kept at 10:1; similar to the ratio that was used in the co-solute reactions (see Methods section). No significant HQ oxidation was observed in the presence of Zn^2+^-TPPS, Mn^2+^-TPPS and Ni^2+^-TPPS (Fig. 5 panel A). For Cu^2+^-TPPS, up to ~4% of HQ oxidation was observed after four hours. Interestingly, we observed significantly enhanced HQ oxidation, of up to 46%, in the presence of Co^2+^-TPPS complex after four hours, showing >4 turn over number (TON) (Fig. 5, panels A and B and Supplementary Table 14). This shows that upon coordination with TPPS, the catalytic activity of Co^2+^ ion increases remarkably. In the case of Fe^3+^-TPPS, ~10.4% HQ oxidation (TON =1) was observed immediately after addition of the M-TPPS complex, which remained unchanged even after four hours of incubation (Fig. 3 panel B). However, this percentage of HQ oxidation by Fe^3+^-TPPS (10.4%) was lower when compared to HQ oxidation that was facilitated just by the free Fe^3+^ ions (~ 30%) and what even in the co-solute reaction of Fe^3+^ and TPPS (51%) (Fig. 5 panel D).

**Figure 5:**
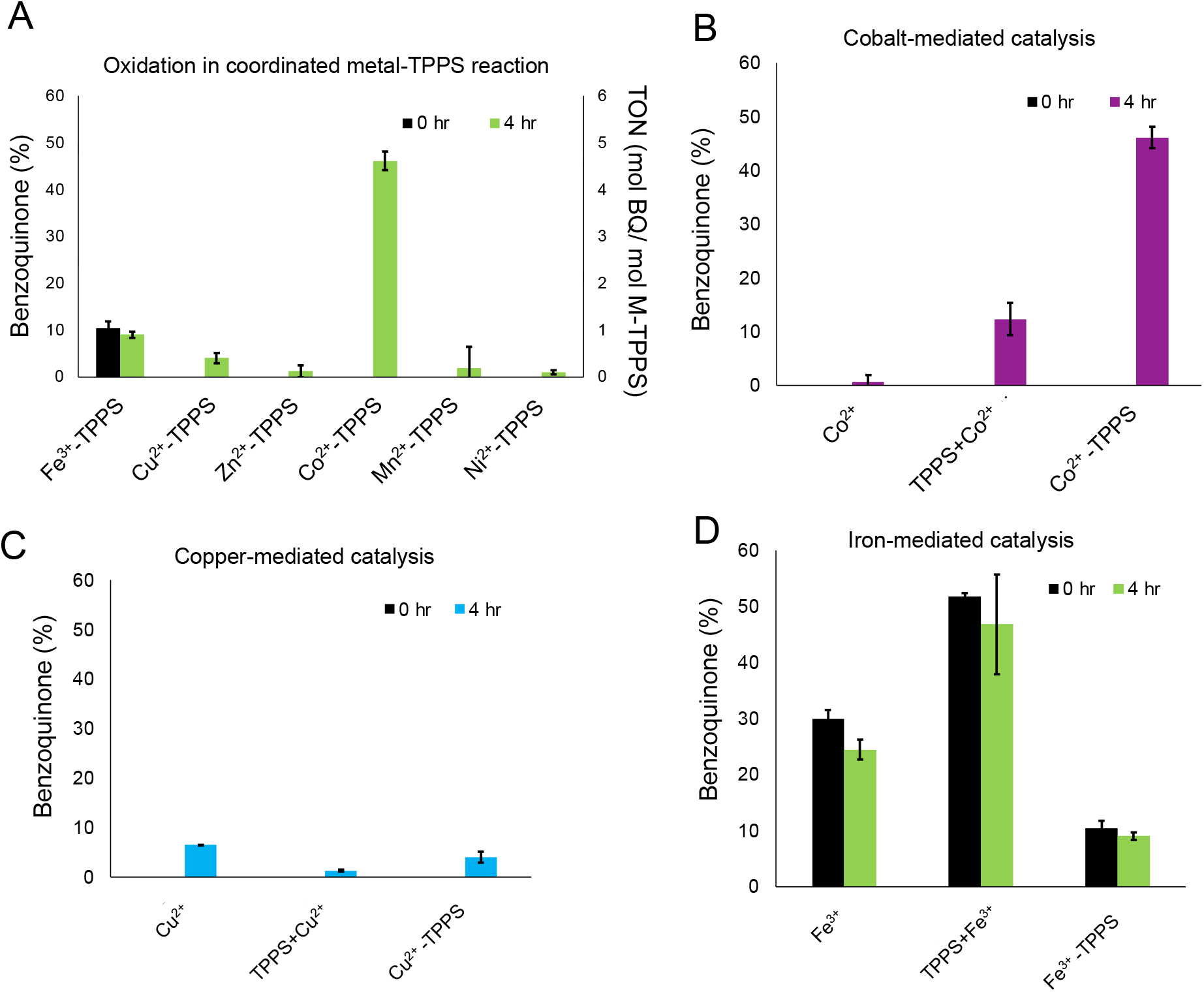
Oxidation of HQ by coordinated metal-TPPS (M-TPPS) complexes. A) Reactions of HQ with the respective M-TPPS complexes. Left Y-axis shows BQ (%) produced and right Y-axis shows the turn over number (TON) at the initiation (0 hr, black bars) and at the end of the reaction (4 hr, green bars). Panels B-D show the comparison of oxidation by free metal ions vs the respective metal ion with TPPS in a co-solute reaction vs the coordinated porphyrin complex (indicated in the X-axis). Time points shown are from the initiation of the reaction (black bars) and after four hours of incubation for Co^2+^ (panel B; purple bars), Cu^2+^ (panel C; blue bars) and Fe^3+^ (panel D; green bars) based reactions, respectively. N=2; error bars depict standard deviation.

The decrease in the activity of Cu^2+^-TPPS and Fe^3+^-TPPS when compared to the free metal ions, could be because of the formation of near-planar M-TPPS coordinated complexes as observed in the 3D model, which was predicted by energy minimization simulations using MolView (Supplementary Figs. 34 and 35)^54,55^. The presence of the rigid planar macrocycle ring of TPPS is known to restrict the metal’s ability to change its stereochemistry and hence, its oxidation state^56^. However, in the case of Co^2+^, a distortion in the planarity of porphyrin was observed upon coordination in the 3D model and this can allow the change in the stereochemistry of Co^2+^-TPPS, which, in turn can potentially enhance its catalytic ability (Supplementary Fig. 33). Nevertheless, the lower oxidizing activity of the coordinated Fe^3+^-TPPS still does not justify the high oxidation capability that was seen in the co-solute reaction containing Fe^3+^ and TPPS. Moreover, from UV and fluorescence spectroscopy, it was confirmed that Fe^3+^ was not able to coordinate with TPPS to form the Fe^3+^-TPPS complex, under our co-solute reaction conditions (Fig. 6 panel A).

**Figure 6:**
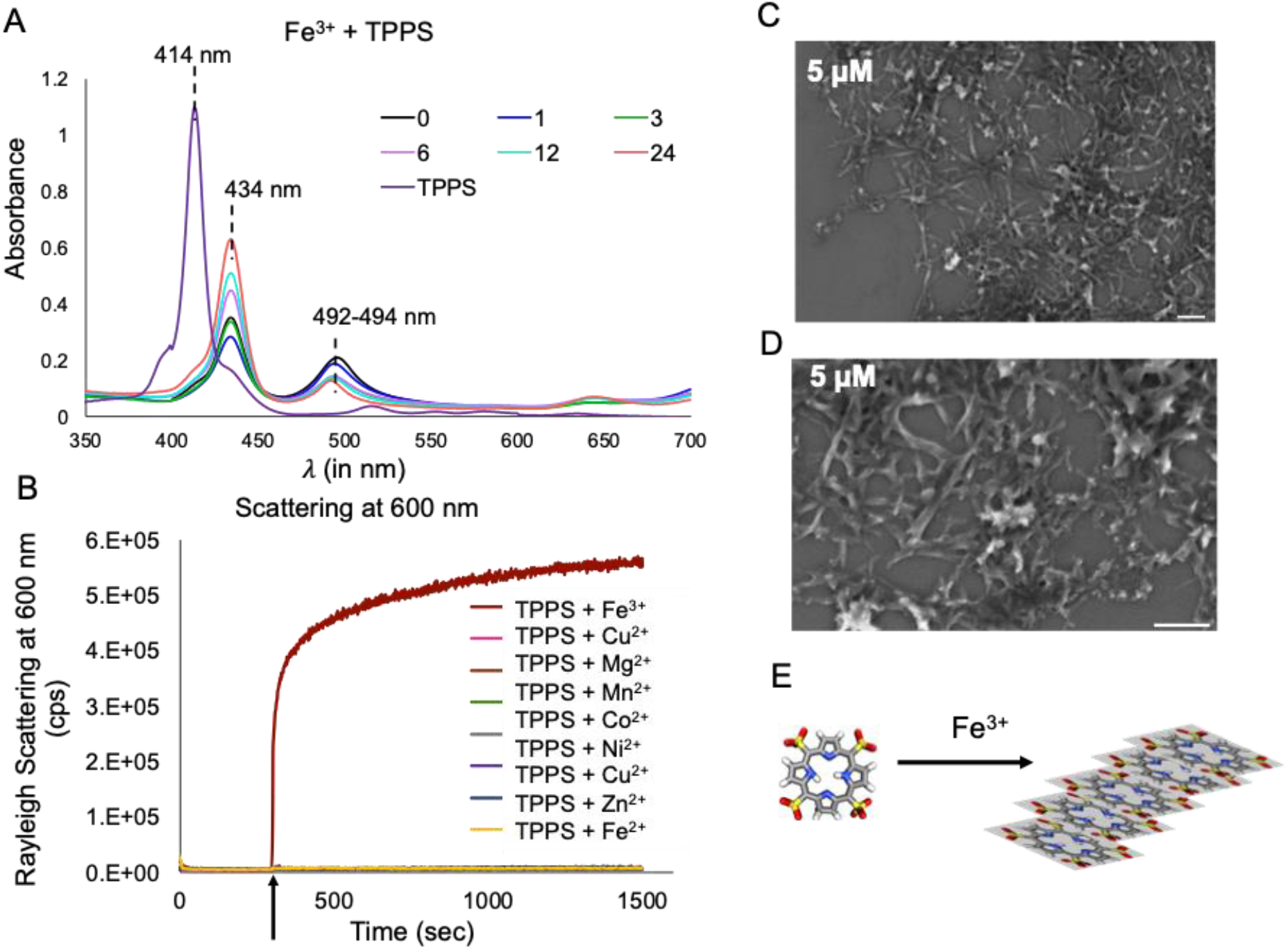
Characterization of Fe^3+^ and TPPS non-coordinated aggregates (TPPS**Fe^3+^). A) UV absorbance spectrum of co-solute reaction containing Fe^3+^ and TPPS, after different time periods (at the initiation (0), after 1 hour (1), 3 hours (3), 6 hours (6), 12 hours (12) and 24 hours (24), respectively) along with TPPS as reference. Different colors depict different time periods as shown in the legend; purple color is the UV spectrum of the reference TPPS sample. The red shift of 414 nm soret band to 434 nm and 492 nm bands was observed, indicating the formation of J-aggregates. B) Rayleigh scattering at 600 nm confirming the formation of higher order structures only in the Fe^3+^ and TPPS containing co-solute reaction. The black arrow indicates the time when the addition of different metal ions to the TPPS solution was done. Different colors represent the kinetic curve obtained after the addition of the different metal ions (as shown in the Figure legend). N=3. Panel C shows the Field Emission Scanning electron microscopy (FESEM) image of 5 µM of TPPS solution containing 50 µM of Fe^3+^ ions, along with its magnified version in panel D. The scale bar is 500 nm. Appearance of elongated rod-like structures confirmed the presence of J-aggregates. E) Illustration showing the Fe^3+^ induced formation of J-aggregates of TPPS.

To unmask the reason behind the increased catalytic activity of the co-solute reaction containing Fe^3+^ and TPPS, we chose to systematically explore it further. We assessed the possibility of the formation of non-coordinated Fe and TPPS complexes (TPPS**Fe^3+^) via Fe^3+^ ion interaction with the TPPS moiety. Visible precipitates were also observed in this co-solute reaction upon centrifugation at 5000g for 2 minutes, indicating the presence of large aggregates in the solution. The formation of these aggregates was further investigated by looking at the scattering of this solution at 600 nm (the mixture does not absorb light at this wavelength). We observed a drastic increase in the scattering of the TPPS containing solution upon addition of the Fe^3+^ ions, signifying the formation of higher order structures (Fig. 6 panel B). Additionally, the presence of aggregates in the solution were confirmed using optical microscopy (for details see Methods section). Large clusters of rod-shaped aggregates were observed for the mixture containing TPPS and Fe^3+^, while the control solutions containing Fe^3+^ alone, TPPS alone or preformed Fe^3+^-TPPS alone failed to show any aggregation (Supplementary Fig. 41). From previous literature, it is known that porphyrin complexes are known to predominantly form side-by-side aggregates (J-aggregates) and plane-to-plane aggregates (H-aggregates)^50,57–61^. Studies have also suggested that the catalytic activity of porphyrin increases upon aggregation^59,60,62^. Given this, it seemed like a reasonable premise to base our assumption that the increase in the catalytic activity in the co-solute reactions containing Fe^3+^ and TPPS, could possibly be because of the formation of aggregates.

Upon formation of J-aggregates and H-aggregates, the soret absorption band of porphyrin is known to undergo red-shift and blue shift, respectively^57,59,63,64^. The absorption λ_max_ of J-aggregates of TPPS was reported to be near ~ 490 nm in aqueous solution ^57,59,63,64^. The co-solute reaction containing Fe^3+^ and TPPS showed a red-shifted band with a λ_max_ at 494 nm, alluding to the formation of TPPS**Fe^3+^ J-aggregates (Fig. 6 panel A). The presence of J-aggregates was further confirmed by Field emission Scanning Electron Microscopy (FESEM), followed by the elemental analysis of the observed aggregates (Fig. 6 panels C and D, Supplementary Figs. 42 and 43). The presence of elongated planar structures, confirmed the presence of J-aggregates^59^. Further, the elemental analysis of these structures showed the presence of Fe, C, N, O and S, indicating that the structures visible in the viewing field were indeed aggregates containing Fe^3+^ and TPPS moieties (Supplementary Fig. 43). Moreover, prolonged heating of the TPPS**Fe^3+^ aggregates at 100°C, resulted in the diminishing of the absorption peak at 494 nm that corresponds to the J-aggregates (Supplementary Fig. 44). This could result from the instability of the aggregated species at this high temperature, thereby forming Fe^3+^-TPPS coordinated complex instead. Given this, in the case of Fe^3+^-TPPS formation, it seems that the coordination is formed via the attack of Fe^3+^ ion on the aggregated TPPS at elevated temperature (100°C). Interestingly, such a mechanism has been proposed for the formation of coordinated Cu^2+^-porphyrin complex in the presence of porphyrins in its aggregated form (J-aggregates)^65^. Additionally, whether the other metal ions studied here (Mn^2+^, Fe^2+^, Co^2+^, Ni^2+^, Cu^2+^ and Zn^2+^), could also form similar aggregates with TPPS, was evaluated by looking at the scattering of their respective solutions (Fig. 6 panel B). No increase in solution scattering, along with absence of visual aggregates upon centrifugation, and the absence of absorption peaks corresponding to J- and H-aggregates, eliminated the possibility of the other metal ions forming non-covalent aggregates with TPPS.

#### B. Oxidation of NADH to NAD by M-TPPS complexes

The spontaneous emergence of catalysts on the early Earth would have been governed by a multitude of factors, making their ready emergence a non-trivial exercise^3^. Thus, the early protoenzymes would possibly have evolved the ability to catalyze a wide array of prebiotically relevant reactions. Towards this, we investigated the ability of M-TPPS to catalyze oxidation of NADH at 25° C (see Methods section). Along with quinones, NADH is another prebiotically plausible cofactor that is involved in extant redox reactions and is a key molecule in metabolism^15,66,67^. This was chosen as a representative of nucleotide-based cofactors like flavin adenine dinucleotide (FAD) and flavin mononucleotide (FMN), which are utilized extensively in contemporary biology. NAD is characterized by a UV absorption band at 260 nm. On the other hand, NADH absorbs at 340 nm in addition to 260 nm. Therefore, in order to monitor the oxidation of NADH, the disappearance of 340 nm absorption signal was monitored. In the control reactions, i.e., in the absence of M-TPPS, only 1.5% oxidation was observed even after four hours of incubation, indicating negligible spontaneous oxidation of NADH (Supplementary Fig. 45). Our results showed that up to 12.3% NADH was oxidized immediately after the addition of Co^2+^-TPPS to the NADH solution (Fig. 7 and Supplementary Fig. 48). The NADH oxidation was observed to increase gradually and reach up to 59.9% after four hours. Additionally, the control reactions containing either the free Co^2+^ ions or just the TPPS scaffold showed negligible oxidation (Supplementary Figs. 46 and 47). Along with Co^2+^-TPPS, other M-TPPS were also observed to oxidize NADH to varying extent. The oxidation ability of different M-TPPS towards NADH followed this order: Co^2+^-TPPS (59.9%) > Cu^2+^-TPPS (52.8%) > Fe^3+^-TPPS (50.4%) > Mn^2+^-TPPS (29.3%) > Zn^2+^-TPPS (26.9%) > Ni^2+^-TPPS (12.9%) (Fig. 7). The M-TPPS mediated NADH oxidation displayed different TON ranging from ~6 for Co^2+^-TPPS to >1 for Ni^2+^-TPPS (Fig. 7). The capability of M-TPPS to oxidize NADH, emphasizes their versatility for using multiple substrates; a property that would have been hugely advantageous on the prebiotic Earth. A protracted process of molecular evolution over very long-time scales would have subsequently allowed for the emergence of highly specific and efficient catalysts for individual substrates.

**Figure 7:**
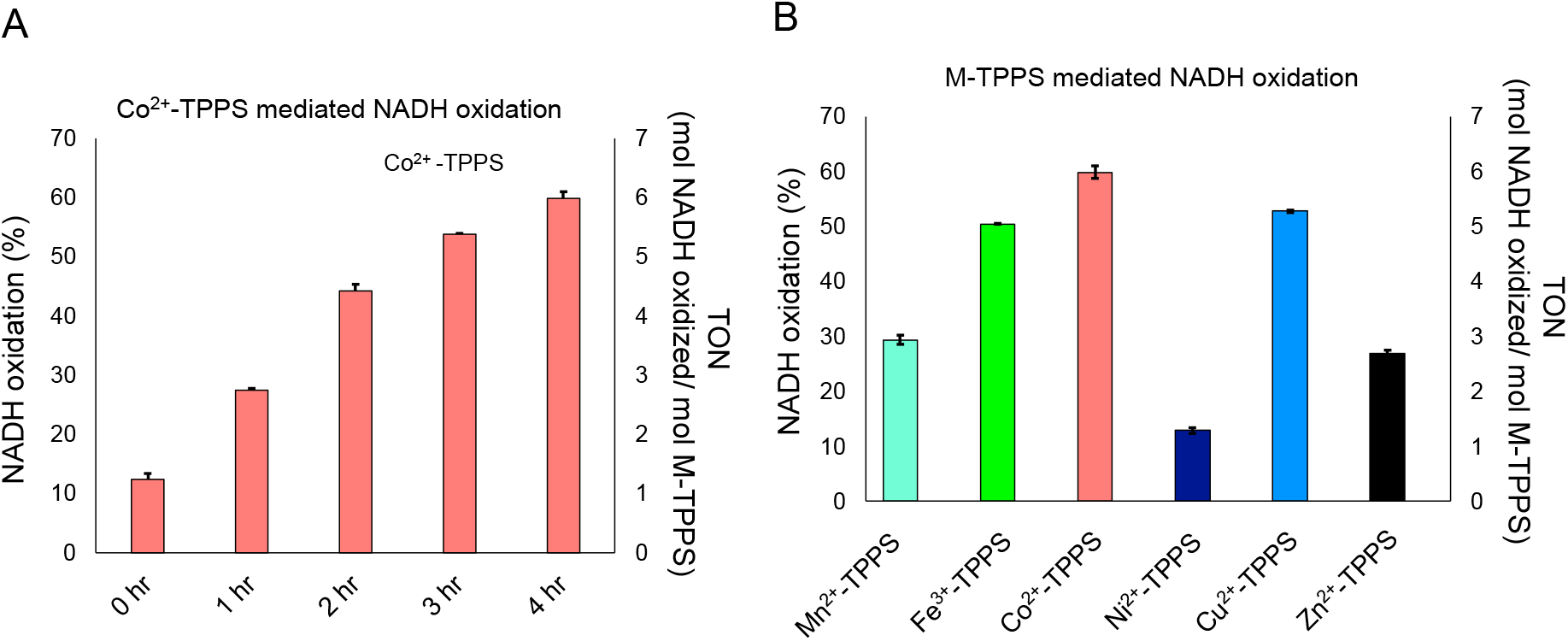
Oxidation of NADH mediated by preformed M-TPPS complexes. A) Co^2+^-TPPS mediated NADH oxidation after different time periods (salmon bars). X-axis shows the different time periods i.e., at the initiation of the reaction (0 hr), after one hour (1 hr), two hours (2 hr), three hours (3 hr) and four hours (4 hr) of incubation, respectively, at 25°C. Left Y-axis depicts the percentage of NADH oxidized and right Y-axis depicts the turn over number (TON) of Co^2+^-TPPS. B) Comparison of NADH oxidation mediated by different M-TPPS. Left Y-axis depicts the percentage of NADH oxidized and right Y-axis depicts the turn over number (TON) of the M-TPPS. N=2; error bars depict standard deviation.

## Discussion

Porphyrin-containing metalloenzymes are suggested to be some of the most ancient catalysts to be found on the early Earth^1,16,68^. However, their catalytic ability in prebiotically pertinent reactions, has largely remained unexplored. In this work, we investigated the influence of porphyrin scaffolds, on the modulation of oxidation ability of different metal ions. In case of free metal ion-mediated reactions, the highest oxidation was observed for Fe^3+^ followed by Cu^2+^, Fe^2+^ and Mn^2+^ ions (Fig. 2). In the co-solute reactions, the HQ oxidation was found to be increased for Fe^3+^ (from 30% to 51.7%) and Co^2+^ (from negligible to 12.3%) ions in the presence of TPPS, when compared to just the corresponding free metal ion-mediated oxidation (Fig. 5). Contrarily, the HQ oxidation by Cu^2+^ was found to diminish in presence of TPPS while it remained unaltered for the rest of the metal ions. Given this data, we conclude that the presence of TPPS as a co-solute increases the catalyzing activity of Fe^3+^ and Co^2+^ ions, while damping the activity of Cu^2+^ ions.

Changes in the UV-absorption and fluorescence emission profiles in the co-solute reactions, indicated that the change in the catalytic activity of the metal ions in the presence of TPPS stems from the interaction between the metal ions and the TPPS. This was further confirmed by similar changes seen in the activity of the pre-formed coordinated complexes of Cu^2+^-TPPS and Co^2+^-TPPS mediated oxidation reactions. In contrast to Co^2+^, inactivity of Cu^2+^-TPPS could be attributed to the inability of copper to change its oxidation state when it is coordinated to TPPS, which is a planar rigid structure^56^ (Supplementary Fig. 34). Importantly, this observation highlighted that the catalytic activity of metal ions, does not necessarily increase upon complexation, but rather can vary greatly between the metal ions. This underscores the interesting possibility that coordination/interaction of metals with organic scaffolds such as TPPS, could have acted as a selectivity barrier for the formation and functioning of protometalloenzymes. Importantly, the M-TPPS complexes were also found to catalyze the oxidation of NADH to NAD, signifying its substrate versatility. Similar to HQ based reactions, the highest oxidation of NADH was observed for Co^2+^-TPPS. This capability to recognize different types of substrates would have been beneficial on the early Earth where spontaneous emergence of efficient catalysts would have been a formidable task.

While characterizing the prebiotic plausibility of metalloporphyrins, we observed that Cu^2+^, Co^2+^, Mn^2+^, Ni^2+^ and Zn^2+^ ions were able to coordinate with TPPS under prebiotic settings. However, the rate of the formation of M-TPPS varied greatly as coordination with TPPS could occur in a few seconds (as was observed for Cu^2+^, Zn^2+^ and Co^2+^), or could even happen on the order of days (as was seen for Ni^2+^). Therefore, the affinity of the metal ions for the organic scaffold may have been another important factor by which selection of the metal ions would have come about. These selected metal ions would have then eventually used for driving catalysis by protometalloporphyrins on the early Earth.

Interestingly, the oxidizing capability of the Fe^3+^-TPPS (coordinated) complex was found to be lower when compared to HQ oxidation in the presence of free Fe^3+^ ions or the co-solute reactions that contained Fe^3+^ ions and TPPS. Instead of coordinating with TPPS, Fe^3+^ was found to form non-coordinated aggregates with TPPS under our reaction conditions. From UV spectroscopy, Rayleigh scattering and FESEM, Fe^3+^ ions were shown to assemble into J-aggregates with TPPS which potentially resulted in enhanced oxidation activity (increased by 1.7-fold when compared to the free Fe^3+^ ion containing reaction). Interestingly, approximately, 27% of extant oxidoreductase enzymes utilize Fe^3+^/ Fe^2+^ redox couple to catalyze reactions^69,70^. Previous studies have indicated that the metal ion availability in the Archean Ocean would have followed the order mentioned here: Fe^+3/+2^> Mn^+2^, Ni^+2^, Co^+2^>> Cd^+2^, Zn^+2^> Cu^+2 32,71^. Given this abundance of Fe^+3/+2^ on the prebiotic Earth, along with its ability to catalyze reactions on its own, it would have readily induced aggregation resulting in non-coordinated catalytically active metal-organic scaffolds. This potentially indicates a pivotal role for Fe^+3/+2^ in catalyzing prebiotically relevant reactions.

To summarize, our work demonstrates how change in oxidizing activity of different metal ions can be brought about by coordination/interaction with prebiotically plausible organic scaffolds like porphyrin (Fig. 8). We showed the influence of porphyrin in modulating the oxidizing ability of the free metal ions in positive or negative manner, depending on the metal ion in question. We also demonstrate the formation of different M-TPPS (coordinated) complexes and Fe^3+^ containing J-aggregates under pertinent prebiotic conditions. The eventual refinement of the usage of a specific ion type, in conjunction with a suitable scaffold would have resulted in highly precise and effective contemporary catalysts. This refinement would have resulted over millions of years of evolution. Various factors, starting from the availability of the metal ions, to the ease of formation of metal-organic scaffolds, to its influence on the actual catalytic activity, would have played an important role in the selection and evolution of metalloenzymes. In this context, our study provides preliminary insights as to why certain metal ions might have been selected for over the course of evolution. It also underscores the central role of the complex interactions and cross-talk that can occur in a heterogenous prebiotic soup between relevant molecules, and their potential role in the emergence and molecular evolution of early metalloenzymes.

**Figure 8:**
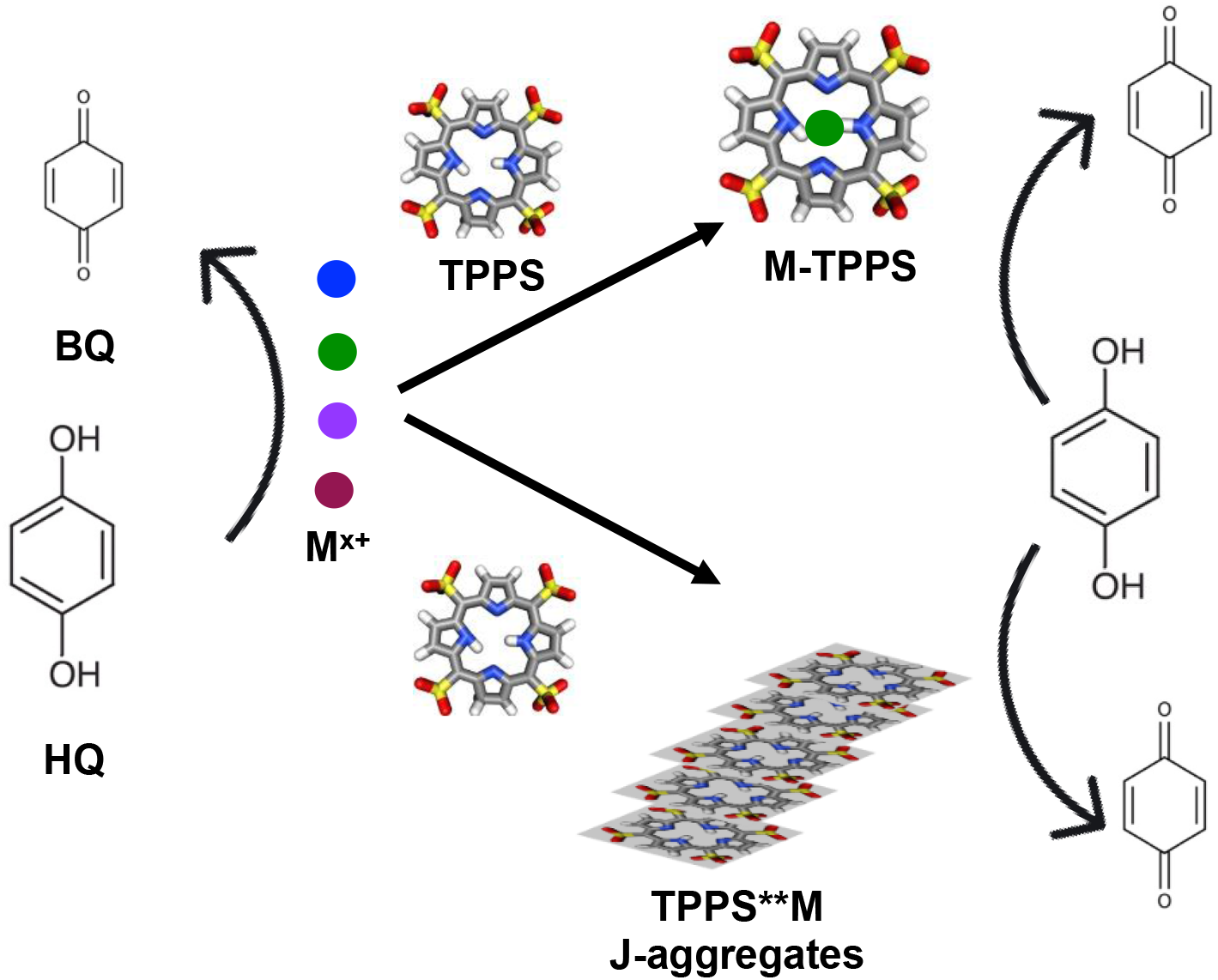
Schematic showing the two different pathways described in this study by which oxidation of HQ to BQ is made feasible. Different color spheres show the different metal ions. These metal ions in the presence of TPPS can then either coordinate to form M-TPPS complex or induce J-aggregate formation (TPPS**M; a non-coordinated complex). These complexes (coordinated or non-coordinated) have different catalytic efficiencies towards HQ oxidation.

## Materials and Methods

### Materials

All reagents were from Sigma Aldrich and used without further purification. The C18 chromatography column was purchased from Agilent Technologies (Santa Clara, CA, USA). All the oxidation reactions were performed in the anaerobic glove bag from Coy Labs under 95% N_2_ and 5% H_2_ (usually used to maintain anoxygenic conditions with oxygen levels < 200 ppm) environment. Nanopure water was filtered through 0.22-micron filter, degassed and purged with N_2_ to remove oxygen. The samples were transferred to glass vials covered with caps with air tight septum for HPLC-based analysis.

### Methods

High-performance liquid chromatography (HPLC): HPLC analysis was performed using a Shimadzu HPLC system (CBM-20 A, quaternary pump LC-20AD; Shimadzu Corporations, Kyoto, Japan). The analytes were chromatographically resolved in the reverse-phase (Zorbax Eclipse Plus C18 column; 4.6 X 150 mm, 3.5 μm particle size, Agilent). For separation of Hydroquinone (HQ)/ Benzoquinone (BQ), a gradient of methanol was used as the eluting solvent. The HPLC run started with 100% nanopure water and 0% methanol and at a flow rate of 1 ml/ min. The methanol was increased to 15% over 15 min and was kept at this percentage until 18 min. Following this, it was reduced to 0% by 19 min and the column was equilibrated at this concentration for 2 more minutes. The analytes were detected using a photo Diode-Array Detector (DAD). A standard solution of 0.2 mM HQ eluted at 5.5 minutes and 0.2 mM BQ eluted at 10.03 and 10.6 minutes, respectively. The BQ was quantified by integrating the corresponding HPLC peaks. For TPPS complex containing reactions, in addition to the 21 minutes method, 100% methanol was run from 22 min to 29 min followed by equilibration with nanopure water from 30 min to 33 min.

Oxidation reactions with metal salts: An aqueous solution of each metal sulfate salt (20 mM for Fe_2_(SO_4_)_3_ and 40 mM for other salts) was prepared. Unless otherwise reported, a typical reaction contained 0.3 mM hydroquinone (HQ) and 0.3 mM metal ion. For Fe^3+^, one molecule of Fe_2_(SO_4_)_3_ contains two atoms of Fe per formula unit. Therefore, in these reactions, 0.3 mM and 0.15 mM of sulfate salt concentration was used for to keep the molar ratio of HQ and metal ions constant as 1:2 and 1:1, respectively. The reaction mixture was then incubated at 40°C for four hours under anoxygenic conditions (O_2_ < 100 parts per million (ppm)). 30 µl of sample was withdrawn immediately after the addition of metal salt in the reaction mixture (0 hr.) and then again after four hours of incubation (4 hrs.), out of which 25 µl was loaded onto the column for HPLC-based analysis.

Oxidation reactions with metal salts and TPPS as co-solute: A typical reaction contained 0.3 mM HQ, 0.3 mM metal ion and 0.03 mM TPPS (1:1:0.1 molecular ratio). Control reactions with only HQ or containing only TPPS along with HQ were also performed. Other conditions such as incubation at 40°C for four hours were kept constant, following which the samples were analyzed using HPLC by loading 25µl of the respective sample at a given time.

UV-vis spectroscopy: A UV-1800 Shimadzu spectrophotometer was used to measure UV-vis absorption spectra (scan range from 350 nm to 700 nm, interval = 1 nm) for the samples after various times. UV-vis spectrum was recorded at room temperature.

Steady state fluorescence spectroscopy: A Fluoromax-4 spectrofluorometer (Horiba Scientific, Kyoto, Japan) was used for fluorescence measurements i.e., fluorescence quenching in the kinetics experiments and scattering in aggregation studies. For fluorescence-based assays the light with a wavelength 414 nm was used for excitation. The excitation and emission slits were fixed at 2 nm and 1 nm, respectively. The fluorescence intensity for the emission signals of a given sample was then scanned from 500 to 800 nm with 1 nm interval at 70°C, after different time periods of incubation.

For fast-kinetics experiments (in the case of Zn^2+^, Cu^2+^ and Co^2+^), the fluorescence intensity was measured at 693 nm for 0.03 mM TPPS solution at 70°C, till 300 seconds, after which 0.3 mM of corresponding metal ion was added and mixed. Following this, the fluorescence was monitored till 1500 seconds.

Competition experiments for the formation of M-TPPS: 0.03 mM TPPS was incubated with different mixtures of metal ions at 70°C and rotated at 350 rpm. The concentration of each metal ion was 0.03 mM. Four different combinations of different metal ions were used. Set 1 contained Mn^2+^, Fe^3+^, Co^2+^, Ni^2+^, Cu^2+^ and Zn^2+^. Set 2 contained Mn^2+^, Fe^3+^, Co^2+^, Ni^2+^ and Zn^2+^. Set 3 contained Mn^2+^, Fe^3+^, Co^2+^ and Ni^2+^. Set 4 contained Mn^2+^, Fe^3+^ and Ni^2+^. UV-vis absorption spectra of each of these combinations were recorded after different time periods to monitor the formation of different M-TPPS complexes.

For Rayleigh scattering to study aggregation, the light with a wavelength 414 nm was used for excitation and emission was also measured at 414 nm. The excitation and emissions slits were fixed at 2 nm. The fluorescence intensity was measured for 0.03 mM TPPS solution till 300 seconds, after which 0.3 mM of corresponding metal ions was added and mixed at room temperature. Following this, the fluorescence was then monitored till 1500 seconds.

Field-emission scanning electron microscopy (FE-SEM): FE-SEM images were recorded using Zeiss Ultra Plus scanning electron microscope. Approximately 2.5 µl of the sample composed of 5 µM TPPS and 25 µM of Fe_2_(SO_4_)_3_ (1:10 of TPPS: Fe^3+^ ion) was prepared by drop casting on silicon wafers.

Differential Interference Contrast (DIC) Microscopy: Microscopic analysis of TPPS**Fe^3+^ non-coordinated aggregates was done using a DIC microscope AxioImager Z1 (Carl Zeiss, Germany), (NA = 0.75) under 40X objective. Approximately 10 µl mixture of 0.03 mM TPPS and 0.15 mM Fe_2_(SO_4_)_3_ (1:10 of TPPS: Fe^3+^ ion) was spread on a glass slide and covered by an 18X18 glass coverslip. Followed by which the sides of the cover slip were sealed with liquid paraffin and was observed under microscope.

Oxidation reactions with metal-coordinated TPPS (M-TPPS) complexes: The reaction mixture contained 0.3 mM HQ and 0.03 mM (1:0.1 atomic ratio) of corresponding M-TPPS complexes. All other reaction conditions such as incubation at 40°C for four hours, were kept the same as mentioned above. The reactions were then analyzed using HPLC.

Co^2+^-TPPS mediated oxidation reactions of NADH: The reaction mixture contained 0.1 mM NADH and 0.01 mM (10: 1 molar ratio) of Co^2+^-TPPS complexes. The mixture was then incubated at 25°C for four hours under anoxygenic condition (O2 < 200 ppm). The reaction was analyzed using UV spectrophotometer by monitoring the disappearance of the signal at 340 nm, which corresponds to NADH absorption. Control reactions were performed i.e., spontaneous oxidation containing only NADH and reactions containing only Co^2+^ or only TPPS (Supplementary Figs. 45-47).

#### Statistical analysis

All statistical analysis was performed using Microsoft Excel 2016. Two-tailed t-test was used to check the significance of difference between the values within the reactions and also to compare between values of particular time points across reactions. Values were considered statistically significant for values with p < 0.05.

## Author contributions

S.D., S.S and S.R. designed the experiments. S.S. performed optical microscopy and the fluorescence-based kinetics experiments for studying M-TPPS formation. S.D. performed all the remaining experiments. S.D., S.S. and S.R. analyzed the data and wrote the manuscript.

## Acknowledgments

The authors wish to acknowledge the Microscopy, HRMS and FE-SEM facility at IISER Pune. This research was supported by grants from the Science and Engineering Research Board (SERB), Department of Science and Technology, Govt. of India [EMR/2015/000434], the Department of Biotechnology, Govt. of India [BT/PR19201/BRB/10/1532/2016] and IISER Pune. S.D. and S.S. acknowledge CSIR, Govt. of India and IISER Pune, respectively, for their fellowship.

