## Supplementary Information for "Porphyrin in prebiotic catalysis: Ascertaining a route for the emergence of early metalloporphyrins"

TABLE OF CONTENTS

TABLE OF CONTENTS …..........................................................................................2-4

EXPERIMENTAL DATA

Representative HPLC trace showing separation of Hydroquinone and Benzoquinone…5
HPLC trace showing oxidation of HQ in various $Cu^{2+}$ containing reactions………………6
HPLC trace showing oxidation of HQ in various of $Co^{2+}$ containing reactions…………...7
HPLC trace showing oxidation of HQ in various of $Fe^{3+}$ containing reactions……………8
Standard curve for Hydroquinone (HQ)………………………………………………………9
Standard curve for Benzoquinone (BQ)……………………………………………………..10
Oxidation of HQ in presence of $Cu^{2+}$ ions in varying ratios……………………………….11
Oxidation of HQ in presence of different metal ions in 1:2 ratio………………………12-13
Oxidation of HQ in presence of different ratios of $Cu^{2+}$ and $Fe^{3+}$…..............................14
UV spectrum of TPPS at different pH……………………………………………………15
UV absorption kinetics for $Mn^{2+}$-TPPS formation…………………………………………..16
UV absorption kinetics for $Fe^{3+}$-TPPS formation…………………………………………...17
UV absorption kinetics for $Co^{2+}$-TPPS formation…………………………………………..18
UV absorption kinetics for $Ni^{2+}$-TPPS formation……………………………………………19
UV absorption kinetics for $Cu^{2+}$-TPPS formation…………………………………………..20
UV absorption kinetics for $Zn^{2+}$-TPPS formation……………………………………..……21
Fluorescence quenching kinetics for $Mn^{2+}$-TPPS formation…………………………...….22
Fluorescence quenching kinetics for $Fe^{3+}$-TPPS formation………………………………23
Fluorescence quenching kinetics for $Co^{2+}$-TPPS formation………………………………24
Fluorescence quenching kinetics for $Ni^{2+}$-TPPS formation………………………………..25
Fluorescence quenching kinetics for $Cu^{2+}$-TPPS formation……………………………....26
Fluorescence quenching kinetics for $Zn^{2+}$-TPPS formation………………………………27
UV spectrum for coordinated $Mn^{2+}$-TPPS………………………………….……………….28
UV spectrum for coordinated $Fe^{3+}$-TPPS……………………………….….……………28
UV spectrum for coordinated $Co^{2+}$-TPPS……………………………….……………….29
UV spectrum for coordinated $Ni^{2+}$-TPPS……………………………………………………29
UV spectrum for coordinated $Cu^{2+}$-TPPS……………………….……………………..……30
UV spectrum for coordinated $Zn^{2+}$-TPPS………………………………………………….30
Exponential decay fitting for $Cu^{2+}$-TPPS formation kinetics using fluorescence………..31
Exponential decay fitting for $Zn^{2+}$-TPPS formation kinetics using fluorescence………..32
Exponential decay fitting for $Co^{2+}$-TPPS formation kinetics using fluorescence………..33
Competition experiments for the affinity of different metal ions towards TPPS monitored using UV spectroscopy…………………………………………………………………………… 34
Predicted 3D model of $Co^{2+}$-TPPS using MolView…………………………………………35
Predicted 3D model of $Cu^{2+}$-TPPS using MolView ……………………………………35
Predicted 3D model of $Fe^{3+}$-TPPS.Cl using MolView …………………………………...36
Predicted 3D model of $Mn^{2+}$-TPPS using MolView ………………………………………36
Predicted 3D model of $Ni^{2+}$-TPPS using MolView …………………………………….…37
Predicted 3D model of $Zn^{2+}$-TPPS using MolView ……………………………………..…37
Rayleigh scattering to monitor different metal ion induced aggregate/higher order structure formation…………………………….…………………………………………38

Rayleigh scattering to monitor $Fe^{3+}$ ion mediated aggregate formation in M-TPPS solution ……………………………………………………………………………………38
Differential Interference Contrast (DIC) microscopy analysis of $Fe^{3+}$ ion mediated aggregates in TPPS and $Fe^{3+}$ mixture……………………………………………………….39
Field Emission Scanning Electron Microscopy (FESEM) images for $Fe^{3+}$ ion mediated aggregates in TPPS and $Fe^{3+}$ mixture ……………………………………………………40
Elemental analysis using Electron Dispersive Spectroscopy (EDS) for $Fe^{3+}$ ion mediated aggregates in TPPS and $Fe^{3+}$ mixture ……………………………………………………41
UV characterization of $Fe^{3+}$-TPPS formed upon heating TPPS and $Fe^{3+}$ mixture ……..42
Spontaneous oxidation of NADH monitored by UV……………………………………..…43
$Co^{2+}$ -mediated oxidation of NADH monitored by UV……………………………………...43
TPPS mediated oxidation of NADH monitored by UV……………………………………..44
$Co^{2+}$-TPPS mediated oxidation of NADH monitored by UV……………………………44

TABLES

Reduction potentials of the investigated metal ions and respective oxidation state change……………………………………....………………………………………….…..45
Measured pH of the solution at the initiation (0 hour) and at the end (4 hours) of different oxidation reactions………………………………………………………………………46
Change in the $\lambda_{max}$ in UV absorbance spectrum of co-solute reactions consisting of TPPS and metal ions……………………………………..…………………………...47
High -Resolution Mass Spectrometry (HRMS) analysis of formed M-TPPS……………47

STATISTICAL ANALYSIS

$Cu^{2+}$ mediated oxidation across different time periods for varying atomic ratios……....48

$Cu^{2+}$ mediated oxidation across varying atomic ratios at different time periods………..49

Metal ion mediated oxidation after varying time with respect to the oxidation at 0 hr (at the initiation of the reactions) for 1:2 atomic ratio……………………………………….50

Metal ion mediated oxidation across varying time periods for 1:2 atomic ratio…………50

Metal ion mediated oxidation after four hours with respect to the oxidation at 0 hr (at the initiation of the reactions) for 1:1 atomic ratio……………………………………………50

Metal ion and TPPS co-solute mixture mediated oxidation after four hours with respect to the oxidation at 0 hr (at the initiation of the reactions)…………………………………51

Comparison of oxidation across TPPS and its various co-solute reactions with different metal ions after 4 hrs ……………………………………………………………………52

Comparison of oxidation across different metal ions and their co-solute reactions with TPPS, after 4 hrs………………………………………………………………………………53

Comparison of $Fe^{3+}$-TPPS complex mediated oxidation with the free $Fe^{3+}$ ion and its co-solute reaction with TPPS, after the initiation of the reaction (0 hr) and after 4 hrs…....53

Comparison of M-TPPS mediated oxidation for different metal ions with the respective free metal ion and their co-solute reaction with TPPS at the initiation of the reaction (0 hr) and after 4 hr……………………………………………………………………………54

References………………………………………………………………………………..55

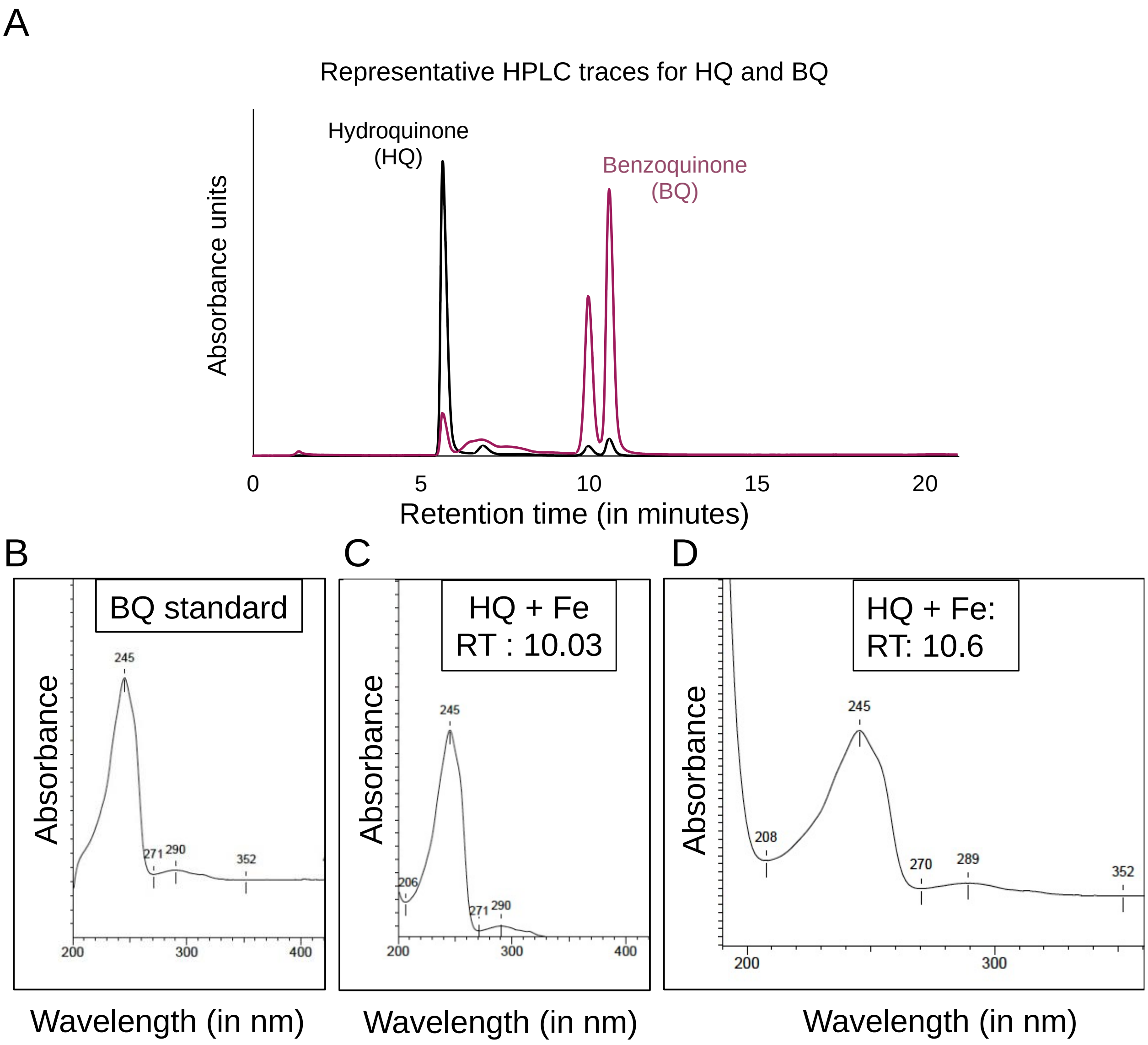

Supplementary Fig. 1: A) Representative High Performance Liquid Chromatography (HPLC) trace showing the elution of Hydroquinone (HQ) at 5.5. minutes and Benzoquinone (BQ) at 10.03 and 10.6 minutes. Black and purple colors show absorption at 288 nm and 244 nm, respectively. Y-axis shows the absorption curve 288 nm (for HQ) and 244 nm (for BQ). X-axis shows the retention time in minutes. Panel B-D show comparison of representative UV absorbance spectrum of BQ standard (B) with two BQ peaks used for quantification i.e., peaks at 10.03 minutes (C) and 10.6 minutes (D) in $Fe^{3+}$-mediated HQ oxidation. Both the peaks show $\lambda_{max}$ at 245 nm corresponding to BQ, which is similar to the spectrum of BQ standard.

**HPLC traces showing oxidation of HQ in presence of $Cu^{2+}$, $Co^{2+}$ and $Fe^{3+}$**

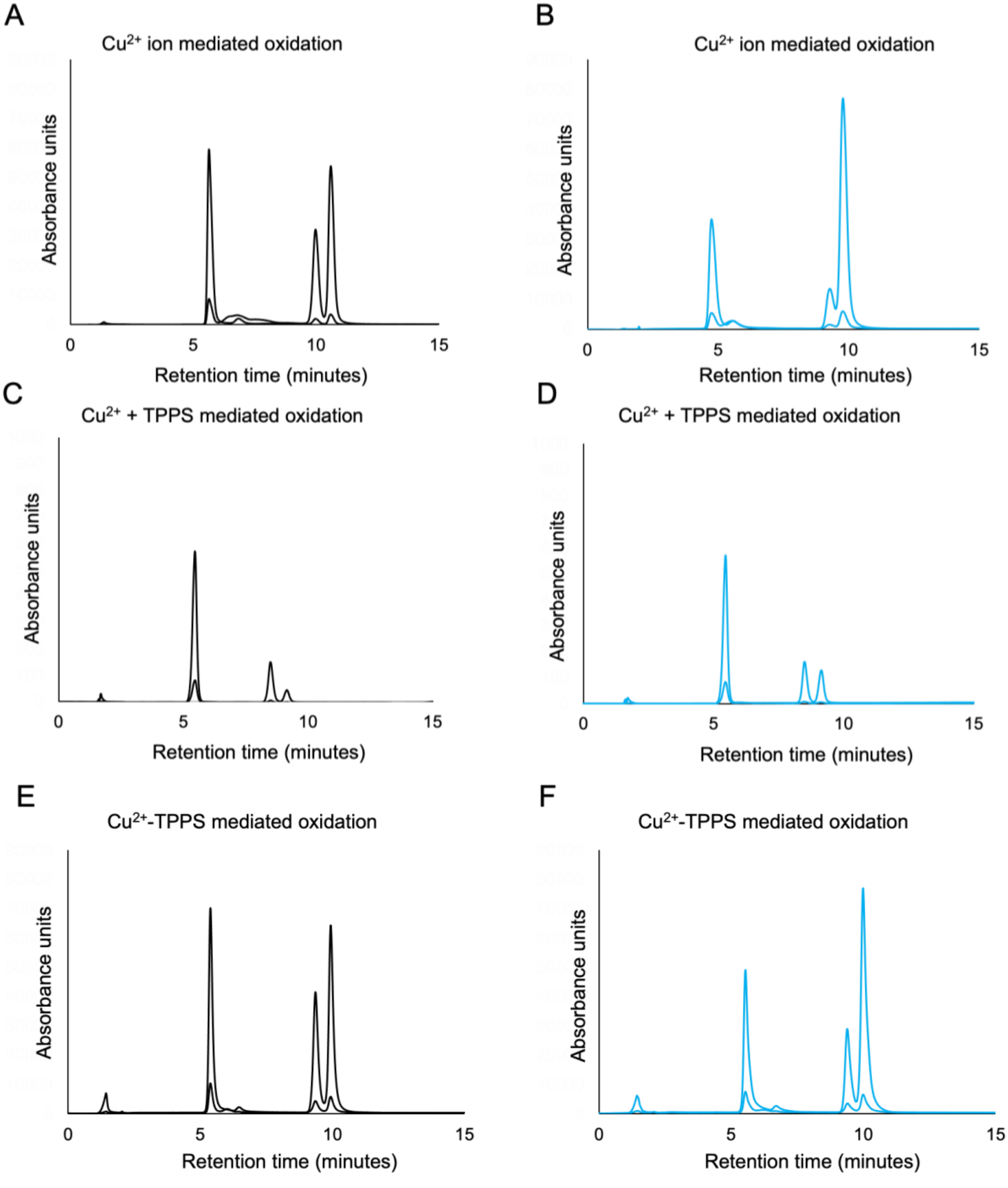

Supplementary Fig. 2: Oxidation of HQ in the presence of $Cu^{2+}$ containing reactions. Panels A and B show the HPLC traces for the oxidation of HQ (5.5. minutes) to BQ (elution time 10.03 and 10.6 minutes), in $Cu^{2+}$ mediated reactions, after the initiation of the reaction (black) and after 4 hours (blue), respectively. Panels C and D show the HQ oxidation in $Cu^{2+}$ and TPPS co-solute reactions, after the initiation of the reaction and after 4 hours, respectively. Panels E and F show the $Cu^{2+}$-TPPS mediated oxidation, after the initiation of the reaction and after 4 hours, respectively.

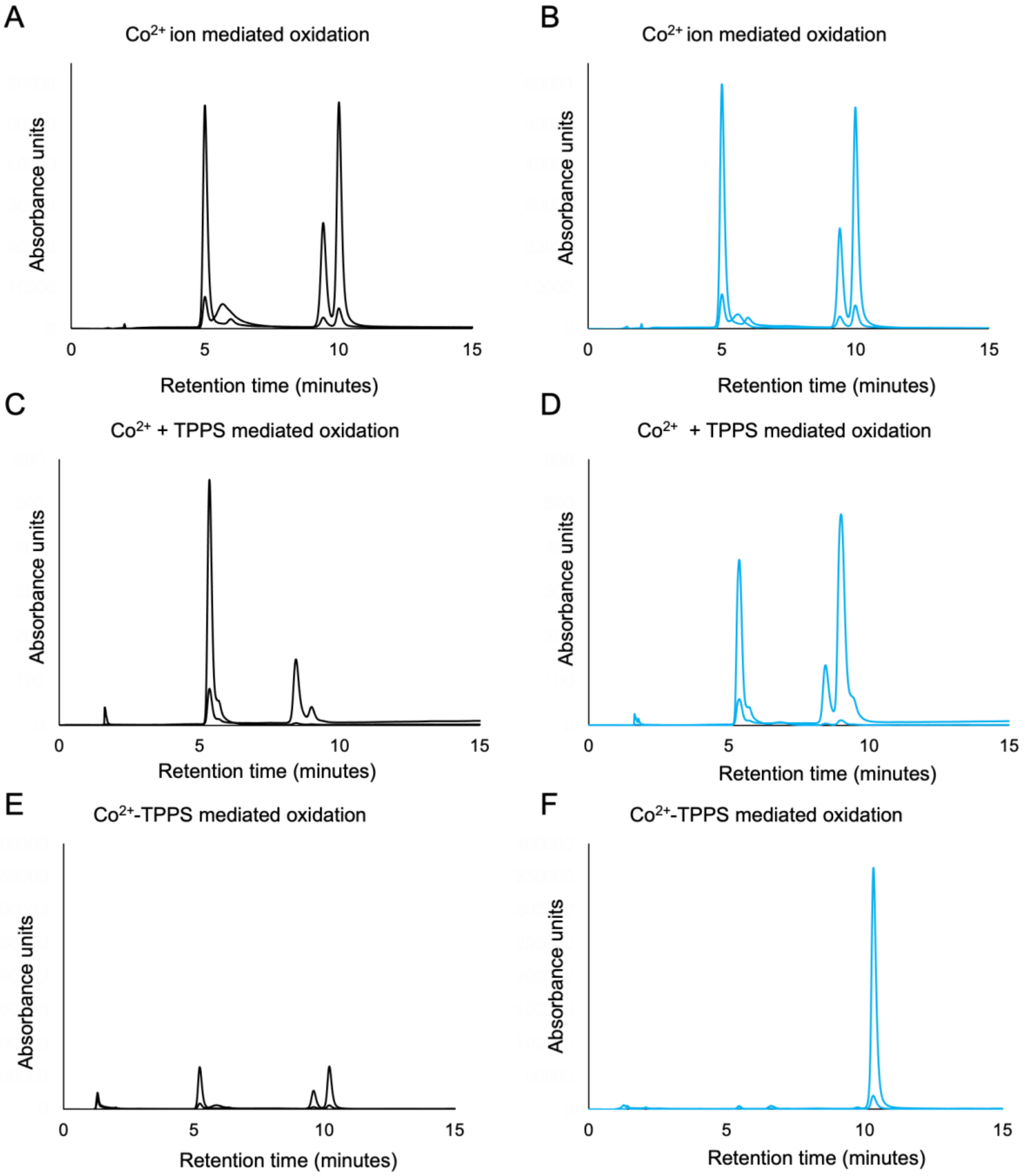

Supplementary Fig. 3: Oxidation of HQ in the presence of $Co^{2+}$ containing reactions. Panels A and B show the HPLC traces for the oxidation of HQ (5.5. minutes) to BQ (elution time 10.03 and 10.6 minutes), in $Co^{2+}$ mediated reactions, after the initiation of the reaction (black) and after 4 hours (blue), respectively. Panels C and D show the HQ oxidation in $Co^{2+}$ and TPPS co-solute reactions, after the initiation of the reaction and after 4 hours, respectively. Panels E and F show the $Co^{2+}$-TPPS mediated oxidation, after the initiation of the reaction and after 4 hours, respectively.

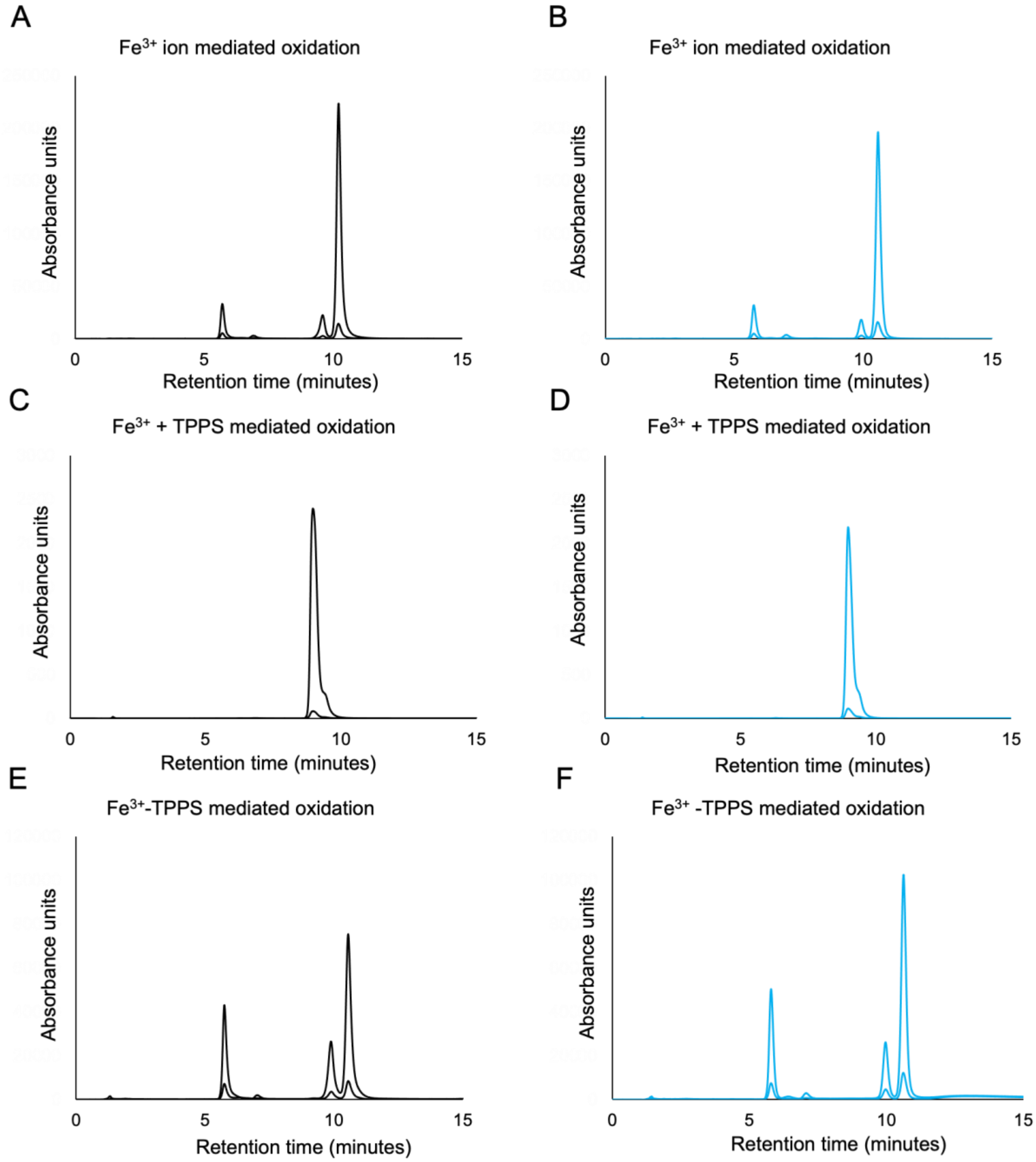

Supplementary Fig. 4: Oxidation of HQ in the presence of $Fe^{3+}$ containing reactions. Panels A and B show the HPLC traces for the oxidation of HQ (5.5. minutes) to BQ (elution time 10.03 and 10.6 minutes), in $Fe^{3+}$ mediated reactions, after the initiation of the reaction (black) and after 4 hours (blue), respectively. Panels C and D show the HQ oxidation in $Fe^{3+}$ and TPPS co-solute reactions, after the initiation of the reaction and after 4 hours, respectively. Panels E and F show the $Fe^{3+}$-TPPS mediated oxidation, after the initiation of the reaction and after 4 hours, respectively.

### HPLC standard curves for quantification of HQ and BQ

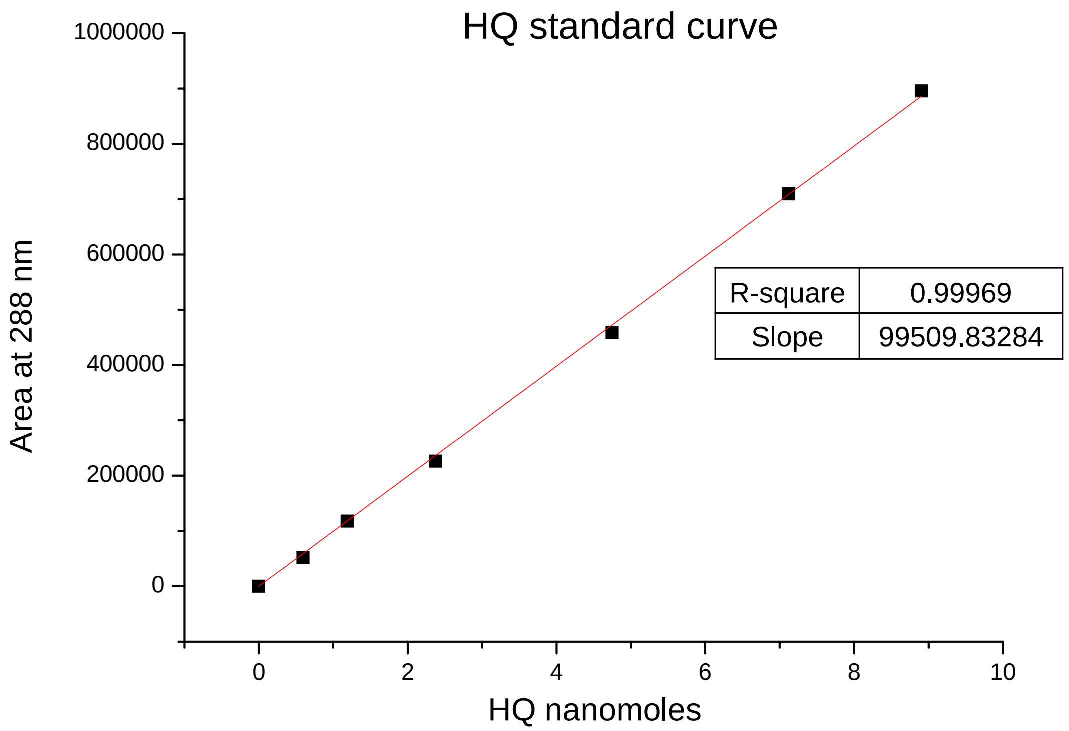

Supplementary Fig. 5: Standard graph for the area under the curve for the HPLC peak corresponding to Hydroquinone (HQ) at 288 nm. Y-axis shows the area under the curve at and absorption of 288 nm. X-axis shows the HQ nanomoles loaded onto the HPLC. Inset shows the R-square value and slope for the linear fitting (shown with red line) of the graph.

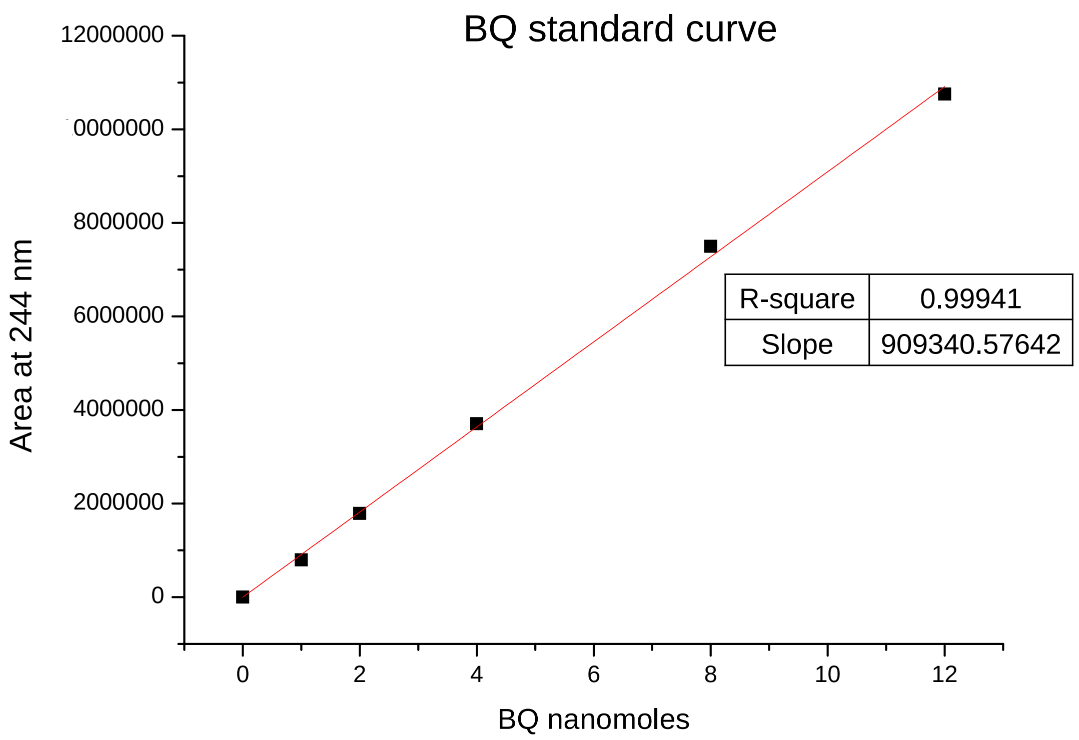

Supplementary Fig. 6: Standard graph for the area under the curve for the HPLC peaks corresponding to Benzoquinone (BQ) at 244 nm. Y-axis shows the area under the curve at and absorption of 244 nm. X-axis shows the BQ nanomoles loaded onto the HPLC. Inset shows the R-square value and slope for the linear fitting (shown with red line) of the graph.

**Oxidation of HQ in presence of different ratios of $Cu^{2+}$ ions**

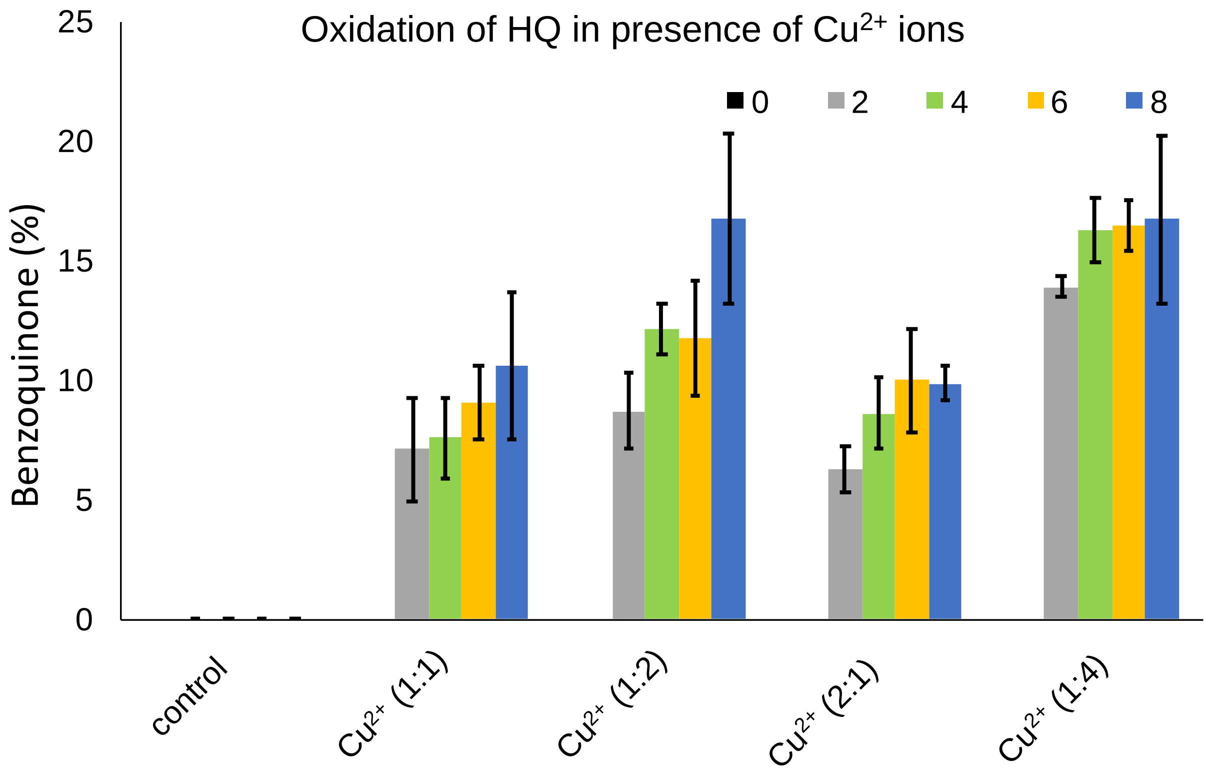

Supplementary Fig. 7: Oxidation of HQ to BQ in presence of $Cu^{2+}$ ions. The graph shows the percentage of benzoquinone (BQ) that is produced from reactions of hydroquinone (HQ), when $CuSO_4$ is used. The variations evaluated include no metal control, and varying ratios of HQ: $Cu^{2+}$ i.e., 1:1 ($Cu^{2+}$ (1:1)), 1:2 ($Cu^{2+}$ (1:2)), 2:1 ($Cu^{2+}$ (2:1)) and 1:4 ($Cu^{2+}$ (1:4)), as depicted on X-axis. Y-axis shows the percentage of benzoquinone produced. Different colors indicate the different time points after which samples were retrieved i.e., immediately after salt addition (0), after 2hrs (2), 4hrs (4), 6hrs (6) and 8hrs (8), respectively, as shown in figure legend. Error bars depict standard deviation; N=3.

**Oxidation of HQ in presence of different metal ions**

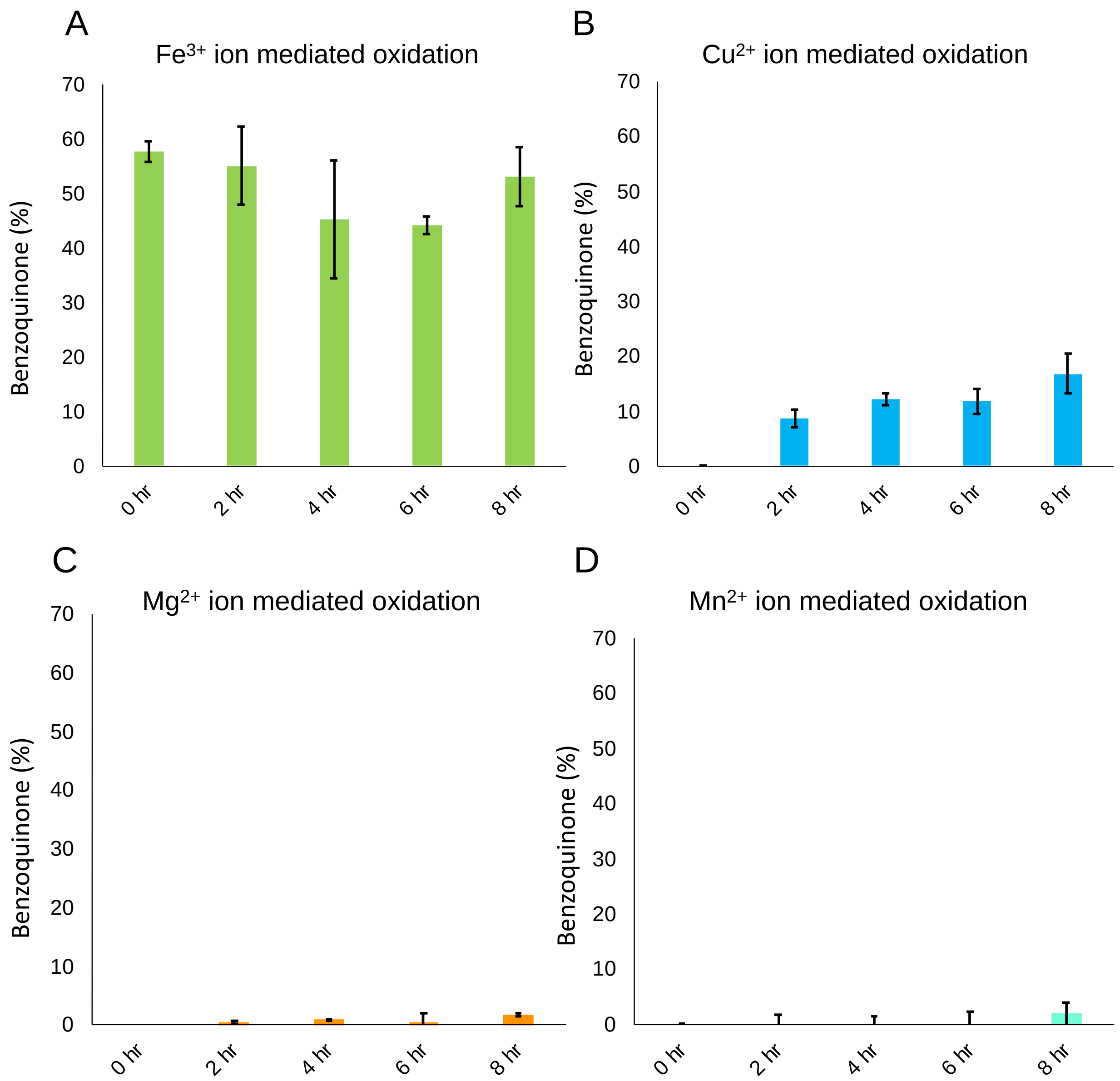

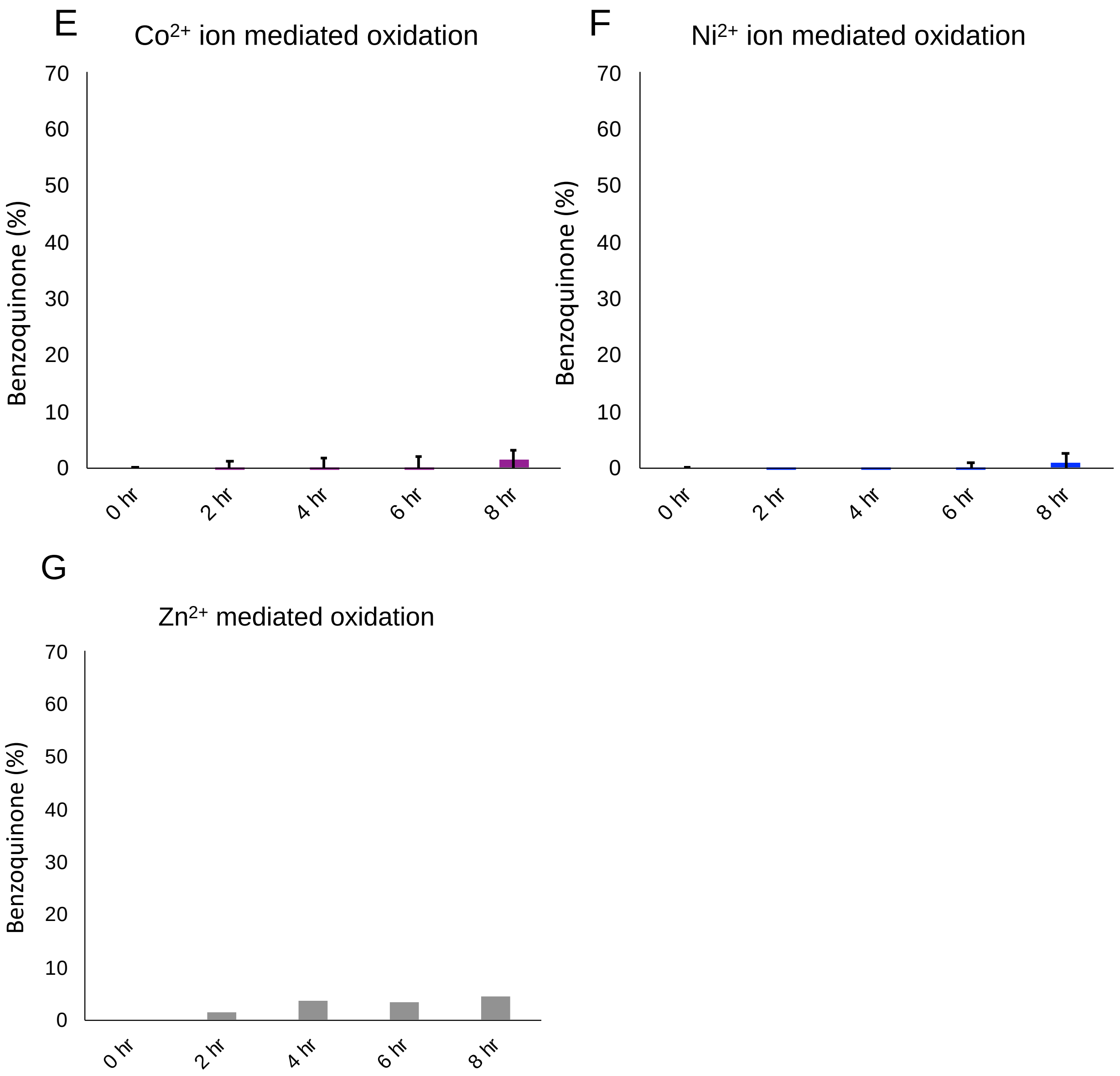

Supplementary Fig. 8: Oxidation of HQ to BQ in the presence of different metal ions. Bars indicate the percentage of benzoquinone (BQ) produced from hydroquinone (HQ) in reactions with different metal ions in 1:2 ratio of HQ to metal ions, at different time points as has been indicated. Panels A to G show reactions involving different metal ions i.e., $Fe^{3+}$, $Cu^{2+}$, $Mg^{2+}$, $Mn^{2+}$, $Co^{2+}$, $Ni^{2+}$ and $Zn^{2+}$, respectively. Y-axis shows the percentage of benzoquinone produced; X-axis depicts the different time points; immediately after the addition of salt (0 hr), after 2hrs (2 hr), 4hrs (4 hr), 6hrs (6 hr) and 8hrs (8 hr). Error bars depict standard deviation; N=2.

**Oxidation of HQ in presence of different ratios of $Cu^{2+}$ and $Fe^{3+}$**

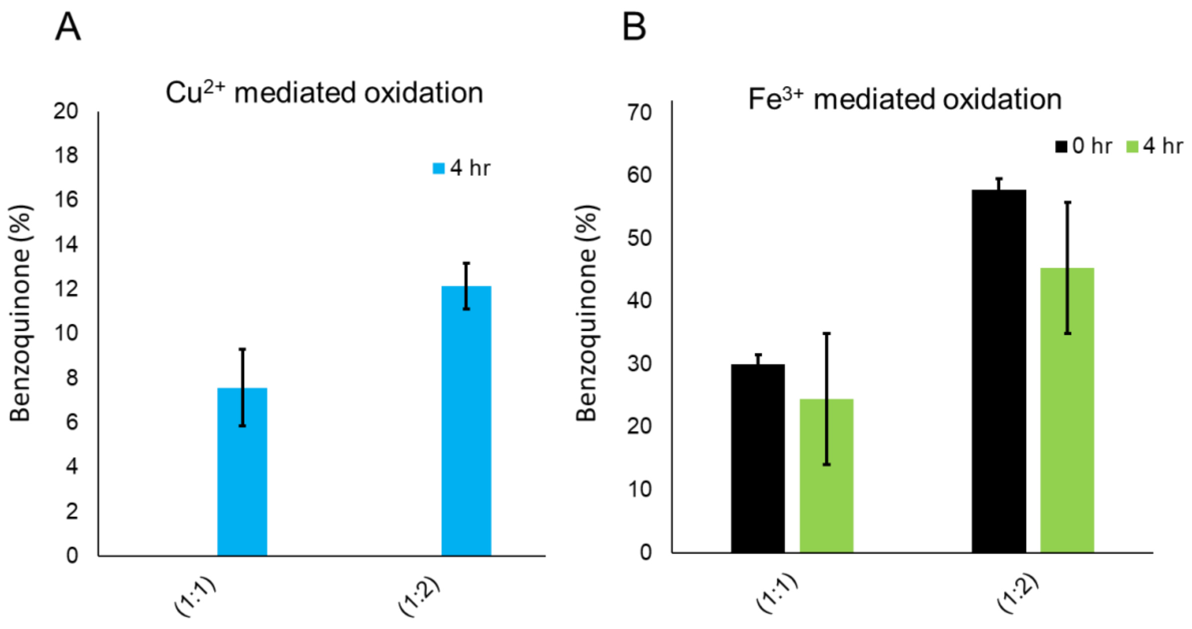

Supplementary Fig. 9: Oxidation of HQ to BQ in presence of different ratios of $Cu^{2+}$ (A) and $Fe^{3+}$ (B). Bars are showing the percentage of benzoquinone (BQ) produced from reactions of hydroquinone (HQ) with: A) $Cu^{2+}$ in 1:1 and 1:2 atomic ratio of HQ to metal ions after four hours (4 hr., blue bar) and B) $Fe^{3+}$ ions in 1:1 and 1:2 ratio of HQ to metal ions immediately after the addition of salt (0 hr., black bar) and after four hours of incubation (4 hr., green bar). Y-axis shows the percentage of benzoquinone produced; Error bars depict standard deviation; N=3.

**UV characterization of tetraphenyl porphyrin tetra sulfonic acid (TPPS) at different pH**

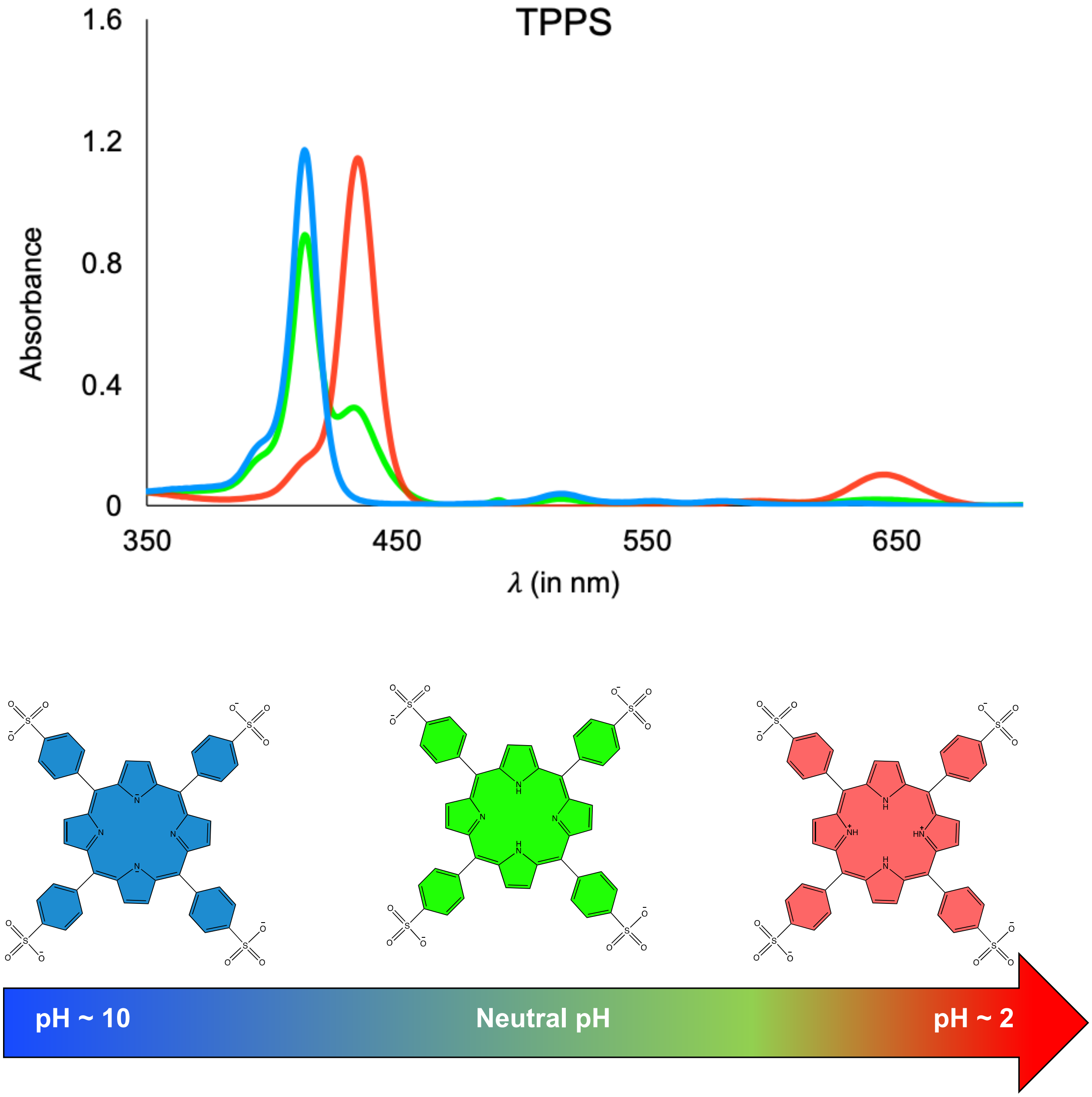

Supplementary Fig. 10: Upper panel indicates the UV spectrum of tetraphenyl porphyrin tetra sulfonic acid (TPPS) at different pH i.e., at pH 2 (red curve), pH 7 (green curve) and pH~10 (blue curve), respectively. X-axis shows the wavelength in nm and Y-axis shows the absorbance. Lower panel shows the different protonation states of TPPS at the aforementioned pH (as indicated by arrow), highlighted in the corresponding color.

**Kinetics of formation of different M-TPPS complexes monitored by UV absorption**

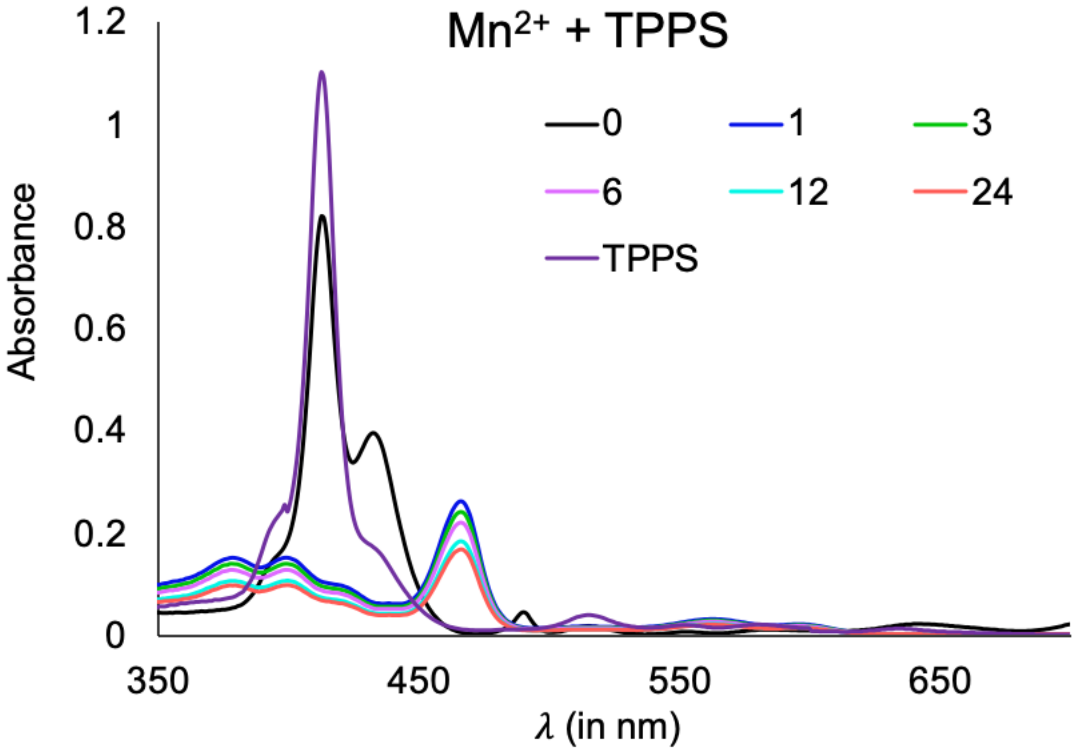

Supplementary Fig. 11: Kinetics of formation of $Mn^{2+}$-TPPS complex monitored by UV absorption. Different colors depict UV spectrum after different time period of incubation at 70ºC (i.e., immediately after the addition of salt (0), after 1hr (1), 3hrs (3), 6hrs (6), 12hrs (12), and 24hrs (24); along with TPPS UV spectrum (TPPS) (purple trace, which corresponds to TPPS spectrum under reaction conditions). Please note that the $\lambda_{max}$ changes to 467 nm. N = 3.

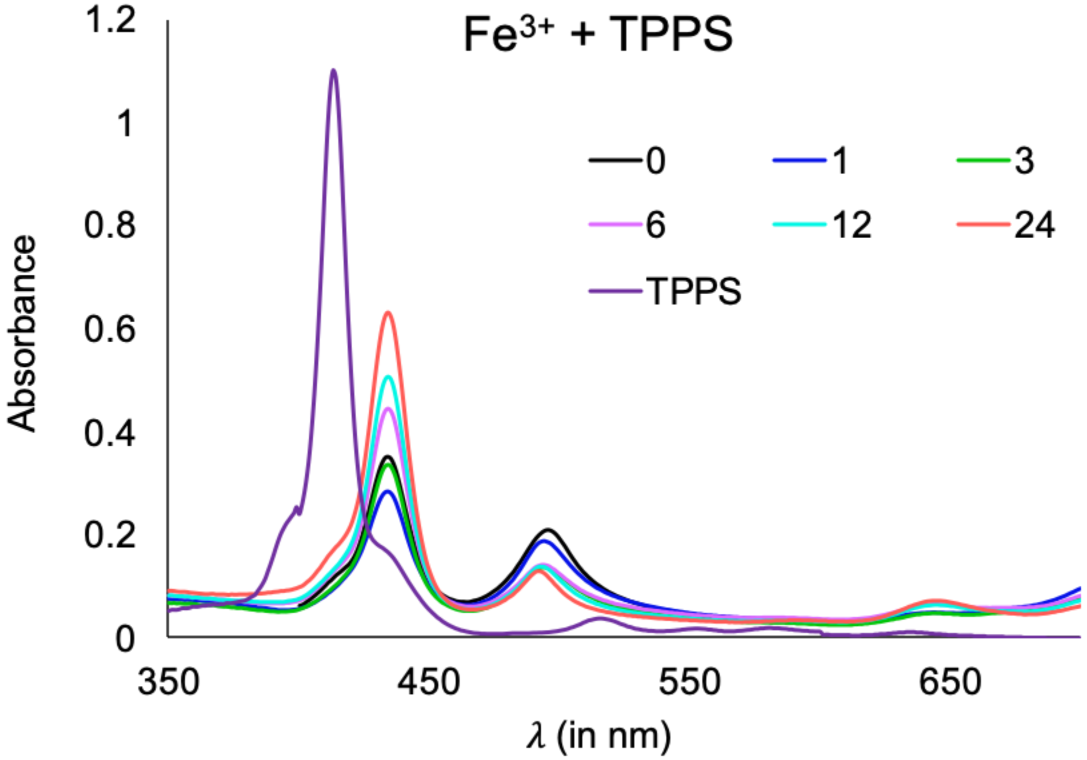

Supplementary Fig. 12: Kinetics of formation of $Fe^{3+}$-TPPS complex monitored by UV absorption. Different colors depict UV spectrum after different time period of incubation at 70ºC (i.e., immediately after the addition of salt (0), after 1hr (1), 3hrs (3), 6hrs (6), 12hrs (12), and 24hrs (24); along with TPPS UV spectrum (TPPS) (purple trace, which corresponds to TPPS spectrum under reaction conditions). Please note that the $\lambda_{max}$ changes to 432 nm and 493 nm. N = 3.

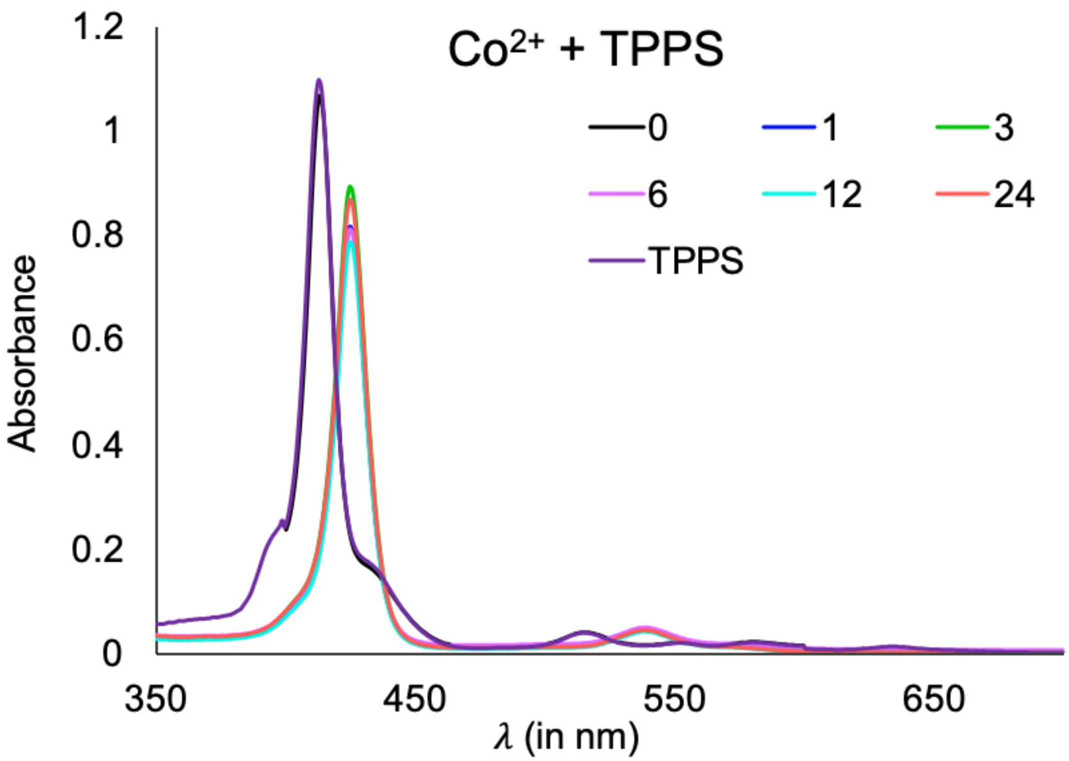

Supplementary Fig. 13: Kinetics of formation of $Co^{2+}$-TPPS complex monitored by UV absorption. Different colors depict UV spectrum after different time period of incubation at 70ºC (i.e., immediately after the addition of salt (0), after 1hr (1), 3hrs (3), 6hrs (6), 12hrs (12), and 24hrs (24); along with TPPS UV spectrum (TPPS) (purple trace, which corresponds to TPPS spectrum under reaction conditions). Please note that the $\lambda_{max}$ changes to 425 nm. N = 3.

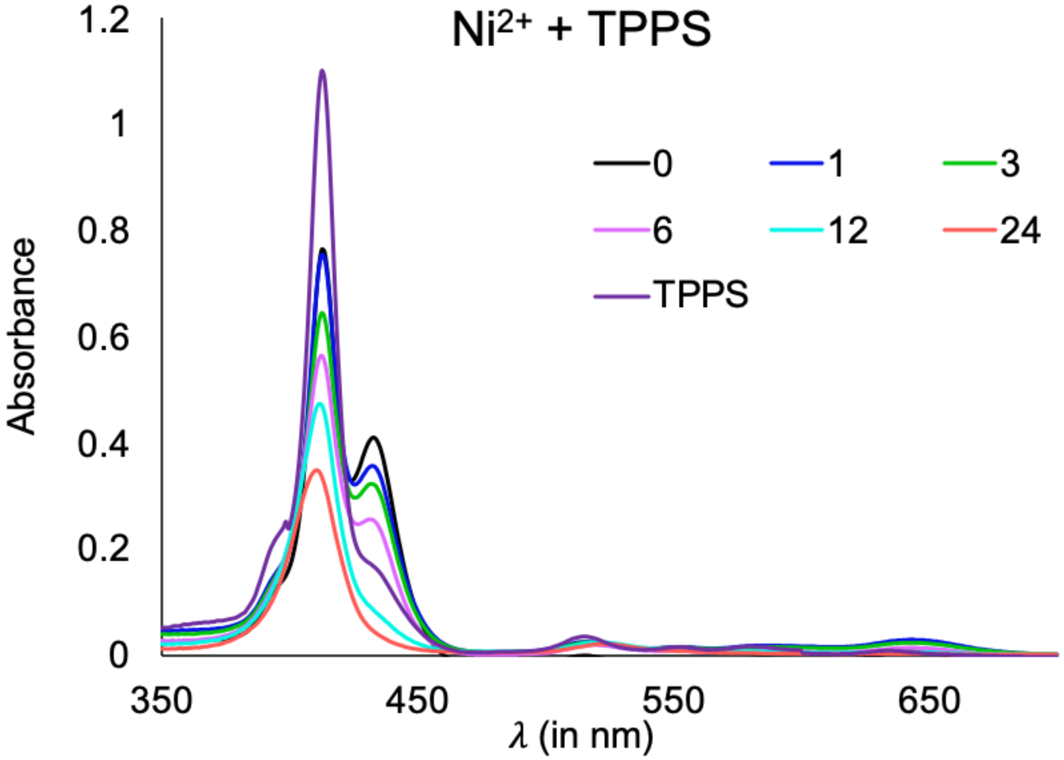

Supplementary Fig. 14: Kinetics of formation of $Ni^{2+}$-TPPS complex monitored by UV absorption. Different colors depict UV spectrum after different time period of incubation at 70ºC (i.e., immediately after the addition of salt (0), after 1hr (1), 3hrs (3), 6hrs (6), 12hrs (12), and 24hrs (24); along with TPPS UV spectrum (TPPS) (purple trace, which corresponds to TPPS spectrum under reaction conditions). Please note that the $\lambda_{max}$ changes to 410 nm. N = 3.

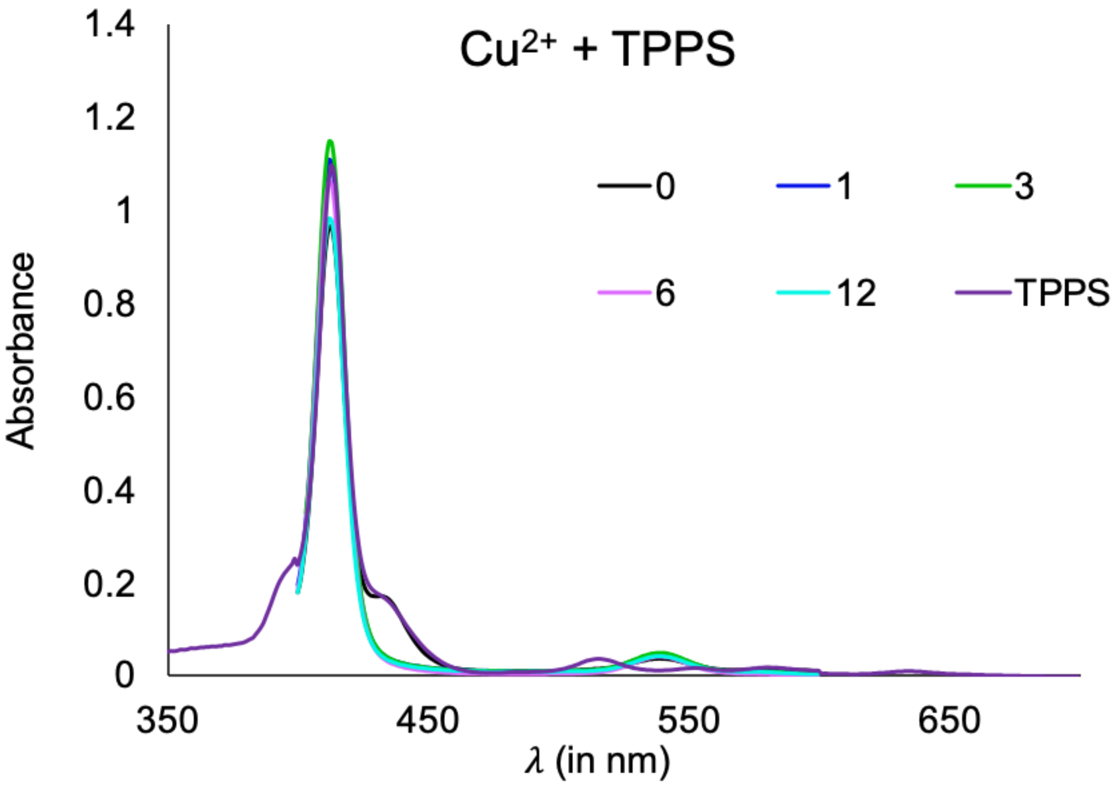

Supplementary Fig. 15 Kinetics of formation of $Cu^{2+}$-TPPS complex monitored by UV absorption. Different colors depict UV spectrum after different time period of incubation at 70ºC (i.e., immediately after the addition of salt (0), after 1hr (1), 3hrs (3), 6hrs (6), 12hrs (12), and 24hrs (24); along with TPPS UV spectrum (TPPS) (purple trace, which corresponds to TPPS spectrum under reaction conditions). Please note that the $\lambda_{max}$ changes to 412 nm. N = 3.

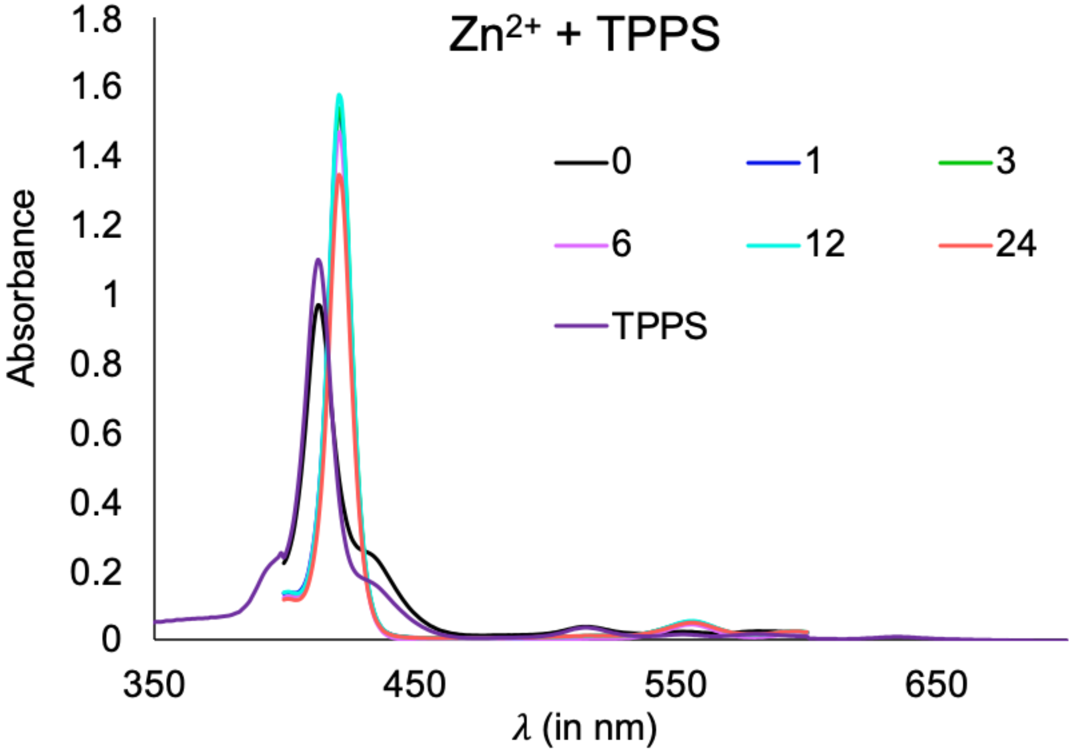

Supplementary Fig. 16: Kinetics of formation of $Zn^{2+}$-TPPS complex monitored by UV absorption. Different colors depict UV spectrum after different time period of incubation at 70ºC (i.e., immediately after the addition of salt (0), after 1hr (1), 3hrs (3), 6hrs (6), 12hrs (12), and 24hrs (24); along with TPPS UV spectrum (TPPS) (purple trace, which corresponds to TPPS spectrum under reaction conditions). Please note that the $\lambda_{max}$ changes to 421 nm. N = 3.

**Kinetics of formation of different M-TPPS complexes monitored using fluorescence**

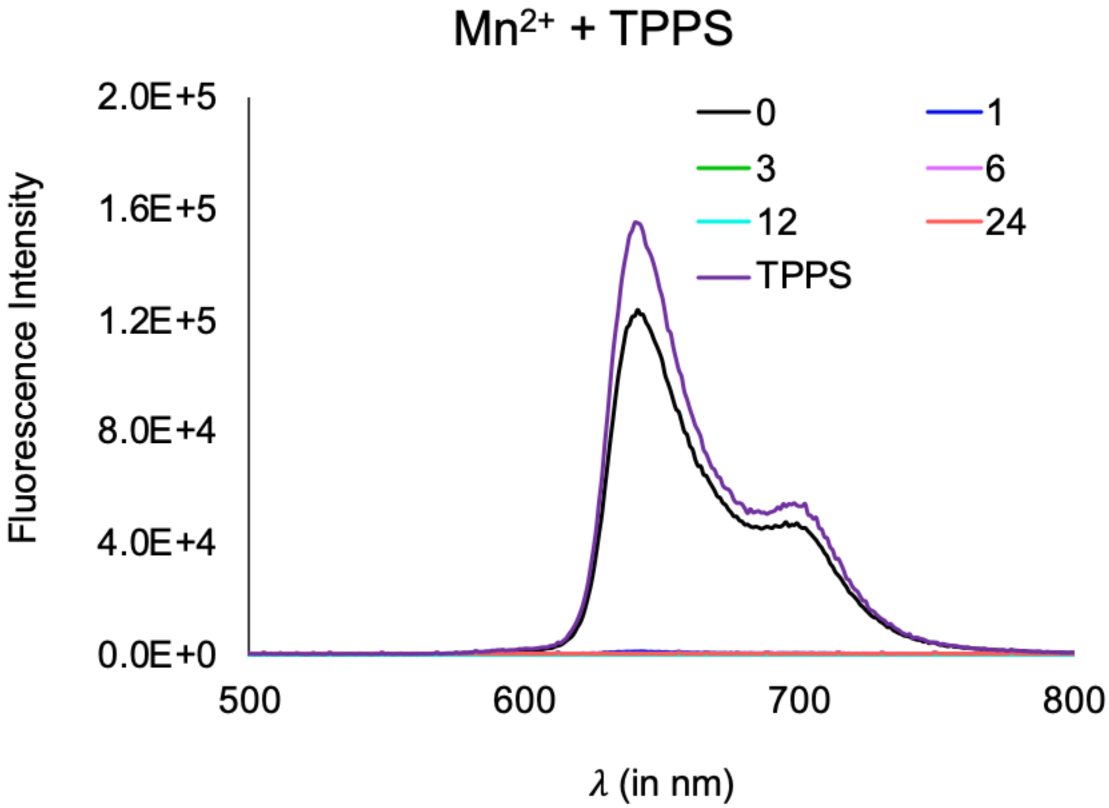

Supplementary Fig. 17: Kinetics of formation of $Mn^{2+}$-TPPS complex monitored by fluorescence emission spectrum by using 414 nm as excitation wavelength. Different colors depict the UV spectrum after different time periods of incubation at 70ºC (i.e., immediately after salt addition (0), after 1hr (1), 3hrs (3), 6hrs (6), 12hrs (12), and 24hrs (24) along with TPPS control spectrum (TPPS; purple trace, which corresponds to TPPS fluorescence under our reaction conditions). Upon metalation, the fluorescence of TPPS gets quenched. N = 3.

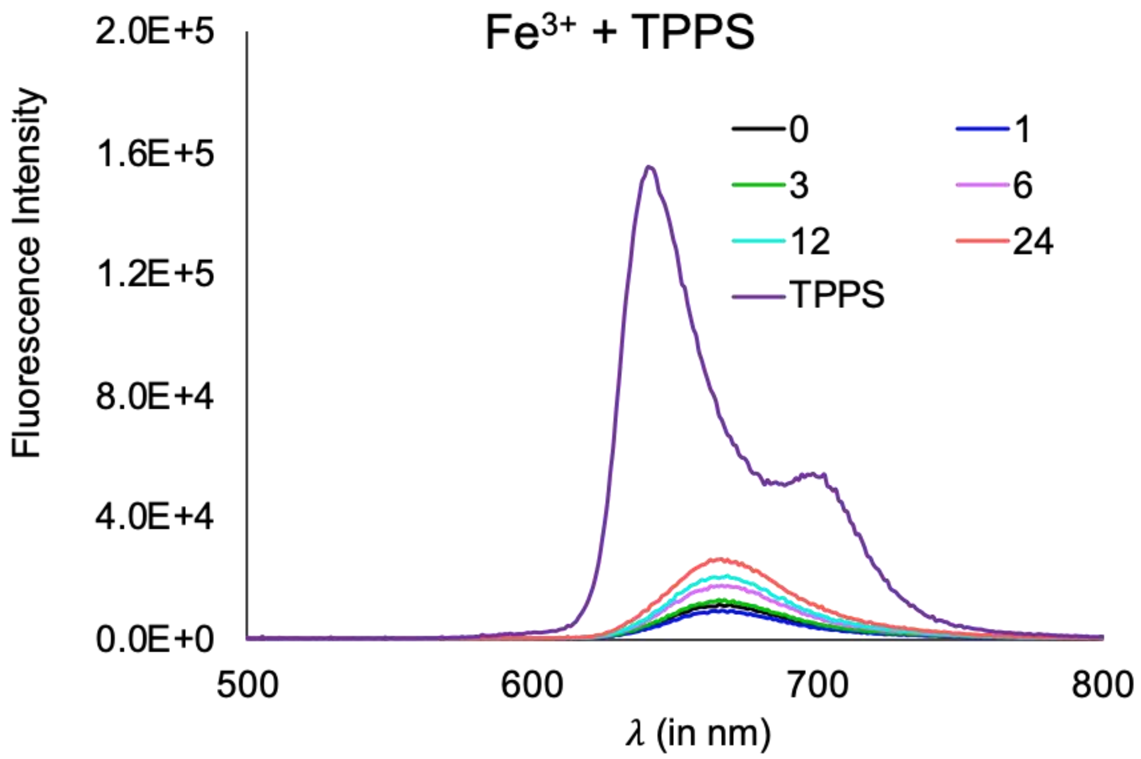

Supplementary Fig. 18: Kinetics of formation of $Fe^{3+}$-TPPS complex monitored by fluorescence emission spectrum by using 414 nm as excitation wavelength. Different colors depict the UV spectrum after different time periods of incubation at 70ºC (i.e., immediately after salt addition (0), after 1hr (1), 3hrs (3), 6hrs (6), 12hrs (12), and 24hrs (24) along with TPPS control spectrum (TPPS; purple trace, which corresponds to TPPS fluorescence under our reaction conditions). Upon metalation, the fluorescence of TPPS gets quenched. N = 3.

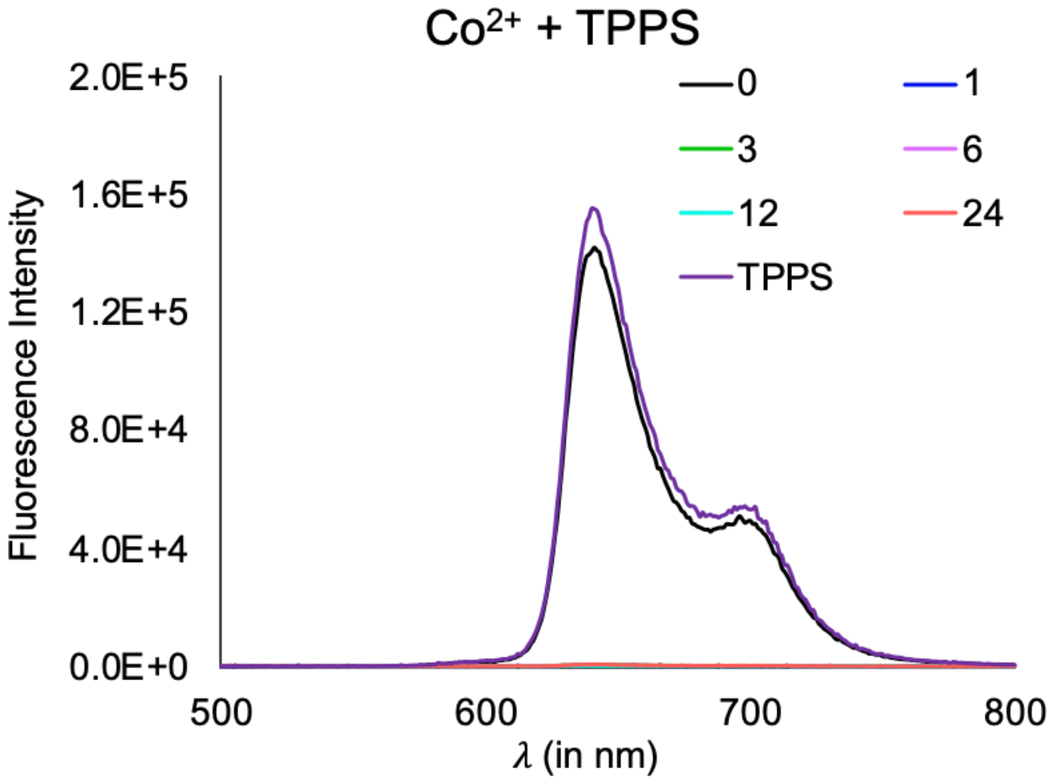

Supplementary Fig. 19: Kinetics of formation of $Co^{2+}$-TPPS complex monitored by fluorescence emission spectrum by using 414 nm as excitation wavelength. Different colors depict the UV spectrum after different time periods of incubation at 70ºC (i.e., immediately after salt addition (0), after 1hr (1), 3hrs (3), 6hrs (6), 12hrs (12), and 24hrs (24) along with TPPS control spectrum (TPPS; purple trace, which corresponds to TPPS fluorescence under our reaction conditions). Upon metalation, the fluorescence of TPPS gets quenched. N = 3.

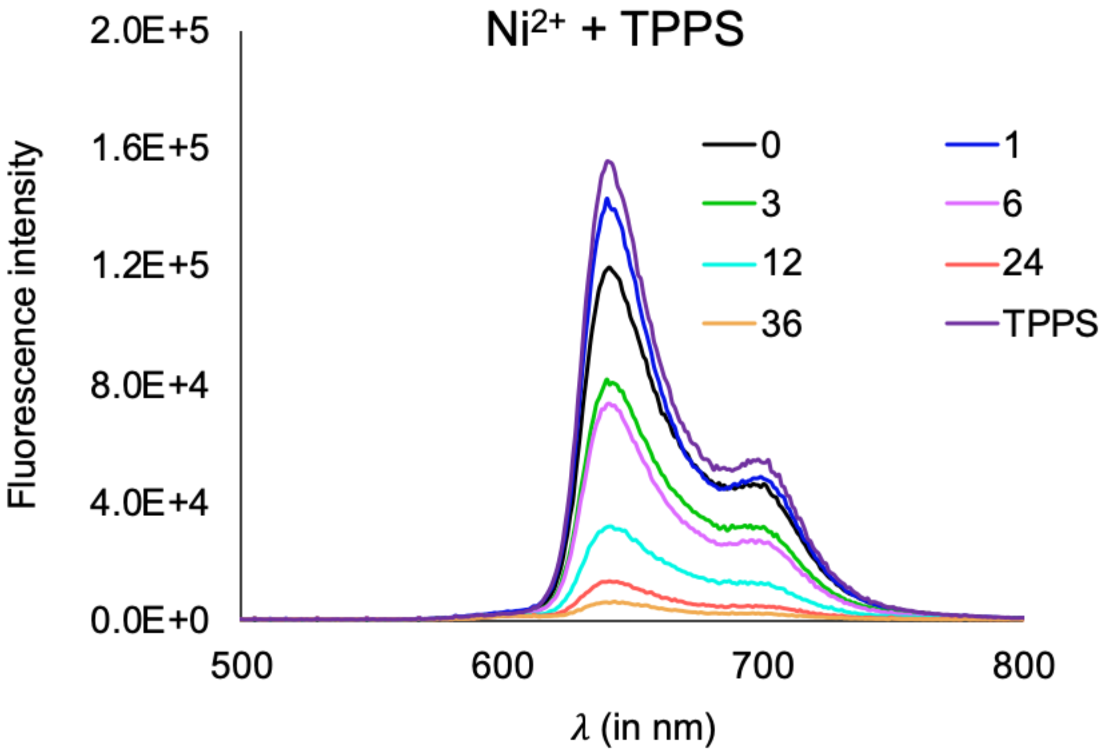

Supplementary Fig. 20: Kinetics of formation of $Ni^{2+}$-TPPS complex monitored by fluorescence emission spectrum by using 414 nm as excitation wavelength. Different colors depict the UV spectrum after different time periods of incubation at 70ºC (i.e., immediately after salt addition (0), after 1hr (1), 3hrs (3), 6hrs (6), 12hrs (12), and 24hrs (24) along with TPPS control spectrum (TPPS; purple trace, which corresponds to TPPS fluorescence under our reaction conditions). Upon metalation, the fluorescence of TPPS gets quenched. N = 3.

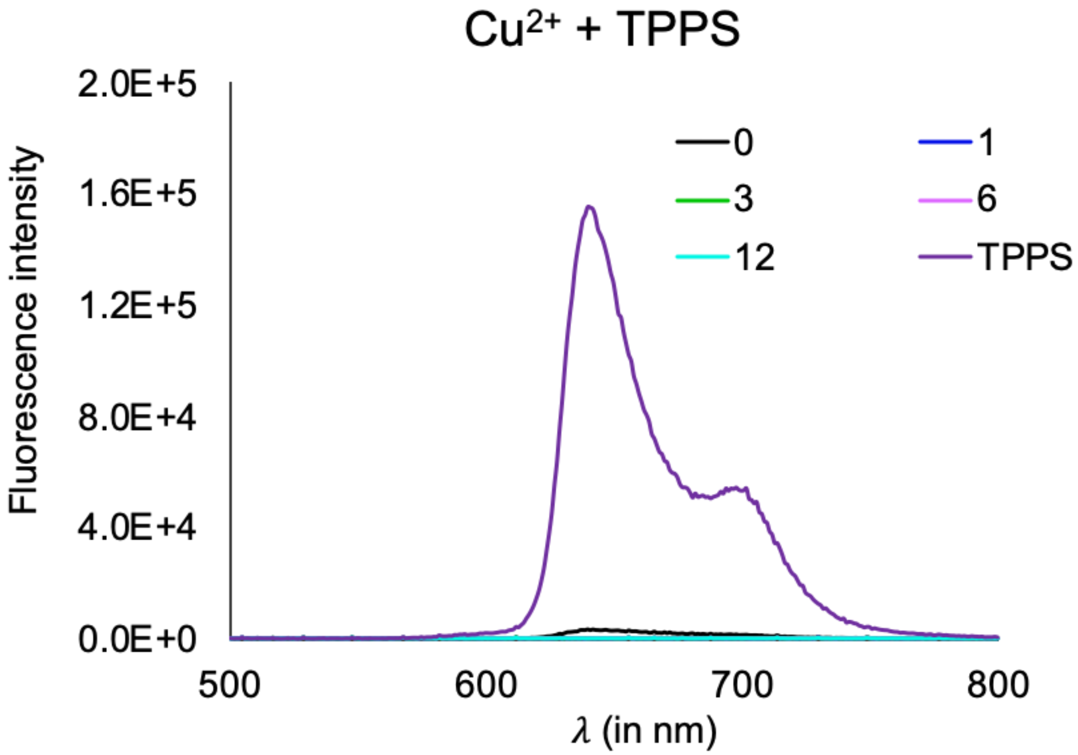

Supplementary Fig. 21: Kinetics of formation of $Cu^{2+}$-TPPS complex monitored by fluorescence emission spectrum by using 414 nm as excitation wavelength. Different colors depict the UV spectrum after different time periods of incubation at 70ºC (i.e., immediately after salt addition (0), after 1hr (1), 3hrs (3), 6hrs (6), 12hrs (12), and 24hrs (24) along with TPPS control spectrum (TPPS; purple trace, which corresponds to TPPS fluorescence under our reaction conditions). Upon metalation, the fluorescence of TPPS gets quenched. N = 3.

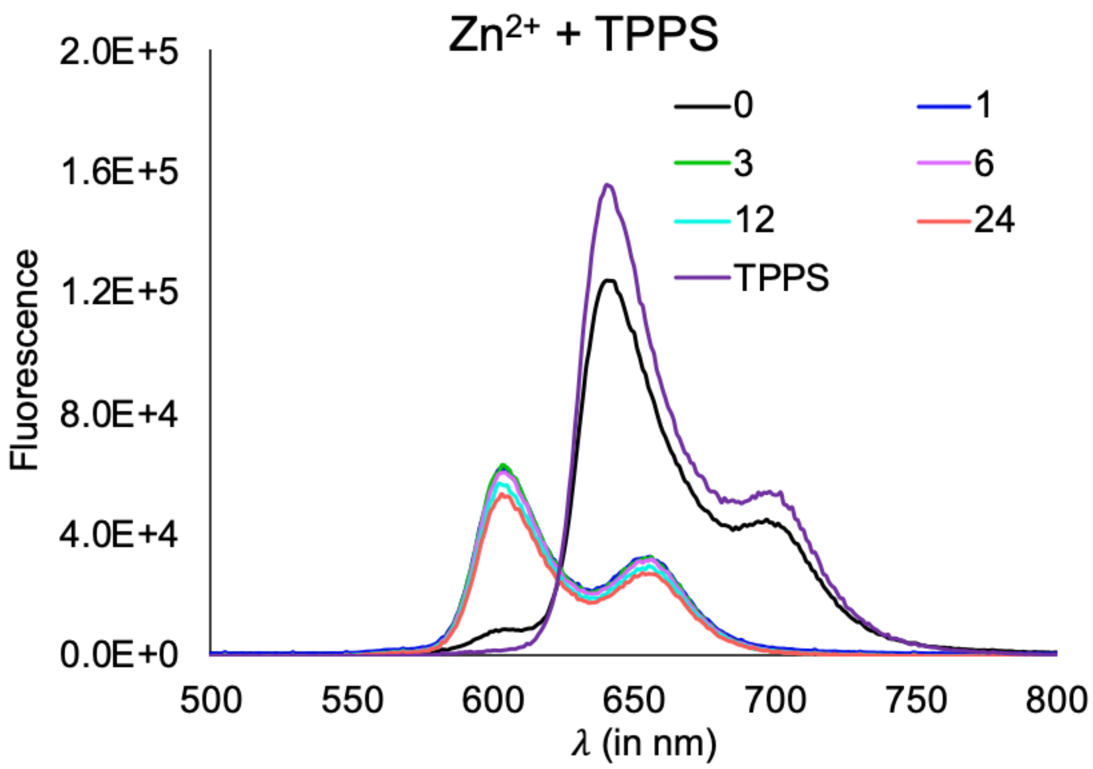

Supplementary Fig. 22: Kinetics of formation of $Zn^{2+}$-TPPS complex monitored by fluorescence emission spectrum by using 414 nm as excitation wavelength. Different colors depict the UV spectrum after different time periods of incubation at 70ºC (i.e., immediately after salt addition (0), after 1hr (1), 3hrs (3), 6hrs (6), 12hrs (12), and 24hrs (24) along with TPPS control spectrum (TPPS; purple trace, which corresponds to TPPS fluorescence under our reaction conditions). In the case of $Zn^{2+}$, upon metalation, a shift in $\lambda_{emission}$ to 603 nm and 655 nm was observed. N = 3.

**UV absorption of different coordinated M-TPPS complexes**

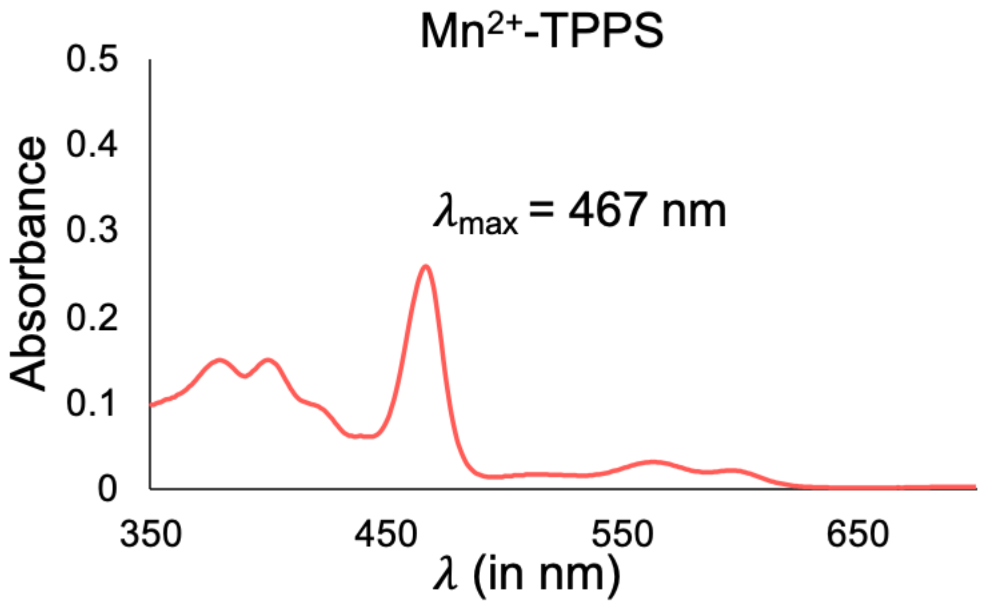

Supplementary Fig. 23: UV spectrum of preformed $Mn^{2+}$-TPPS complex, which is characterized by a strong band with λmax at 467 nm.

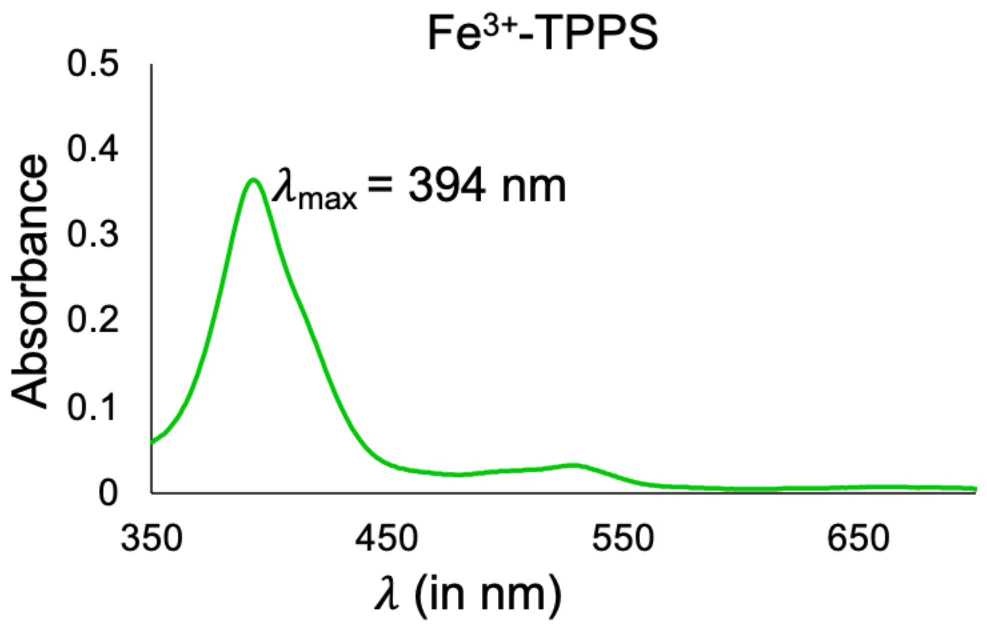

Supplementary Fig. 24: UV spectrum of commercially acquired $Fe^{3+}$-TPPS.Cl complex, which is characterized by a strong band with λmax at 394 nm.

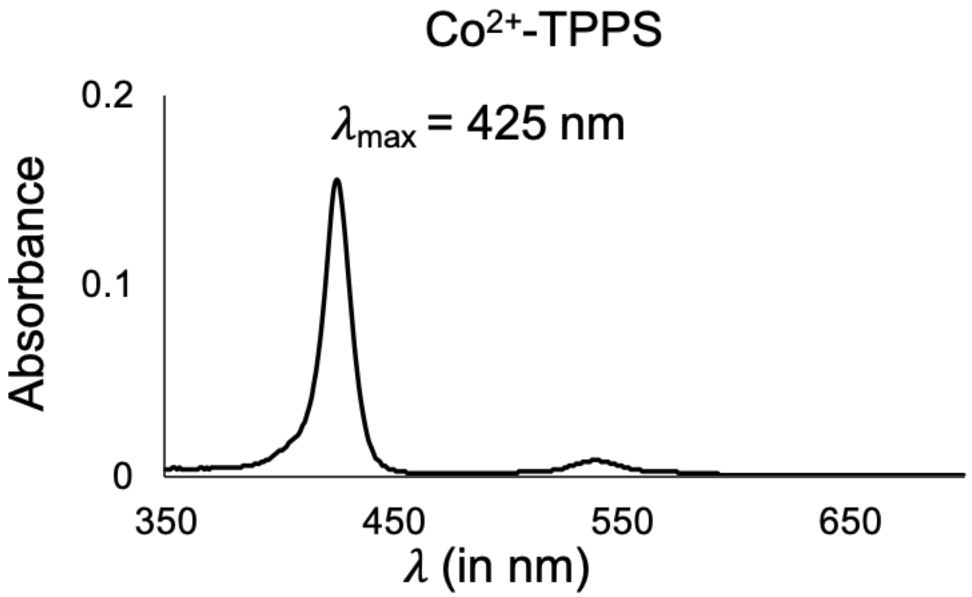

Supplementary Fig. 25: UV spectrum of preformed $Co^{2+}$-TPPS complex, which is characterized by a strong band with λmax at 425 nm.

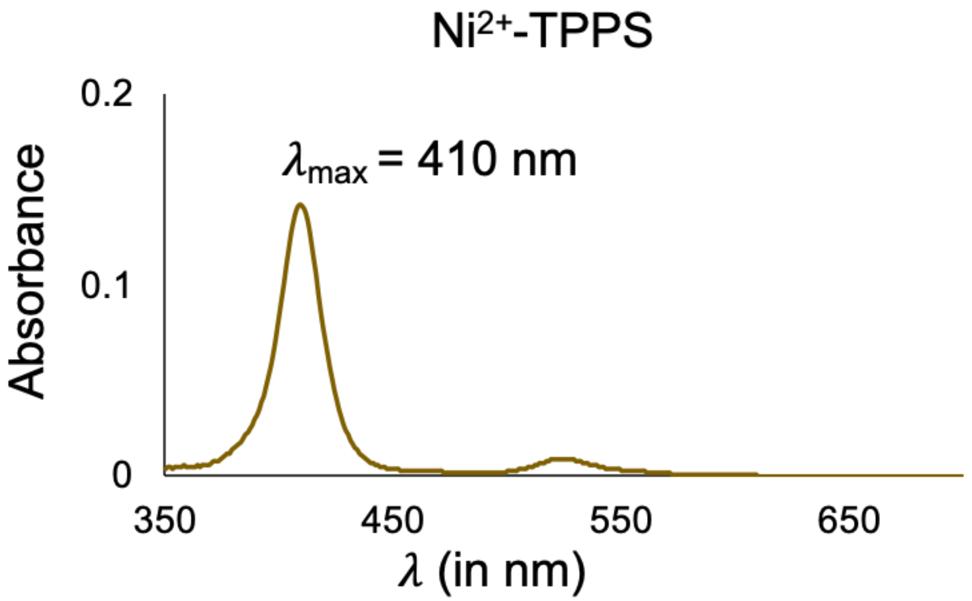

Supplementary Fig. 26: UV spectrum of preformed $Ni^{2+}$-TPPS complex, which is characterized by a strong band with λmax at 410 nm.

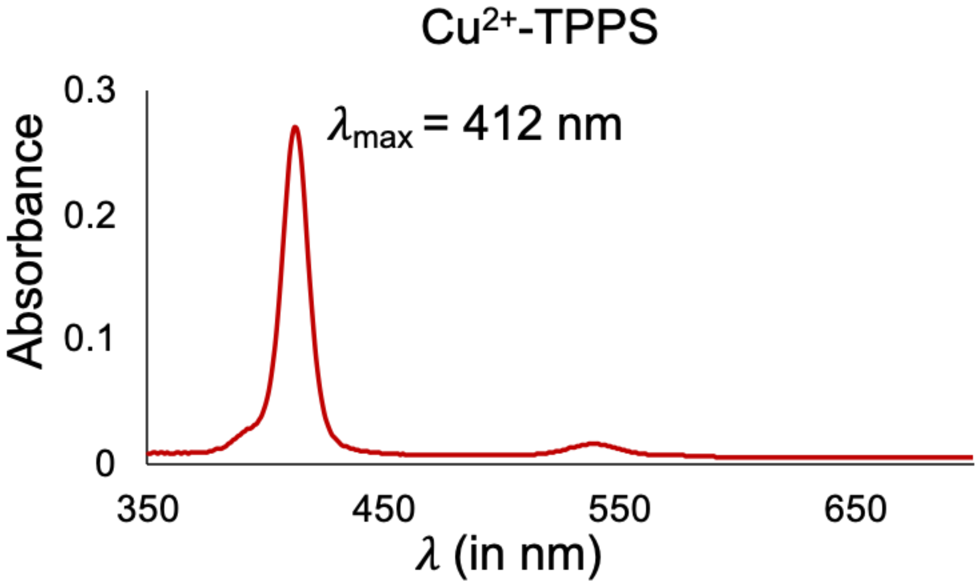

Supplementary Fig. 27: UV spectrum of preformed $Cu^{2+}$-TPPS complex, which is characterized by a strong band with λmax at 412 nm.

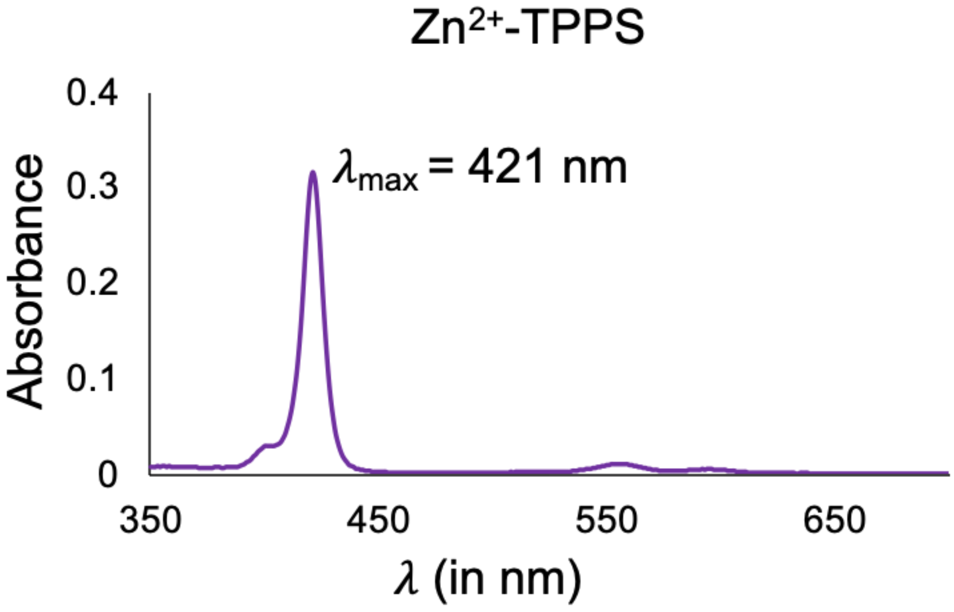

Supplementary Fig. 28: UV spectrum of preformed $Zn^{2+}$-TPPS complex, which is characterized by a strong band with λmax at 421 nm.

### Fast-kinetic measurements using fluorescence quenching at 693 nm for M-TPPS formation

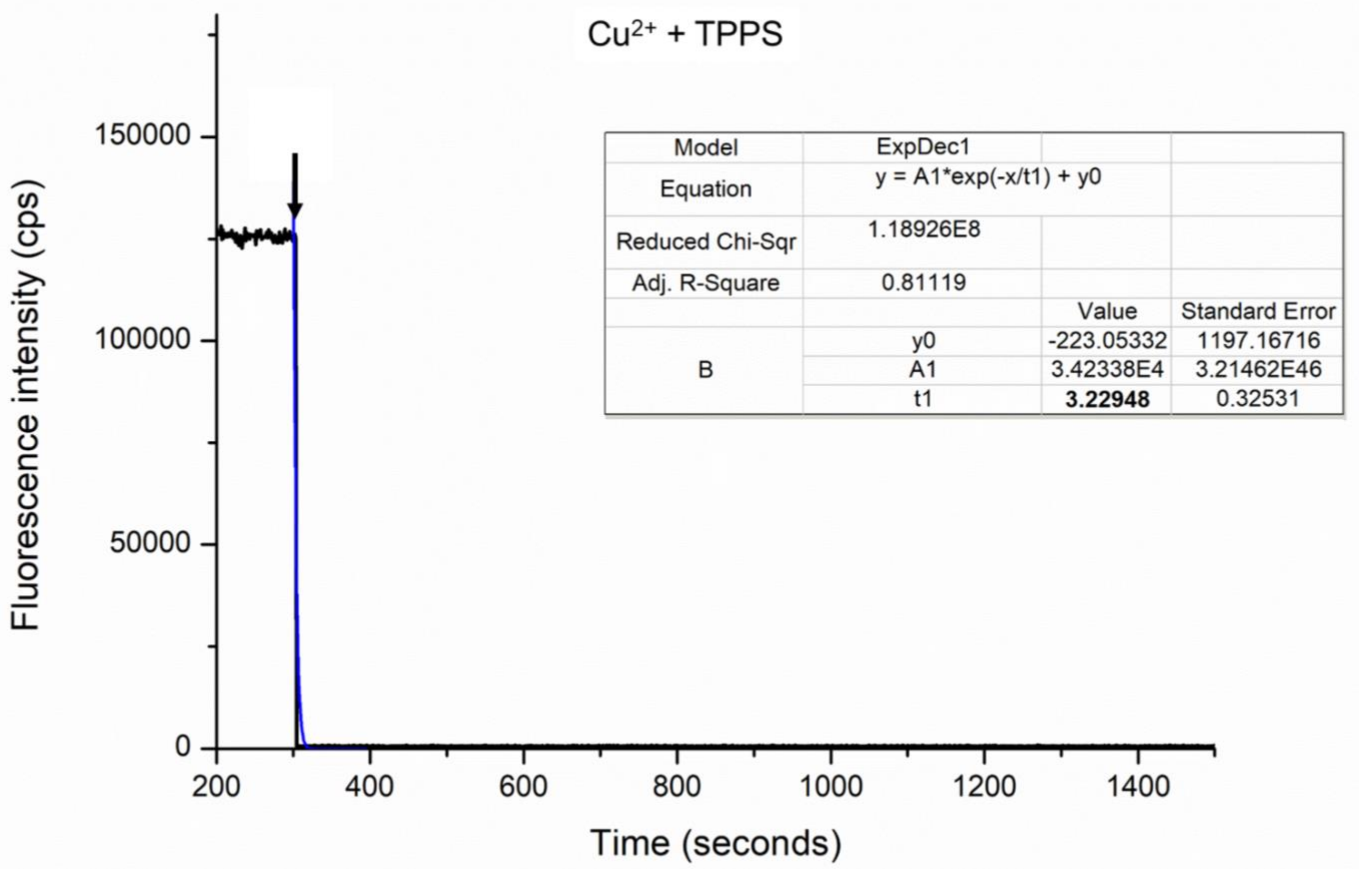

| Model | ExpDec1 | | |
|---|---|---|---|
| Equation | y = A1*exp(-x/t1) + y0 | | |
| Reduced Chi-Sqr | 1.18926E8 | | |
| Adj. R-Square | 0.81119 | | |
| | | Value | Standard Error |
| B | y0 | -223.05332 | 1197.16716 |
| | A1 | 3.42338E4 | 3.21462E46 |
| | t1 | **3.22948** | 0.32531 |

Supplementary Fig. 29: The fast-kinetics for the fluorescence quenching at 693 nm in the case of solution containing $Cu^{2+}$ (black curve). The dark blue line shows the exponential decay fitting of the data. Inset shows the equation for the observed decay, the R-square value and other calculated parameters. The value of time constant t1 for the fluorescence quenching is highlighted in bold, which is indicative of metal-TPPS complex formation. Black arrow shows the time point corresponding to the addition of the $Cu^{2+}$ to the TPPS solution (this was done after 300 seconds). Y-axis shows the fluorescence intensity in cps (counts per second) and X-axis shows the time in seconds. N = 3.

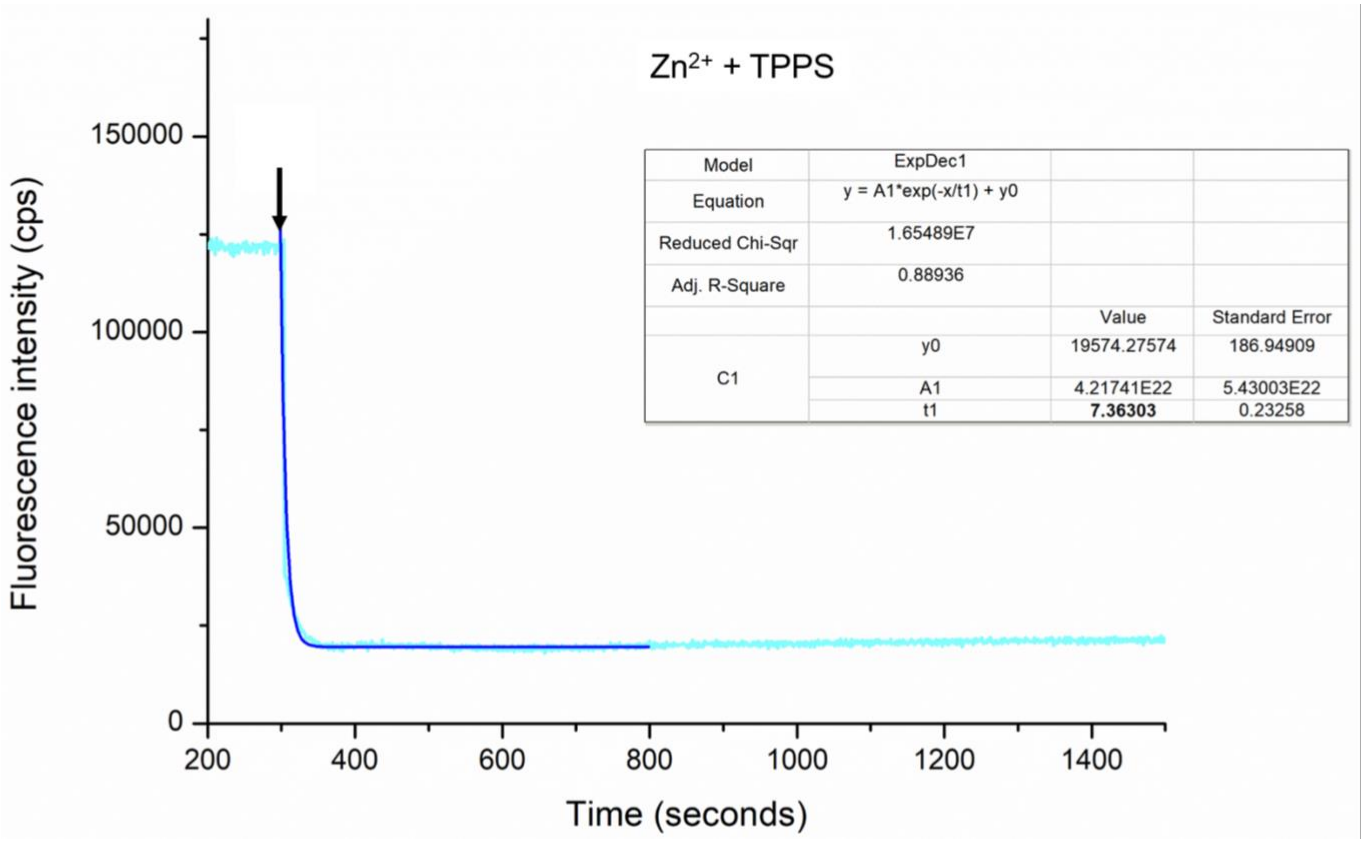

Supplementary Fig. 30: The fast-kinetics for the fluorescence quenching at 693 nm in the case of solutions containing $Zn^{2+}$ (cyan curve). The dark blue line shows the exponential decay fitting of the data. Inset shows the equation for the observed decay, the R-square value and other calculated parameters. The value of time constant t1 for the fluorescence quenching is highlighted in bold, which is indicative of metal-TPPS complex formation. Black arrow shows the time point corresponding to the addition of the $Zn^{2+}$ to the TPPS solution (this was done after 300 seconds). Y-axis shows the fluorescence intensity in cps (counts per second) and X-axis shows the time in seconds. N = 3.

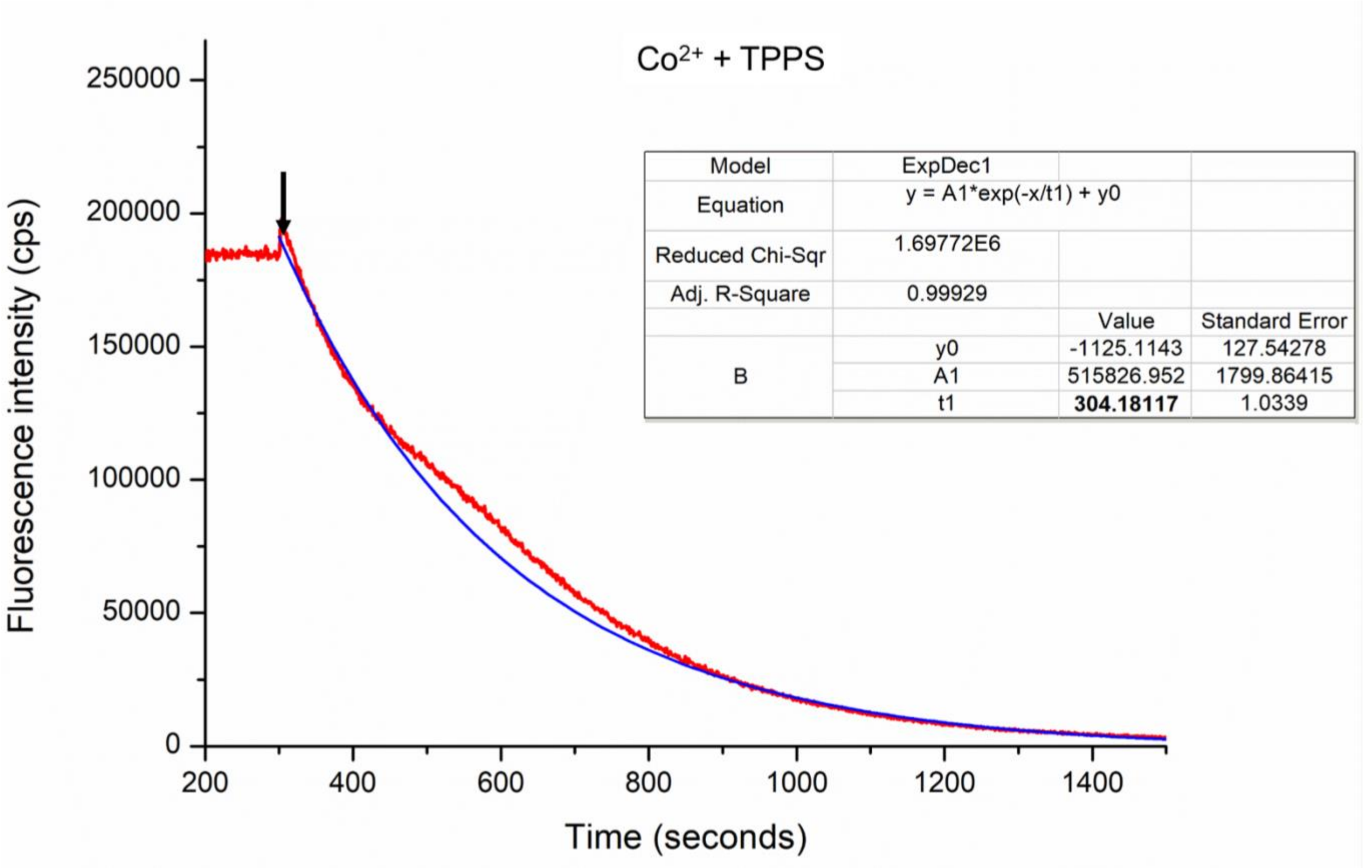

Supplementary Fig. 31: The fast-kinetics for the fluorescence quenching at 693 nm in the case of solutions containing $Co^{2+}$ (red curve). The dark blue line shows the exponential decay fitting of the data. Inset shows the equation for the observed decay, the R-square value and other calculated parameters. The value of time constant t1 for the fluorescence quenching is highlighted in bold, which is indicative of metal-TPPS complex formation. Black arrow shows the time point corresponding to the addition of the $Co^{2+}$ to the TPPS solution (this was done after 300 seconds). Y-axis shows the fluorescence intensity in cps (counts per second) and X-axis shows the time in seconds. N = 3.

**Competition experiments for the affinity of different metal ions towards TPPS:**

**Different set indicates different composition of metal ions:**

**Set 1:** $Mn^{2+}$, $Fe^{3+}$, $Co^{2+}$, $Ni^{2+}$, $Cu^{2+}$ and $Zn^{2+}$
**Set 2:** $Mn^{2+}$, $Fe^{3+}$, $Co^{2+}$, $Ni^{2+}$ and $Zn^{2+}$
**Set3:** $Mn^{2+}$, $Fe^{3+}$, $Co^{2+}$ and $Ni^{2+}$
**Set 4:** $Mn^{2+}$, $Fe^{3+}$ and $Ni^{2+}$

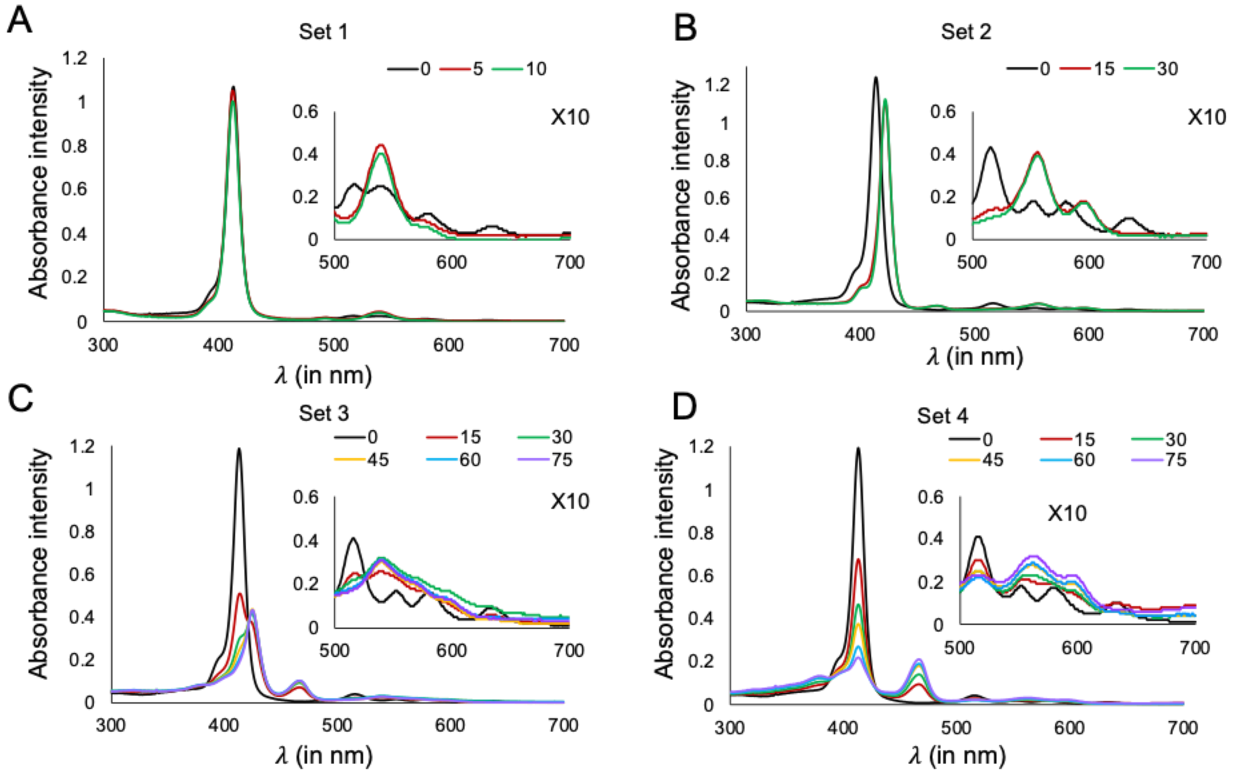

Supplementary Fig. 32: Competition experiments for the affinity of different metal ions towards TPPS, monitored using UV spectroscopy at varying time periods as depicted by different colors (in minutes). Different sets indicate mixtures of different metal ions. Legend indicates varying time periods (in minutes). A) Set 1 shows the preferential formation of $Cu^{2+}$-TPPS ($\lambda_{max}$ = 412 nm) when TPPS was incubated with a mixture of $Mn^{2+}$, $Fe^{3+}$, $Co^{2+}$, $Ni^{2+}$, $Cu^{2+}$ and $Zn^{2+}$. B) Set 2 shows the preferential formation of $Zn^{2+}$-TPPS ($\lambda_{max}$ = 421 nm) when TPPS was incubated with a mixture of $Mn^{2+}$, $Fe^{3+}$, $Co^{2+}$, $Ni^{2+}$ and $Zn^{2+}$(Set 1 without $Cu^{2+}$). C) Set 3 shows the preferential formation of $Co^{2+}$-TPPS ($\lambda_{max}$ = 427 nm) when TPPS was incubated with a mixture of $Mn^{2+}$, $Fe^{3+}$, $Co^{2+}$ and $Ni^{2+}$ (Set 2 without $Zn^{2+}$). D) Set 4 shows the preferential formation of $Mn^{2+}$-TPPS ($\lambda_{max}$ = 467 nm) when TPPS was incubated with a mixture of $Mn^{2+}$, $Fe^{3+}$ and $Ni^{2+}$ (Set 3 without $Co^{2+}$). Inset in all these spectrum shows the 10X zoomed spectra in the region of 500-700 nm (region of Q-bands).

**Predicted 3D model of M-TPPS complex using energy minimization simulations in MolView (molview.org)**

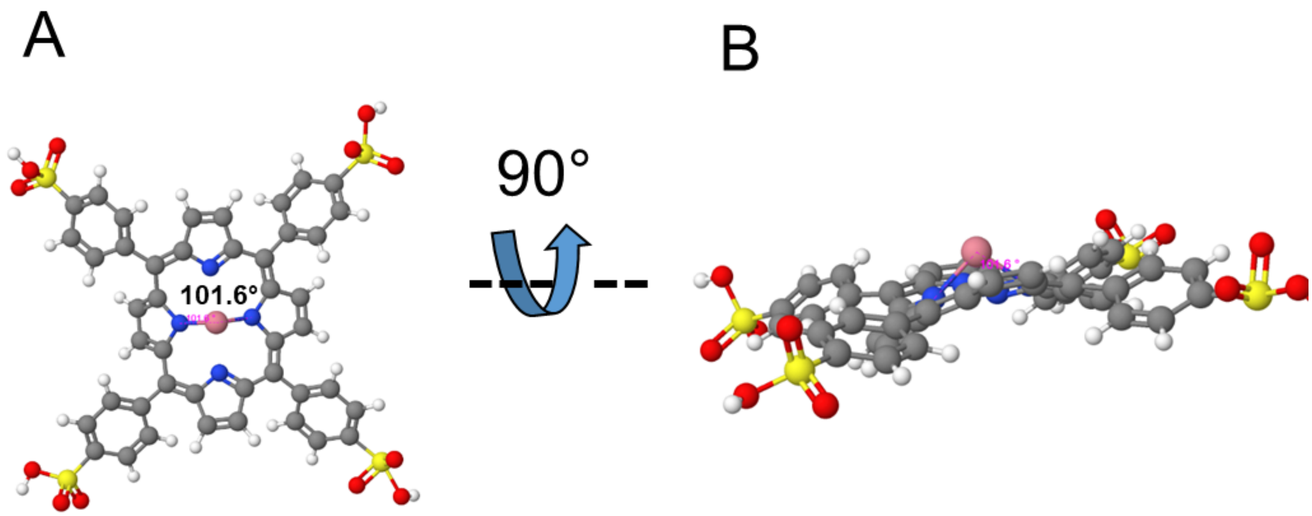

Supplementary Fig. 33: Predicted 3D model of $Co^{2+}$-TPPS complex using energy-minimization simulations in MolView (molview.org). Panels A and B shows the ball and stick representations of the $Co^{2+}$-TPPS at different angles (B is the 90° rotated version of A). The angle between the N-$Co^{2+}$-N in the core was found to be 101.6°.

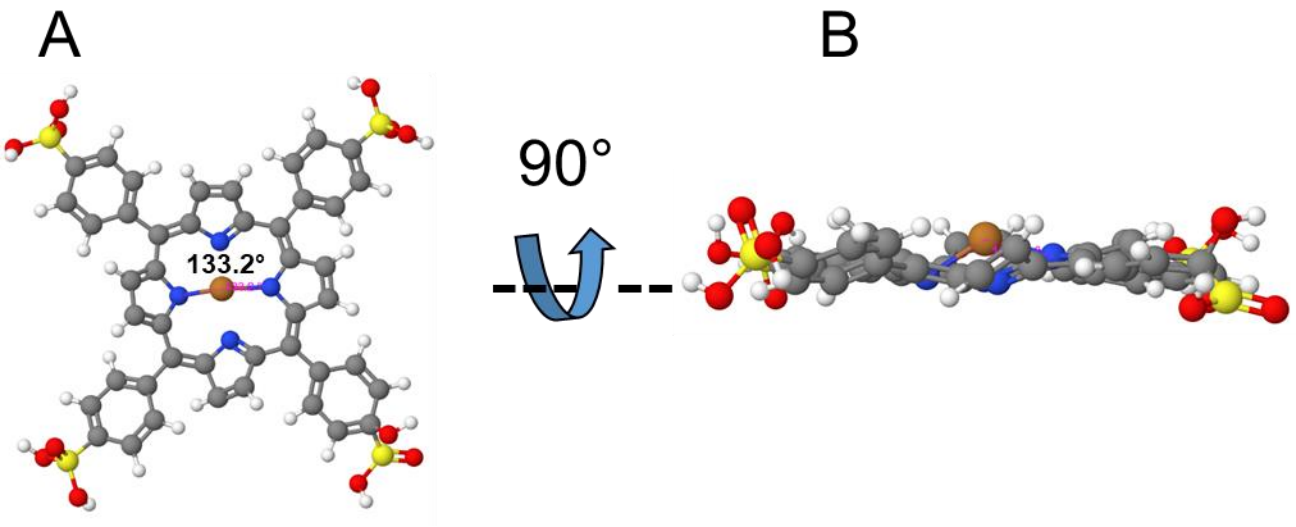

Supplementary Fig. 34: Predicted 3D model of $Cu^{2+}$-TPPS complex using energy-minimization simulations in MolView (molview.org). Panels A and B shows the ball and stick representations of the $Cu^{2+}$-TPPS at different angles (B is the 90° rotated version of A). The angle between the N-$Cu^{2+}$-N in the core was found to be 133.2°.

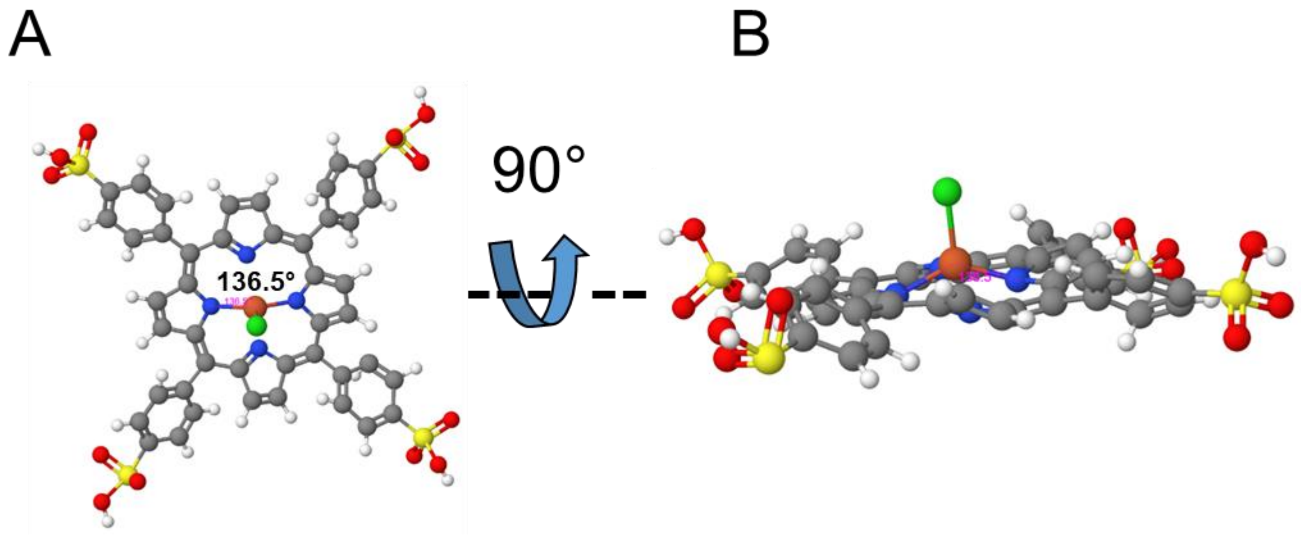

Supplementary Fig. 35: Predicted 3D model of $Fe^{3+}$-TPPS.Cl complex using energy-minimization simulations in MolView (molview.org). Panels A and B shows the ball and stick representations of the of $Fe^{3+}$-TPPS.Cl at different angles (B is the 90° rotated version of A). The angle between the N-$Fe^{3+}$-N in the core was found to be 136.5°.

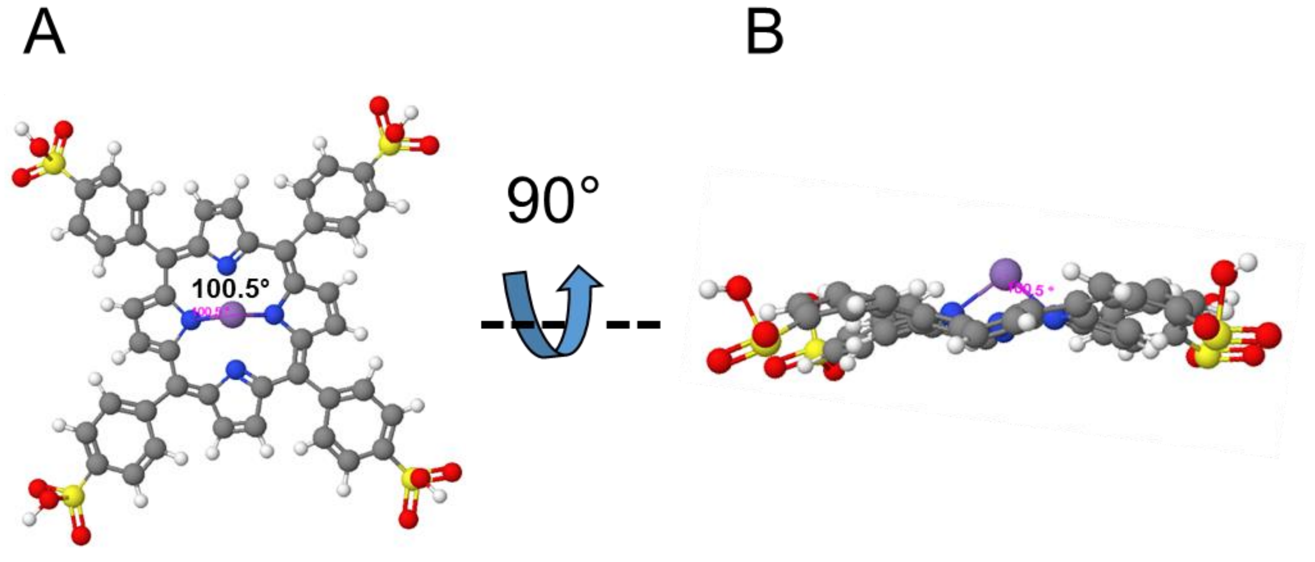

Supplementary Fig. 36: Predicted 3D model of $Mn^{2+}$-TPPS complex using energy-minimization simulations in MolView (molview.org). Panels A and B shows the ball and stick representations of the $Mn^{2+}$-TPPS at different angles (B is the 90° rotated version of A). The angle between the N-$Mn^{2+}$-N in the core was found to be 100.5°.

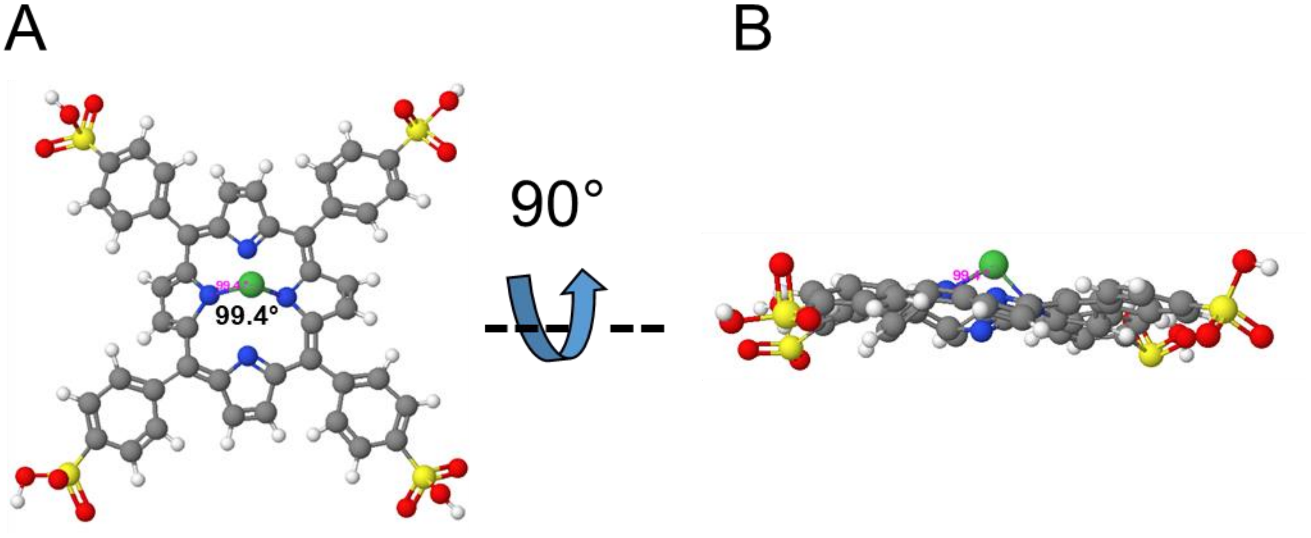

Supplementary Fig. 37: Predicted 3D model of $Ni^{2+}$-TPPS complex using energy-minimization simulations in MolView (molview.org). Panels A and B shows the ball and stick representations of the $Ni^{2+}$-TPPS at different angles (B is the 90° rotated version of A). The angle between the N-$Ni^{2+}$-N in the core was found to be 99.4°.

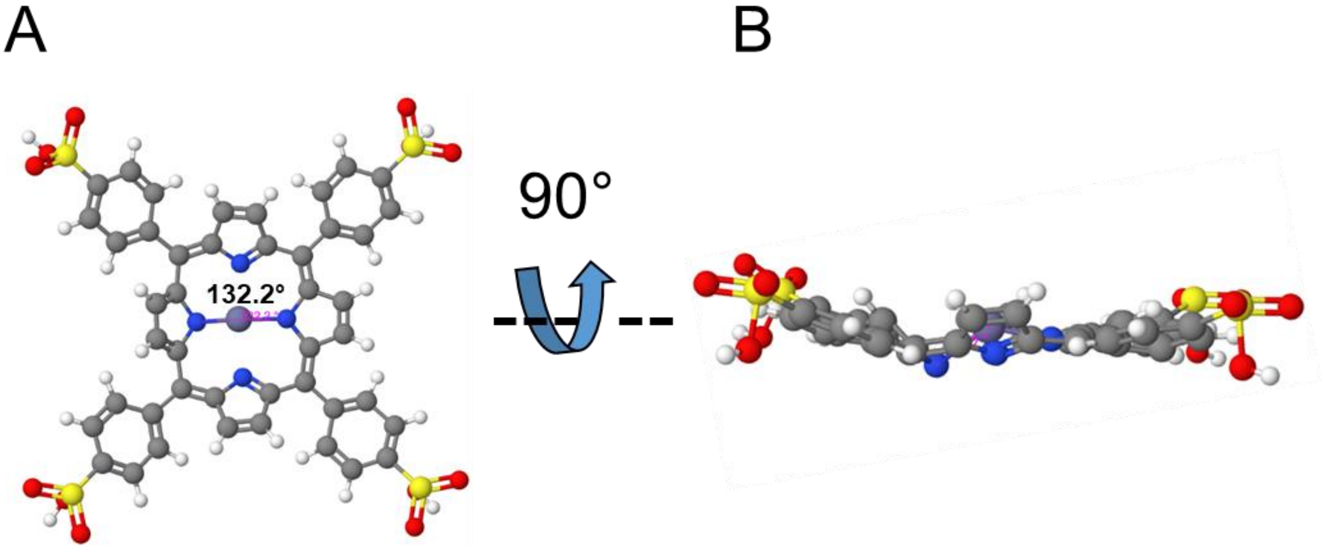

Supplementary Fig. 38: Predicted 3D model of $Zn^{2+}$-TPPS complex using energy-minimization simulations in MolView (molview.org). Panels A and B shows the ball and stick representations of the $Zn^{2+}$-TPPS at different angles (B is the 90° rotated version of A). The angle between the N-$Zn^{2+}$-N in the core was found to be 132.2°.

### Rayleigh scattering at 600 nm to investigate aggregate formation

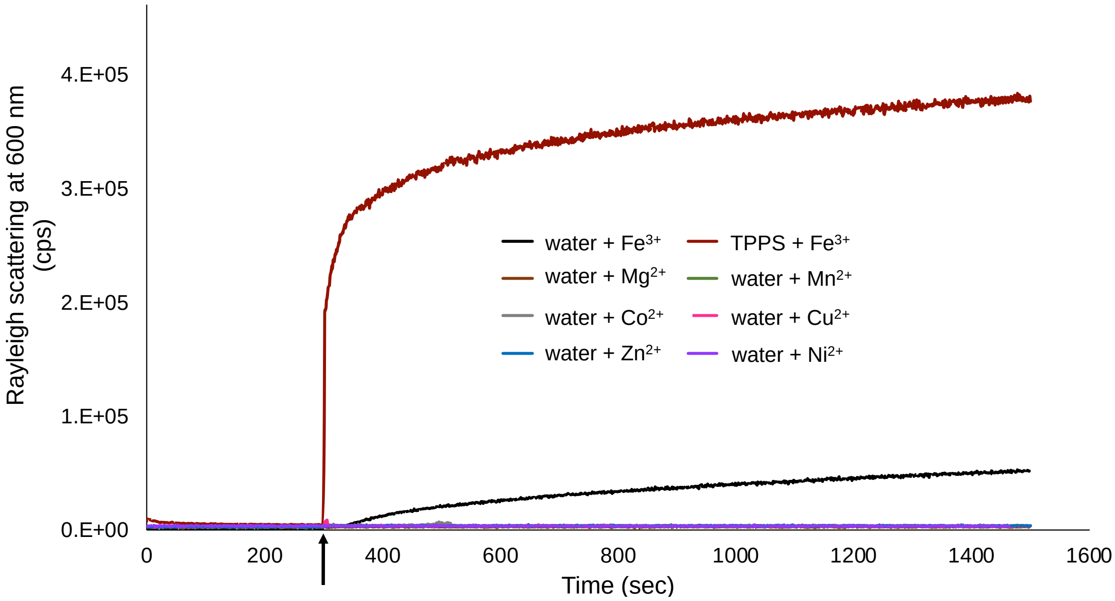

Supplementary Fig. 39: Rayleigh scattering at 600 nm depicting aggregate or any higher order structure (e.g., oxo-hydroxy complexes) formation. Black arrow indicates the addition of metal ions to water (as shown in Figure legend). Brown color shows the addition of $Fe^{3+}$ to TPPS solution for comparison. N=3.

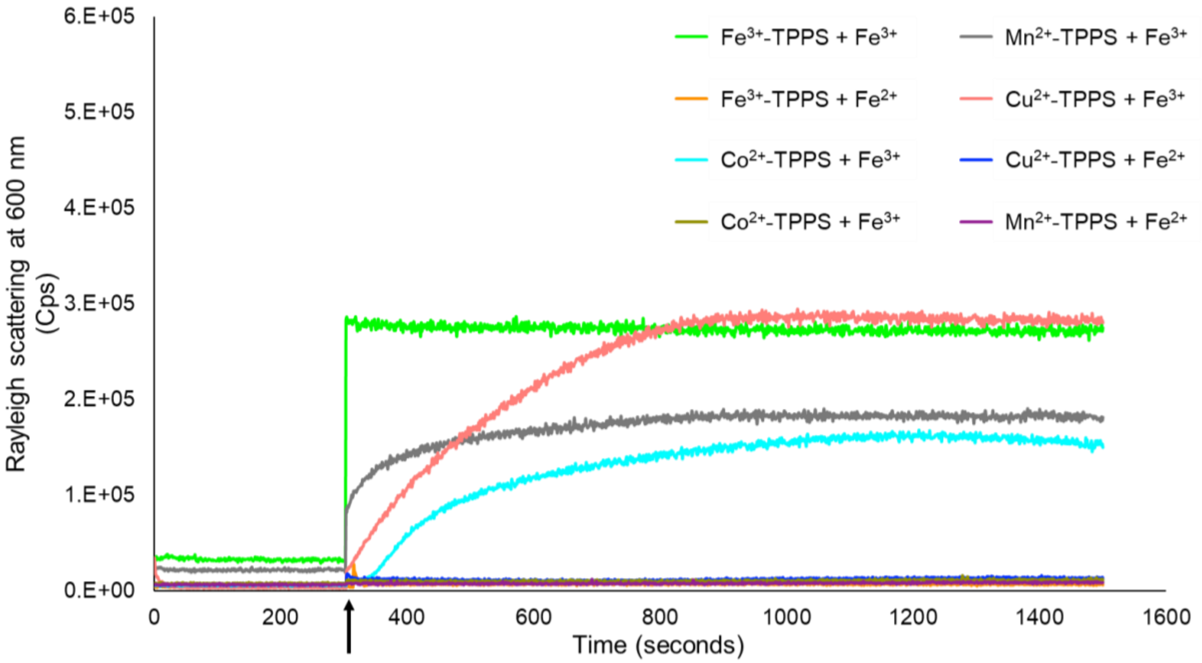

Supplementary Fig. 40: Rayleigh scattering at 600 nm depicting aggregate formation. Black arrow indicates the addition of $Fe^{3+}$ or $Fe^{2+}$ solution in the M-TPPS solution. Different colors show kinetics curve for different M-TPPS (as shown in Figure legend). N=3.

**Differential Interference Contrast (DIC) microscopy characterization of $Fe^{3+}$ and TPPS non-coordinated aggregates (TPPS**$Fe^{3+}$)**

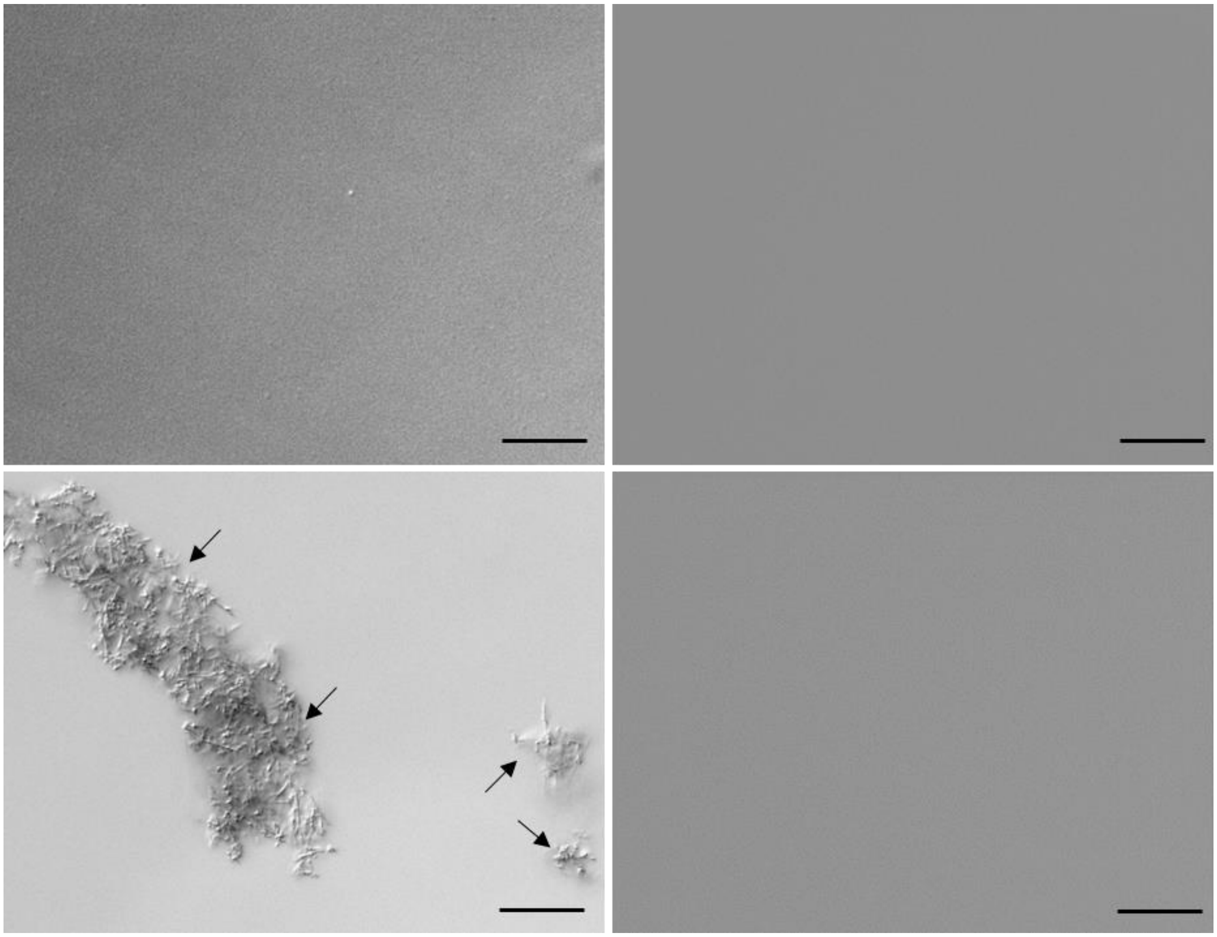

Supplementary Fig. 41: Differential Interference Contrast (DIC) microscopy (under 40X objective). A) 300 µM of $Fe^{3+}$ solution. B) 30 µM TPPS solution. C) co-solute mixture of 300 µM $Fe^{3+}$ and 30 µM TPPS. D) 30 µM $Fe^{3+}$-TPPS solution. Black arrows indicate towards the non-coordinated aggregates. Scale bar = 20 µm.

**Electron Microscopy (EM) characterization of $Fe^{3+}$ and TPPS non-coordinated aggregates (TPPS**$Fe^{3+}$) and its elemental analysis**

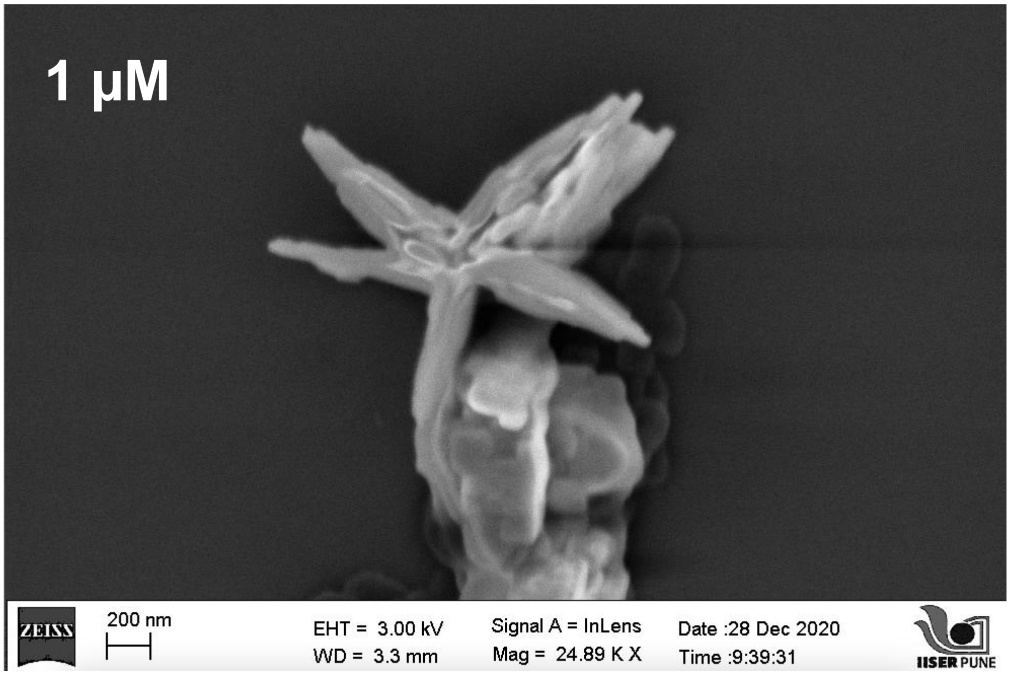

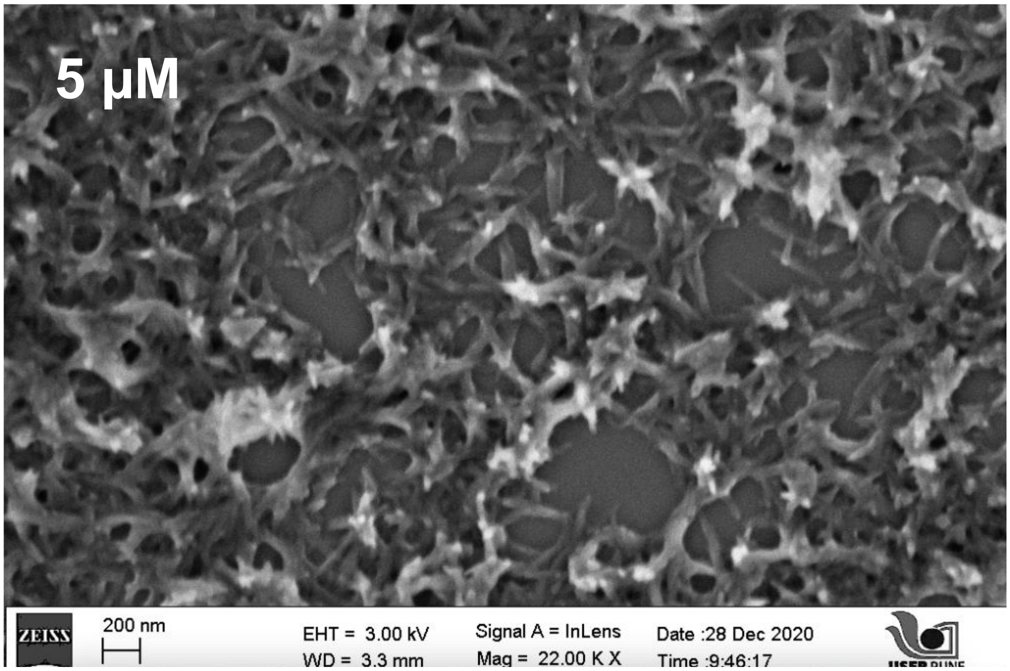

Supplementary Fig. 42: Field Emission Scanning Electron Microscopy (FESEM) images of 1 µM solution of TPPS containing 10 µM of $Fe^{3+}$ ions (A); and 5 µM solution of TPPS containing 50 µM of $Fe^{3+}$ ions.

A

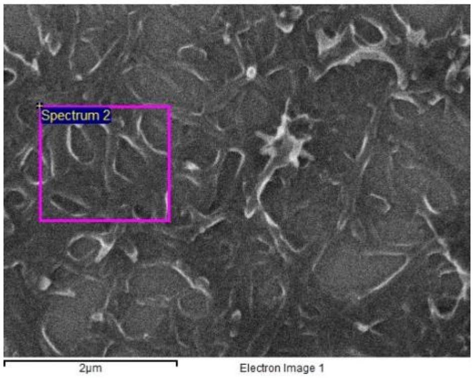

Processing option : All elements analyzed (Normalised)
Number of iterations = 3

| Element | Weight% | Atomic% |
|---|---|---|
| C K | 43.30 | 50.65 |
| N K | 11.59 | 11.62 |
| O K | 41.81 | 36.72 |
| S K | 0.90 | 0.40 |
| Fe K | 2.40 | 0.60 |
| Totals | 100.00 | |

B

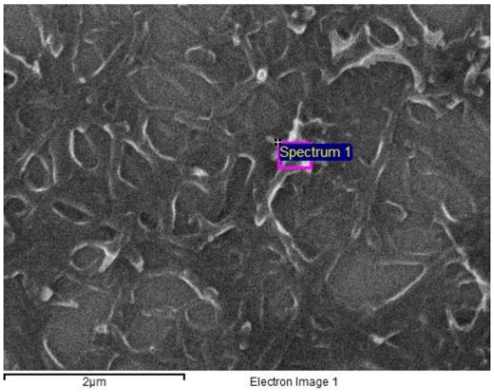

Processing option : All elements analyzed (Normalised)

Number of iterations = 4

| Element | Weight% | Atomic% |
|---|---|---|
| C K | 43.58 | 51.70 |
| N K | 9.99 | 10.16 |
| O K | 40.98 | 36.50 |
| S K | 1.33 | 0.59 |
| Fe K | 4.11 | 1.05 |
| Totals | 100.00 | |

Supplementary Fig. 43: Elemental analysis of two different areas (as shown by lavender boxes in Panel A and B) in the surface of the FESEM images of 5 µM solution of TPPS containing 50 µM of $Fe^{3+}$ ions using Energy Dispersive Spectroscopy (EDS).

**Formation of coordinated $Fe^{3+}$-TPPS complex upon heating the Fe3+ and TPPS co-solute mixture at 100ºC.**

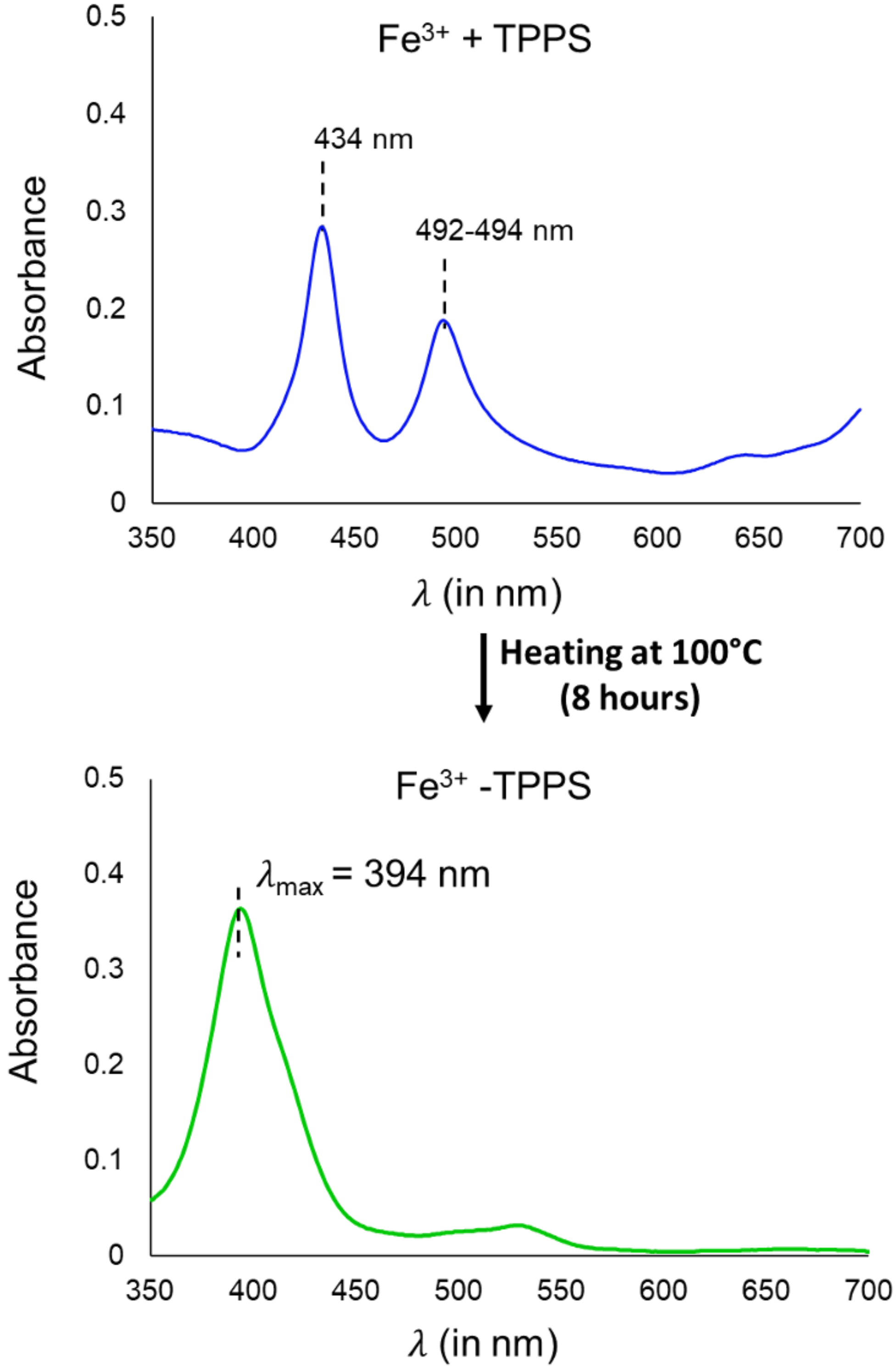

Supplementary Fig. 44: UV spectrum of the preformed $Fe^{3+}$-TPPS coordinated complex with $\lambda_{max}$ at 394 nm (green spectrum, right panel) that results upon heating of the co-solute mixture of 0.3 mM $Fe^{3+}$ and 0.03 mM TPPS (blue spectrum, left panel), at 100ºC for 8 hours.

**Oxidation of NADH monitored by the change in UV absorption**

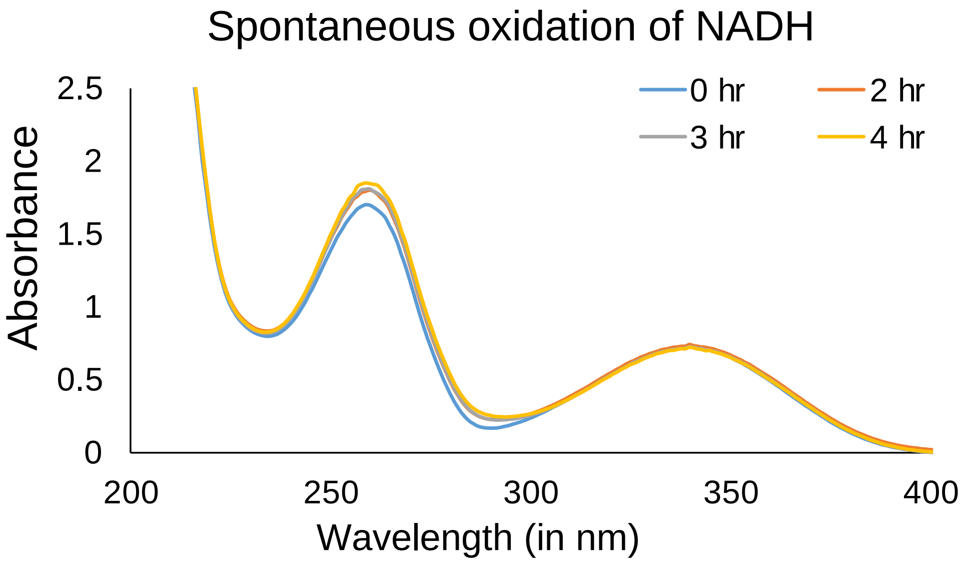

Supplementary Fig. 45: Spontaneous oxidation of NADH. There was no decrease in the characteristic peak of NADH at 340 nm was observed, indicating no oxidation. Different colors show different time points after which the reaction was analyzed; at the initiation of reaction (0 hr), after two hours (2 hr), three hours (3 hr) and four hours (4 hr), respectively. X-axis shows the wavelength and Y-axis shows the absorbance; N=3

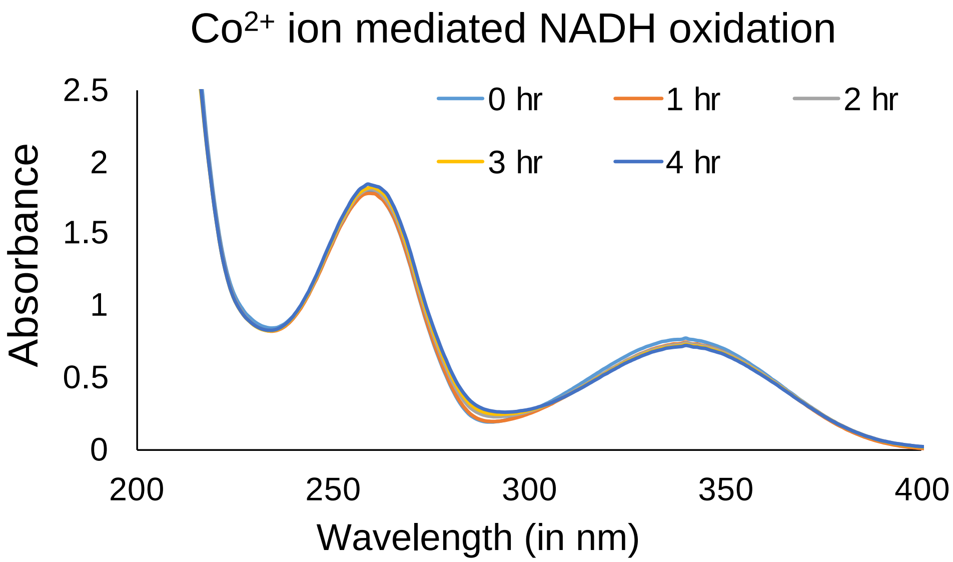

Supplementary Fig. 46: $Co^{2+}$ mediated oxidation of NADH. There was no decrease in the characteristic peak of NADH at 340 nm was observed, indicating no oxidation. Different colors show different time points after which the reaction was analyzed; at the initiation of reaction (0 hr), after one hour (1 hr), two hours (2 hr), three hours (3 hr) and four hours (4 hr), respectively. X-axis shows the wavelength and Y-axis shows the absorbance; N=3

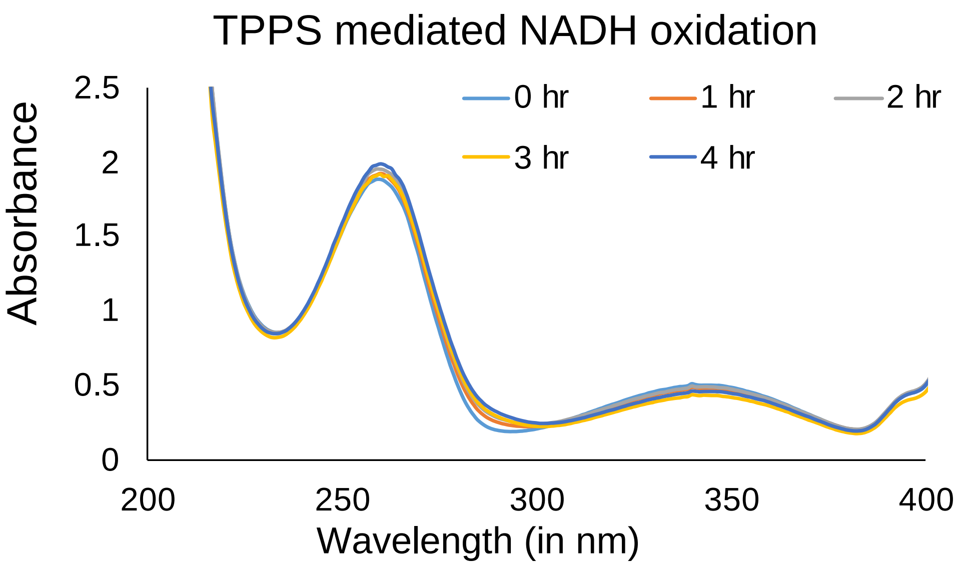

Supplementary Fig. 47: TPPS mediated oxidation of NADH. There was no decrease in the characteristic peak of NADH at 340 nm was observed, indicating no oxidation. Different colors show different time points after which the reaction was analyzed; at the initiation of reaction (0 hr), after one hour (1 hr), two hours (2 hr), three hours (3 hr) and four hours (4 hr), respectively. X-axis shows the wavelength and Y-axis shows the absorbance; N=3

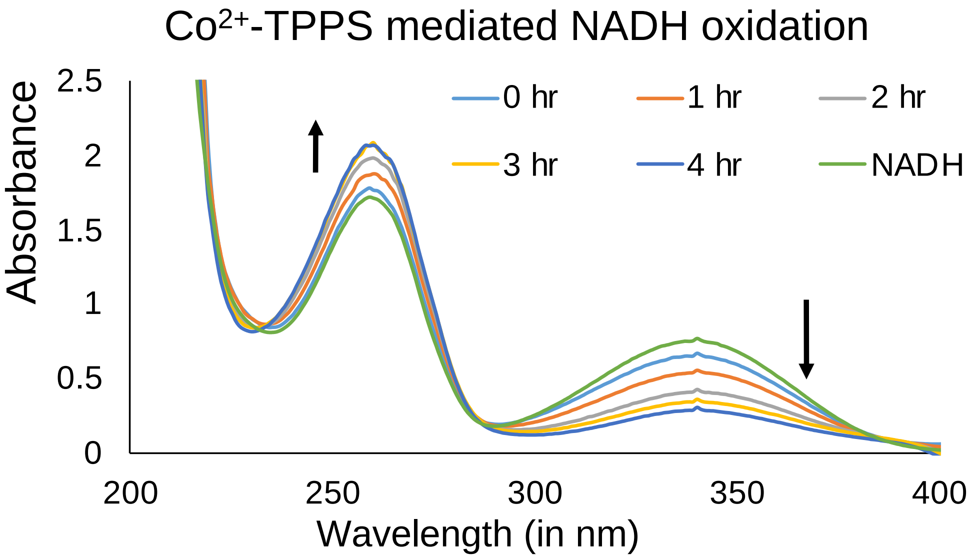

Supplementary Fig. 48: $Co^{2+}$-TPPS mediated oxidation of NADH. The decrease in the characteristic peak of NADH at 340 nm (highlighted by downward black arrow) was monitored. Different colors show different time points after which reaction was analyzed i.e., at the initiation of reaction (0 hr), after one hour (1 hr), two hours (2 hr), three hours (3 hr) and four hours (4 hr). X-axis shows the wavelength and Y-axis shows the absorbance; N=3

**Table S1: Reduction potentials of the investigated metal ions and their respective oxidation state change.**

| Metals | Standard reduction potential (Volts) | Changing oxidation state |
|---|---|---|
| **Benzoquinone** | 0.643[a] | 2 $e^-/2H^+$ |
| | 0.099[a] | 1 $e^-$ |
| **Mg** | -2.356[b] | +2 to 0 |
| **Fe** | +0.771[b] | +3 to +2 |
| | -0.44[b] | +2 to 0 |
| **Co** | -0.277[b] | +2 to 0 |
| **Ni** | -0.257[b] | +2 to 0 |
| **Cu** | 0.3419[b] | +2 to 0 |
| | 0.159[b] | +2 to +1 |
| | 0.520[b] | +1 to 0 |
| **Zn** | -0.762[b] | +2 to 0 |
| **Mn** | -1.170[b] | +2 to 0 |

**[a]: adapted from**
Huynh, M. T. et. al, *JACS.* **138**, 15903–15910 (2016).

**[b]: adapted from**
Bard & AllenJ. *Standard Potentials in Aqueous Solution*. (2017);
Milazzo, G. et. al, *J. Electrochem. Soc.* 125, 261C-261C (1978);
Swift, E. H. & Butler, E. A. *Quantitative measurements and chemical equilibria*. (W. H. Freeman, 1972).

**Table S2: Measured pH of the solution using pH strips with 0.5-unit resolution, at the initiation (0 hour) and at the end (4 hours) of different oxidation reactions.**

| Reactions | pH | |
|---|---|---|
| | 0 hour | 4 hours |
| HQ | 7 | 7 |
| HQ+ $Mg^{2+}$ | 6.5-7 | 6.5-7 |
| HQ+ $Mn^{2+}$ | 7 | 7 |
| HQ+ $Fe^{3+}$ | 5.5 | 5.5 |
| HQ+ $Co^{2+}$ | 7 | 6.5-7 |
| HQ+ $Ni^{2+}$ | 7 | 7 |
| HQ+ $Cu^{2+}$ | 6 | 5.5 |
| HQ+ $Zn^{2+}$ | 7 | 7 |
| HQ+ TPPS | 7 | 6.5 |
| HQ+ TPPS+ $Mg^{2+}$ | 6.5-7 | 6.5-7 |
| HQ+ TPPS+ $Mn^{2+}$ | 7 | 6.5-7 |
| HQ+ TPPS+ $Fe^{3+}$ | 5 | 5 |
| HQ+ TPPS+ $Co^{2+}$ | 7 | 6.5 |
| HQ+ TPPS+ $Ni^{2+}$ | 7 | 7 |
| HQ+ TPPS+ $Cu^{2+}$ | 6.5-7 | 6.5-7 |
| HQ+ TPPS+ $Zn^{2+}$ | 6.5-7 | 6.5-7 |
| HQ+ $Mn^{2+}$-TPPS | 7 | 6-6.5 |
| HQ+ $Fe^{3+}$-TPPS | 6.5-7 | 6.5-7 |
| HQ+ $Co^{2+}$-TPPS | 7 | 6.5 |
| HQ+ $Ni^{2+}$-TPPS | 7 | 7 |
| HQ+ $Cu^{2+}$-TPPS | 5.5-6 | 5-5.5 |
| HQ+ $Zn^{2+}$-TPPS | 7 | 6.5-7 |

**Table S3: Change in the $\lambda_{max}$ in the UV absorbance spectrum of control reactions and co-solute reactions consisting of TPPS and metal ions in (1:10) ratio; at the initiation of the reaction (0 hr $\lambda_{max}$) and after four hours of incubation at 40 ºC (4 hr $\lambda_{max}$), respectively. (Orange color highlights the changes in the value of $\lambda_{max}$).**

| Reaction | 0 hr $\lambda_{max}$ | 4 hr $\lambda_{max}$ |
|---|---|---|
| **TPPS** | 413 | 414 & 434 |
| **TPPS + $Mg^{2+}$** | 414 & 434 | 414 & 434 |
| **TPPS + $Mn^{2+}$** | 414 & 434 | 414 & 434 |
| **TPPS + $Fe^{3+}$** | 432 & 493 | 432 & 493 |
| **TPPS + $Co^{2+}$** | 414 & 434 | 414 & 427 |
| **TPPS + $Ni^{2+}$** | 414 & 434 | 414 & 434 |
| **TPPS + $Cu^{2+}$** | 412 | 412 |
| **TPPS + $Zn^{2+}$** | 414 & 434 | 421 |

**Table S4: High-Resolution Mass spectrometry (HRMS) analysis of formed metal-TPPS complexes.**

| Chemical species | Molecular formula [M] | Species | Calculated Mass | Observed Mass | ppm error |
|---|---|---|---|---|---|
| $Mn^{2+}$-TPPS | $C_{44}H_{28}MnN_4O_{12}S_4$ | $[M-4H]^{4-}$ | 245.7419 | 245.7441 | -8.95 |
| | | $[M-4H]^{3-}$ | 327.9916 | 327.9947 | -9.45 |
| $Fe^{3+}$-TPPS.OH | $C_{44}H_{29}FeN_4O_{13}S_4$ | $[M-4H]^{4-}$ | 250.4926 | 250.4938 | -4.79 |
| $Co^{2+}$-TPPS | $C_{44}H_{28}MnN_4O_{12}S_4$ | $[M-4H]^{4-}$ | 246.7407 | 246.7405 | 0.811 |
| | | $[M-4H]^{3-}$ | 329.3233 | 329.3234 | -0.30 |
| $Ni^{2+}$-TPPS | $C_{44}H_{28}MnN_4O_{12}S_4$ | $[M-4H]^{4-}$ | 246.4912 | 246.4905 | 2.84 |
| $Cu^{2+}$-TPPS | $C_{44}H_{28}MnN_4O_{12}S_4$ | $[M-4H]^{4-}$ | 247.7398 | 247.7386 | 4.84 |
| $Zn^{2+}$-TPPS | $C_{44}H_{28}MnN_4O_{12}S_4$ | $[M-4H]^{4-}$ | 247.9897 | 247.9891 | 2.42 |

**Significance analysis for different reactions**

**Table S5: Comparison of the $Cu^{2+}$ ion-mediated oxidation across varying time periods (2 hrs, 4 hrs, 6 hrs, 8 hrs) and with respect to the initiation of the reaction (0 hr) using two-tailed type 2 t-test for varying atomic ratios of HQ: $Cu^{2+}$ i.e., 1:1, 1:2, 2:1 and 1:4. N = 3.**

| | p-value calculated using two tailed type 2 t-test | | | |
|---|---|---|---|---|
| | $Cu^{2+}$ (1:1) | $Cu^{2+}$ (1:2) | $Cu^{2+}$ (2:1) | $Cu^{2+}$ (1:4) |
| **0 hr vs 2 hrs** | 8.7E-08 | 2.3E-07 | 2.3E-05 | 2.4E-04 |
| **2 hrs vs 4 hrs** | 1.5E-01 | 5.9E-02 | 9.2E-02 | 1.5E-01 |
| **4 hrs vs 6 hrs** | 2.9E-01 | 3.5E-01 | 5.2E-01 | 3.6E-01 |
| **6 hrs vs 8 hrs** | 3.0E-01 | 9.8E-03 | 6.1E-01 | 6.8E-01 |
| **0 hr vs 2 hrs** | 8.7E-08 | 2.3E-07 | 2.3E-05 | 2.4E-04 |
| **0 hr vs 4 hrs** | 1.4E-05 | 2.7E-05 | 7.8E-05 | 3.0E-03 |
| **0 hr vs 6 hrs** | 1.4E-04 | 1.4E-05 | 1.4E-03 | 1.1E-04 |
| **0 hr vs 8 hrs** | 1.7E-04 | 3.4E-08 | 7.2E-06 | 2.3E-02 |

**Table S6: Comparison of the $Cu^{2+}$ ion-mediated oxidation after varying time periods (2 hrs, 4 hrs, 6 hrs, 8 hrs) with respect to different ratios of HQ: $Cu^{2+}$ i.e., control (no Cu added), 1:1, 1:2, 2:1 and 1:4. using two-tailed type 2 t-test. N = 3.**

| | **p-value calculated using two tailed type 2 t-test** | | | |
|---|---|---|---|---|
| | **2 hrs** | **4 hrs** | **6 hrs** | **8 hrs** |
| **Control vs $Cu^{2+}$ (1:1)** | 4.4E-04 | 5.5E-03 | 1.2E-02 | 1.8E-02 |
| **Control vs $Cu^{2+}$ (1:2)** | 3.3E-04 | 4.9E-03 | 5.4E-03 | 1.1E-04 |
| **Control vs $Cu^{2+}$ (2:1)** | 1.4E-03 | 4.7E-03 | 2.5E-02 | 6.7E-04 |
| **Control vs $Cu^{2+}$ (1:4)** | 2.4E-04 | 3.0E-03 | 1.1E-04 | 2.3E-02 |
| **$Cu^{2+}$ (1:1) vs $Cu^{2+}$ (1:2)** | 8.4E-01 | 4.6E-01 | 3.9E-01 | 6.5E-02 |
| **$Cu^{2+}$ (1:1) vs $Cu^{2+}$ (2:1)** | 1.1E-01 | 3.8E-01 | 3.7E-01 | 4.9E-01 |
| **$Cu^{2+}$ (1:1) vs $Cu^{2+}$ (1:4)** | 6.1E-03 | 1.4E-01 | 3.3E-02 | 4.1E-01 |
| **$Cu^{2+}$ (1:2) vs $Cu^{2+}$ (2:1)** | 7.9E-02 | 1.7E-01 | 8.2E-01 | 1.4E-03 |
| **$Cu^{2+}$ (1:2) vs $Cu^{2+}$ (1:4)** | 1.1E-02 | 5.0E-01 | 4.8E-02 | 4.6E-01 |
| **$Cu^{2+}$ (2:1) vs $Cu^{2+}$ (1:4)** | 4.8E-03 | 3.9E-02 | 2.2E-01 | 7.3E-02 |

**Table S7: Comparison of the different metal ion-mediated oxidation when atomic ratio of HQ: corresponding metal ion is 1:2, after varying time periods (2 hrs, 4 hrs, 6 hrs, 8 hrs) with oxidation at the initiation of the reactions (0 hr), using two-tailed type 2 t-test. N = 3.**

| | **1:2** | | | | | |
|---|---|---|---|---|---|---|
| | **p-value calculated using two tailed type 2 t-test** | | | | | |
| | **HQ** | **$Mg^{2+}$** | **$Mn^{2+}$** | **$Fe^{3+}$** | **$Co^{2+}$** | **$Ni^{2+}$** |
| **0 hr vs 2 hrs** | 8.3E-01 | 8.2E-02 | 7.7E-01 | 6.6E-01 | 5.3E-01 | 1.8E-02 |
| **0 hr vs 4 hrs** | 6.9E-01 | 6.3E-06 | 6.2E-01 | 2.5E-01 | 7.1E-01 | 1.5E-01 |
| **0 hr vs 6 hrs** | 3.7E-01 | 9.0E-01 | 6.7E-01 | 1.7E-01 | 6.8E-01 | 5.9E-01 |
| **0 hr vs 8 hrs** | 3.5E-01 | 8.4E-03 | 3.2E-01 | 3.6E-01 | 3.7E-01 | 4.4E-01 |

**Table S8: Comparison of the different metal ion-mediated oxidation when atomic ratio of HQ: corresponding metal ion is 1:2, across varying time periods (2 hrs, 4 hrs, 6 hrs, 8 hrs) using two-tailed type 2 t-test. N = 3.**

| | **1:2** | | | | | |
|---|---|---|---|---|---|---|
| | **p-value calculated using two tailed type 2 t-test** | | | | | |
| | **HQ** | **$Mg^{2+}$** | **$Mn^{2+}$** | **$Fe^{3+}$** | **$Co^{2+}$** | **$Ni^{2+}$** |
| **2 hrs vs 4 hrs** | 6.5E-01 | 1.1E-01 | 8.6E-01 | 4.0E-01 | 8.8E-01 | 2.0E-01 |
| **4 hrs vs 6 hrs** | 4.8E-01 | 7.3E-01 | 8.0E-01 | 9.0E-01 | 9.6E-01 | 5.0E-01 |
| **6 hrs vs 8 hrs** | 7.6E-01 | 3.7E-01 | 4.6E-01 | 1.5E-01 | 4.4E-01 | 4.1E-01 |

**Table S9: Comparison of the metal ion mediated oxidation when atomic ratio of HQ: corresponding metal ion is 1:1, after 4 hrs with respect to the oxidation at the initiation of the reaction i.e., 0 hr using two-tailed type 2 t-test i.e., N = 3.**

| | **1:1** | | | | | | |
|---|---|---|---|---|---|---|---|
| | **p-value calculated using two tailed type 2 t-test** | | | | | | |
| | **$Mn^{2+}$** | **$Fe^{2+}$** | **$Fe^{3+}$** | **$Co^{2+}$** | **$Ni^{2+}$** | **$Cu^{2+}$** | **$Zn^{2+}$** |
| **0 hr vs 4 hrs** | 3.4E-03 | 3.9E-03 | 1.6E-02 | 4.6E-01 | 4.9E-01 | 1.9E-05 | 8.2E-02 |

**Table S10: Comparison of the oxidation in TPPS and its co-solute reactions with various metal ions i.e., $Mg^{2+}$, $Mn^{2+}$, $Fe^{2+}$, $Fe^{3+}$, $Co^{2+}$, $Ni^{2+}$, $Cu^{2+}$ and $Zn^{2+}$, after 4 hrs with respect to the oxidation at the initiation of the corresponding reaction (0 hr), using two-tailed type 2 t-test. N = 3.**

| | **1:1** |
|---|---|
| | **p-value calculated using two tailed type 2 t-test** |
| | **0 hr vs 4 hrs** |
| **TPPS** | 3.9E-03 |
| **TPPS + $Mg^{2+}$** | 8.7E-01 |
| **TPPS + $Mn^{2+}$** | 2.0E-01 |
| **TPPS + $Fe^{2+}$** | 2.6E-03 |
| **TPPS + $Fe^{3+}$** | 5.1E-01 |
| **TPPS + $Co^{2+}$** | 2.9E-02 |
| **TPPS + $Ni^{2+}$** | 1.1E-02 |
| **TPPS + $Cu^{2+}$** | 1.4E-02 |
| **TPPS + $Zn^{2+}$** | 8.2E-03 |

**Table S11: Comparison of the oxidation across TPPS and its various co-solute reactions with different metal ions i.e., $Mg^{2+}$, $Mn^{2+}$, $Fe^{2+}$, $Fe^{3+}$, $Co^{2+}$, $Ni^{2+}$, $Cu^{2+}$ and $Zn^{2+}$, after 4 hrs using two-tailed type 2 t-test. N = 3.**

| | **1:1** |
|---|---|
| | **p-value calculated using two tailed type 2 t-test** |
| | **4 hrs** |
| **TPPS vs TPPS + $Mg^{2+}$** | 3.1E-01 |
| **TPPS vs TPPS + $Mn^{2+}$** | 1.4E-01 |
| **TPPS vs TPPS + $Fe^{2+}$** | 2.6E-02 |
| **TPPS vs TPPS + $Fe^{3+}$** | 8.0E-05 |
| **TPPS vs TPPS +$Co^{2+}$** | 3.4E-02 |
| **TPPS vs TPPS + $Ni^{2+}$** | 1.4E-02 |
| **TPPS vs TPPS +$Cu^{2+}$** | 2.4E-01 |
| **TPPS vs TPPS + $Zn^{2+}$** | 2.5E-01 |

**Table S12: Comparison of the oxidation across different metal ions i.e., $Mn^{2+}$, $Fe^{2+}$, $Fe^{3+}$, $Co^{2+}$, $Ni^{2+}$, $Cu^{2+}$ and $Zn^{2+}$ and their co-solute reactions with TPPS, after 4 hrs using two-tailed type 2 t-test. N = 3.**

| | **1:1** |
|---|---|
| | **p-value calculated using two tailed type 2 t-test** |
| | **4 hrs** |
| **$Mn^{2+}$ vs TPPS + $Mn^{2+}$** | 6.1E-03 |
| **$Fe^{2+}$ vs TPPS + $Fe^{2+}$** | 9.9E-01 |
| **$Fe^{3+}$ vs TPPS + $Fe^{3+}$** | 4.0E-04 |
| **$Co^{2+}$ vs TPPS +$Co^{2+}$** | 3.8E-02 |
| **$Ni^{2+}$ vs TPPS + $Ni^{2+}$** | 4.5E-01 |
| **$Cu^{2+}$ vs TPPS +$Cu^{2+}$** | 8.3E-04 |
| **$Zn^{2+}$ vs TPPS + $Zn^{2+}$** | 6.5E-02 |

**Table S13: Comparison of the oxidation in coordinated $Fe^{3+}$-TPPS complex mediated oxidation with respect to the free $Fe^{3+}$ ion mediated oxidation and its co-solute reaction with TPPS at the initiation of the reaction i.e., 0 hr and after 4 hrs using two-tailed type 2 t-test. N = 3.**

| **1:1** | | |
|---|---|---|
| **p-value calculated using two tailed type 2 t-test** | | |
| | **0 hr** | **4 hrs** |
| **$Fe^{3+}$ vs $Fe^{3+}$-TPPS** | 7.8E-04 | 1.4E-04 |
| **$Fe^{3+}$ + TPPS vs $Fe^{3+}$-TPPS** | 6.8E-04 | 4.0E-03 |

**Table S14: Comparison of the oxidation in coordinated M-TPPS complex mediated oxidation with respect to the corresponding free metal ion and their co-solute reaction with TPPS for different metal ions i.e., $Fe^{3+}$, $Cu^{2+}$, $Zn^{2+}$, $Co^{2+}$, $Ni^{2+}$ and $Mn^{2+}$, at the initiation of the reaction i.e., 0 hr and after 4 hrs using two-tailed type 2 t-test. N = 3.**

| 1:1 | |
|---|---|
| p-value calculated using two tailed type 2 t-test | |
| | **4 hrs** |
| **$Fe^{3+}$ vs $Fe^{3+}$-TPPS** | 1.4E-04 |
| **$Fe^{3+}$ + TPPS vs $Fe^{3+}$-TPPS** | 4.0E-03 |
| **$Cu^{2+}$ vs $Cu^{2+}$-TPPS** | 5.8E-02 |
| **$Cu^{2+}$ + TPPS vs $Cu^{2+}$-TPPS** | 4.4E-02 |
| **$Zn^{2+}$ vs $Zn^{2+}$-TPPS** | 4.6E-02 |
| **$Zn^{2+}$ + TPPS vs $Zn^{2+}$-TPPS** | 6.8E-01 |
| **$Co^{2+}$ vs $Co^{2+}$-TPPS** | 9.9E-05 |
| **$Co^{2+}$ + TPPS vs $Co^{2+}$-TPPS** | 5.8E-04 |
| **$Ni^{2+}$ vs $Ni^{2+}$-TPPS** | 1.7E-01 |
| **$Ni^{2+}$ + TPPS vs $Ni^{2+}$-TPPS** | 1.6E-01 |
| **$Mn^{2+}$ vs $Mn^{2+}$-TPPS** | 4.0E-01 |
| **$Mn^{2+}$ + TPPS vs $Mn^{2+}$-TPPS** | 7.0E-01 |
